# Challenging selective vulnerability in Parkinson’s disease: a systematic review and meta-analysis

**DOI:** 10.64898/2026.05.13.724902

**Authors:** Will Lunt, Jay A. Moore, Emilie Cottard, Alan E. Murphy, Maitri Shah, Ji Sang, Jethro Choi, Hiranyamaya Dash, Sarah Dawson, Nathan Green, Elina Nagaeva, Samuel Burke, Julian P. T. Higgins, Nathan G. Skene

**Affiliations:** UK DRI at Imperial College London, Department of Brain Sciences, London, UK; Imperial College London, London, UK; Bristol Medical School (Population Health Sciences), University of Bristol, Bristol, UK; Universite de Montreal, Montreal, QC, Canada; Faculty of Mathematical & Physical Sciences, University College London, London, UK

## Abstract

Selective vulnerability is widely assumed in Parkinson’s disease (PD), but whether dopaminergic neurons of the substantia nigra are uniquely vulnerable has not been established. Explanations of neuronal degeneration in Parkinson’s disease often centre on the distinctive properties of substantia nigra dopaminergic neurons. Whether these properties are necessary for substantial neuronal loss remains unclear. We preregistered a systematic review of 166 post-mortem case-control studies published between 1963 and 2025 and synthesised neuronal counts and densities from 152 studies using a multilevel meta-analysis. After six decades, only 18% of countable brain atlas labels had been examined, and most populations were represented by a single study. Substantial loss beyond dopaminergic and classically pigmented populations shows that neither dopaminergic identity nor neuromelanin is necessary for marked degeneration, challenging key tenets of selective vulnerability in PD. Comparisons across other anatomical and physiological features remain limited by sparse sampling and uncertainty. We identify less-studied populations with substantial estimated loss and estimate the additional sampling needed to improve precision under specified assumptions. These findings direct replication and comparative counting towards uncertainties that limit explanations of neuronal loss across affected and potentially spared populations.

## Introduction

The identification of nigral neuronal loss, the discovery of striatal dopamine depletion and the success of levodopa established the nigrostriatal system at the centre of Parkinson’s disease (PD) research (Tretiakoff 1919; Parent and Parent 2010; Przedborski 2017). Degeneration of dopaminergic (DA) neurons in the substantia nigra pars compacta (SNc) remains a defining pathological feature (Hughes et al. 1993). However, sleep disturbance, depression, constipation and hyposmia can precede motor onset by years or decades, extending the clinical picture beyond nigrostriatal dysfunction (Braak et al. 2003; Dauer and Przedborski 2003). Explaining PD therefore requires a broader account of which neuronal populations degenerate and what makes them susceptible.

Post-mortem studies have documented loss of noradrenergic (NA) neurons in the locus coeruleus (LC), cholinergic (Ch) neurons in the basal nucleus (BN) and serotonergic (5-HT) neurons in the raphe nuclei (Chan-Palay and Asan 1989; Chui et al. 1986; Dauer and Przedborski 2003; Huynh et al. 2021; Whitehouse et al. 1983). Loss also varies within the DA system: ventral tegmental area (VTA) neurons are relatively preserved, whereas ventrolateral SNc populations are more severely affected (Uhl et al. 1985; Hirsch et al. 1988; Gibb and Lees 1988; Fearnley and Lees 1991). This uneven loss is commonly described as selective vulnerability (Surmeier et al. 2017). Explaining the pattern requires reliable estimates of loss across populations and direct comparisons between them.

Proposed mechanisms of PD include interacting defects in mitochondrial function, protein handling and the cellular environment. Identifying these processes, however, does not explain why their effects differ across neuronal populations. Dopamine metabolism and neuromelanin have both been implicated in vulnerability (Masato et al. 2019; Filimontseva et al. 2025; Surmeier et al. 2017). Quantitative comparisons within and across neuronal classes can test how far these explanations extend, including whether substantial loss occurs in populations without the proposed feature. Such comparisons require estimates from both affected and potentially spared populations.

Neuronal counts have accumulated over decades but remain dispersed across studies. Giguère et al. (2018) compiled reported losses in a narrative review, while Zarow et al. (2003) compared several nuclei within one cohort using matched methods. Neither pooled estimates across the wider literature. Quantitative synthesis can estimate the extent of loss across populations while accounting for uncertainty and differences between studies (Higgins et al. 2009).

We preregistered a systematic review and used Bayesian multilevel meta-analysis to estimate neuronal loss across eligible populations defined by region and neuron type. We examined the distribution of loss and the limits it places on proposed explanations of selective vulnerability. We also identified gaps in anatomical coverage and populations with substantial estimated loss that require independent replication or more precise measurement.

## Methods

### Review design and eligibility

We preregistered the review with PROSPERO on 13 July 2021 (CRD42021265515). The protocol comparison documents subsequent changes, including narrower disease eligibility and revised statistical analysis (Supplementary Table 1). Reporting follows PRISMA 2020 guidance (Page et al. 2021), with outstanding reporting items identified in Supplementary Table 2.

We included human post-mortem case-control studies reporting neuronal counts or densities in idiopathic Parkinson’s disease, with or without dementia, and controls without known neurodegenerative disease. We excluded separately classified dementia with Lewy bodies, atypical or secondary parkinsonism, primary Alzheimer’s disease, familial cases, single-case reports, animal studies, reviews and conference abstracts. Coexisting pathology within otherwise eligible PD cohorts was not uniformly excluded. Meta-analysis required comparable quantitative PD and control measurements; otherwise eligible studies without a usable comparison remained in the review.

We reassessed eligibility during extraction. Study and subgroup exclusions, including how source diagnoses were mapped to eligibility categories, are documented in Supplementary Table 3, Table 4 and Table 5. We recorded age matching, blinding and neuropathological confirmation where reported.

### Search and study selection

We searched MEDLINE, EMBASE, BIOSIS and Web of Science without publication-year restrictions and updated the searches in 2025 (Supplementary Table 6). During search development, we checked whether the searches retrieved studies identified by Giguère et al. (2018). We prioritised English-language publications, but translated and included a Japanese-language study (Ohno et al. 1991). We did not search preprints, grey literature or theses.

After deduplication across databases, four reviewers (WL, MS, JC and JAM) screened titles and abstracts in Rayyan. At least two reviewers assessed each record independently. Disagreements were resolved by discussion, with SB consulted where needed. Five reviewers (WL, MS, JC, JAM and SB) participated in full-text assessment. The PRISMA flow diagram and supplementary study tables document selection and exclusions.

### Data extraction and risk of bias

Five reviewers (WL, MS, JC, JS and JAM) extracted study characteristics, diagnostic groups, brain regions, neuron types, markers, counting methods, group means, sample sizes and measures of variability into a shared database. We also recorded age, disease characteristics, age matching and blinding where available. We digitised values reported only in figures and calculated group summaries from individual observations where available. Disagreements were resolved by checking the source and discussing the extraction. Supplementary Table 7 describes the modelled study-population comparisons; Supplementary Table 8 summarises counting methods, blinding and age matching across included studies.

We assessed risk of bias using an adapted eight-item Joanna Briggs Institute checklist (Joanna Briggs Institute 2020). Supplementary Table 9 provides the adapted questions and aggregate assessments, and Supplementary Methods describes the scoring rules. Scores counted affirmative responses, so higher scores indicated that more criteria were met. Source verification led to revisions of selected assessments, rather than a new independent assessment by two reviewers.

### Data preparation and population definitions

We standardised brain-region, neuron-type and marker labels using curated mappings (Supplementary Table 10; Supplementary Table 5). Where the neuron type was unreported, we inferred a broader class from the region and marker where possible and recorded the inference. We combined SNc and unspecified substantia nigra (SN) DA measurements, and paranigral and VTA DA measurements. We excluded whole-midbrain measurements combining several DA nuclei, nigral m-calpain upregulation measurements and two non-neuronal CA1 populations (Supplementary Table 5; Supplementary Methods: Anatomical harmonisation).

When a study provided several eligible measurements for the same population, we selected one by counting method, then marker, then sample size. Stereology took priority over computer-assisted and manual counting; preferred neuronal markers took priority over surrogate stains (for example, TH over H&E for DA neurons); and larger combined sample sizes broke remaining ties. This reduced 286 eligible measurements to 265 study-population estimates, removing 21 duplicate measurements from 20 study-population combinations. Full selection rules are given in Supplementary Methods: Selection of measurements.

We derived standard deviations from reported measures of variability where possible. For missing values, we used the observed relationship between standard deviation and mean, drawing first on the same paper, then the same population, and finally the wider dataset. Supplementary Methods: Missing variability and group summaries describes the imputation rules and their application.

We mapped anatomical labels to the Allen Brain Atlas ontology (Hawrylycz et al. 2012). Deduplication and exclusion of ventricular, sulcal, subarachnoid and white-matter labels left 597 entries in the coverage denominator. Because parent and child labels can overlap anatomically, coverage refers to atlas labels rather than brain volume. Supplementary Table 10 reports the mappings from source labels to atlas entries.

A shared dictionary standardises region, neuron-class and marker names across the text, figures and analytical tables (Supplementary Table 11). Abbreviations are defined at first use; + and − denote marker-positive and marker-negative labels. The anatomical mapping table retains source labels and atlas identifiers verbatim.

### Descriptive coverage and study accumulation

For Figure 1, we counted distinct publications within each curated region, anatomical group and recorded cell class or marker category. The anatomical overview groups 88 curated regions into broader divisions using an explicit mapping; these groups differ from the countable atlas labels used to calculate coverage in Supplementary Figure 1. Regional totals include all recorded cell categories, whereas population totals refer to a particular region–cell combination. Transmitter and neuropeptide categories are displayed individually; the remaining single-study categories are combined into marker/enzyme, glial and other-annotation groups. Each grouped bar counts distinct publications, so a study represented in several constituent categories is counted only once. A publication can nevertheless contribute to several anatomical or cellular groups. Cumulative trajectories count each publication once per population from its publication year through 2025; separate publications need not represent independent donor cohorts. The anatomical panel displays study counts in uniform badges. Smaller labels identify the two most frequently studied cell categories in each broad anatomical group and the most frequently studied category in each selected nucleus, with ties resolved alphabetically. Category counts can overlap within a study and therefore need not sum to the regional total.

**Figure 1:**
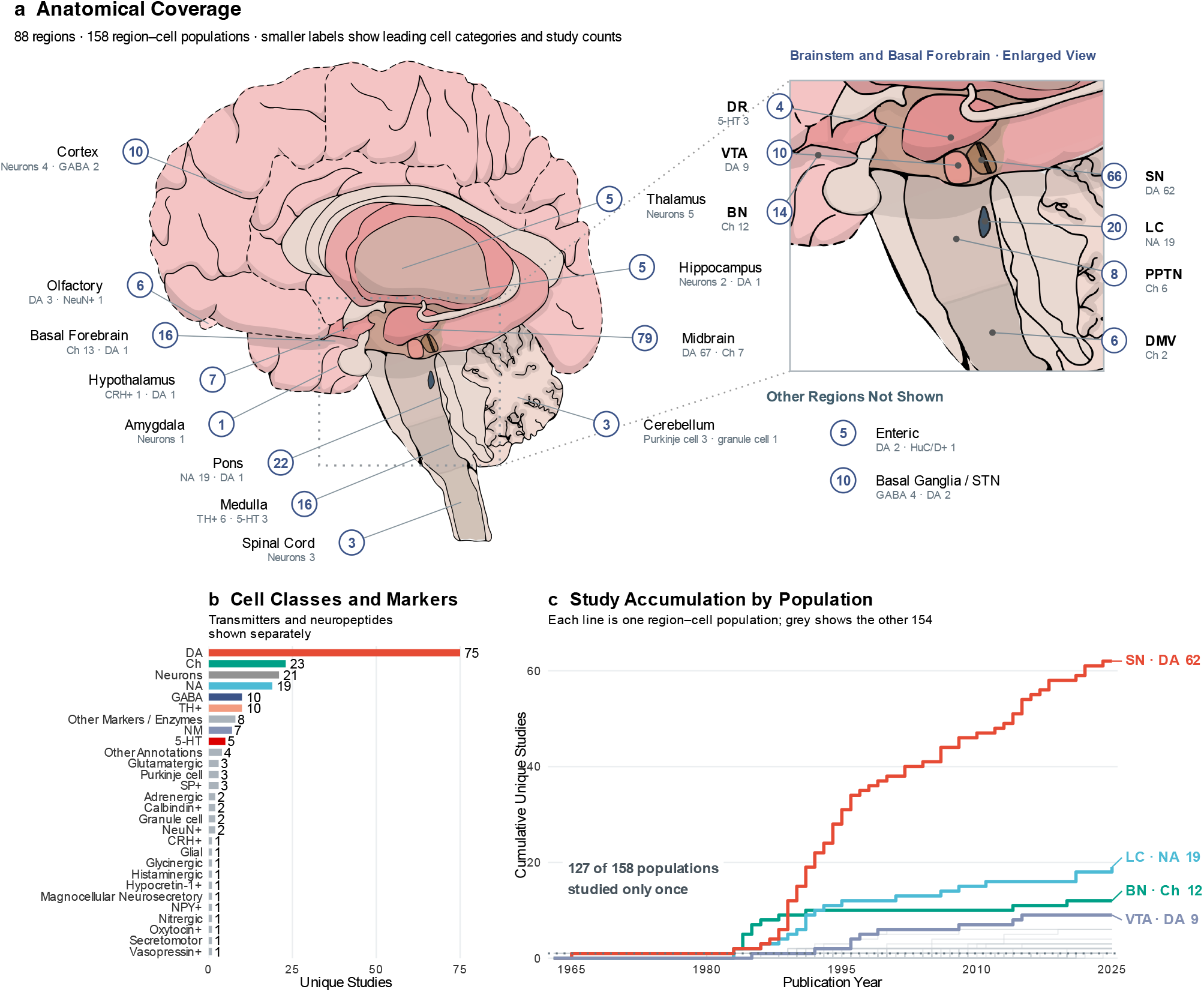
Anatomical and cellular concentration of the PD neuropathology evidence base. (a) Distinct studies by broad anatomical group (left) and selected nucleus (right), displayed in numbered badges. The dotted outline and grey connectors identify the brainstem and basal-forebrain region enlarged at right. Smaller labels show study counts for up to two leading cell categories per broad group and one per selected nucleus, with ties resolved alphabetically. Positions are schematic, including projected lateral structures. Groups summarise 88 curated regions; a study can contribute to several groups. (b) Distinct studies by recorded cell class or marker category, in descending order. Transmitter and neuropeptide categories are shown individually. Remaining single-study marker/enzyme, glial and other annotations are grouped, with each study counted once per bar. Categories can overlap within a publication. (c) Cumulative distinct studies for all 158 region–cell populations, highlighting SN DA, LC NA, BN Ch and VTA DA. Each publication contributes at most once per population. The dotted line marks one study, the final count for 127 populations. Separate publications need not represent independent donor cohorts. SN, substantia nigra; LC, locus coeruleus; BN, basal nucleus of Meynert; VTA, ventral tegmental area; DR, dorsal raphe nucleus; PPTN, pedunculopontine tegmental nucleus; DMV, dorsal motor nucleus of the vagus; STN, subthalamic nucleus; DA, dopaminergic; Ch, cholinergic; NA, noradrenergic; GABA, gamma-aminobutyric acid; TH+, tyrosine hydroxylase-positive; NM, neuromelanin-pigmented; 5-HT, serotonergic; SP+, substance P-positive; NeuN+, neuronal nuclear antigen-positive; CRH+, corticotropin-releasing hormone-positive; NPY+, neuropeptide Y-positive; HuC/D+, HuC/HuD-positive. Full population terminology is provided in Supplementary Table 11.

### Statistical analysis

We used R 4.6.1 (R Core Team 2024), metafor 5.0.1 for effect-size calculations (Viechtbauer 2010), and brms 2.23.0 with Stan for Bayesian meta-analysis (Bürkner 2017).

For each study and population, we calculated the ratio of mean neuronal count or density in PD to that in controls. We analysed the log ratio and its estimated sampling variance, using the small-sample correction implemented in metafor. Results are expressed as percentage loss:

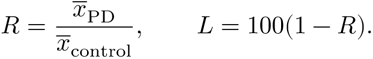

Here, *x̄* is the group mean count or density, *R* the ratio of means and *L* the percentage loss. Positive loss indicates lower counts or densities in PD; negative loss indicates higher values. These are cross-sectional differences relative to controls. Marker-defined counts can also change with marker expression.

We fitted a Bayesian multilevel model with separate random intercepts for study and population. The study intercept captures variation shared by measurements from the same paper, including tissue preparation and other study conditions; the population intercept represents average loss in each population. Partial pooling draws uncertain estimates towards the overall distribution while retaining population differences supported by the data. Thus, the opposing striatal tyrosine hydroxylase-positive (TH+) findings of Porritt et al. (2006) and Huot et al. (2007) contribute to one population estimate despite differences between their protocols. Sampling variances were treated as fixed, and sampling errors as independent conditional on the model effects.

We transformed posterior mean log ratios to obtain population estimates of percentage loss and posterior quantiles to obtain 95% credible intervals. Published anatomical and biological annotations linked measured populations to proposed explanations of vulnerability. Figure 3 groups populations by transmitter identity, classical pigmentation, experimentally described physiological properties and anatomical location. Axonal annotations draw on human and non-human primate histology, population-derived length estimates and rodent reconstructions; evidence on metabolism and firing comes from animal or culture experiments. These properties were not measured in the counted human donors. The comparisons describe how estimated loss relates to proposed features; they do not establish equivalence between populations, adjudicate every version of a mechanism or adjust populations to a common disease stage (Supplementary Methods: Biological annotations).

### Robustness checks

We added counting method and the adapted risk-of-bias score as predictors in separate analyses. To assess sensitivity to inferred neuron types, we compared fits using all labels with fits restricted to reported labels. We examined small-study effects by adding standard error as a predictor to the study–population model. Supplementary Methods gives the specifications and limitations of these analyses.

### Precision planning

We used SampleSizeMeans 1.2.3 to estimate how many additional independent cohorts of 10 PD and 10 control donors would be needed for each population’s mean loss to reach a conditional 95% interval width of at most 30 percentage points (Joseph and Bélisle 2023). This target, approximately ±15 percentage points, defines a working resolution for descriptive mapping rather than a threshold for biological importance or evidence of loss. All 146 fitted populations were included, regardless of the direction of the estimated effect; sensitivity analyses used widths of 20 and 40 points. We approximated each existing log-ratio posterior by a normal distribution and held its mean and the variance of future cohort estimates fixed. The latter combines posterior mean between-study variance with sampling variance estimated from observed coefficients of variation. The calculations estimate precision under these assumptions, rather than the probability that future data will attain the target. We report study replication separately because partial pooling can yield a narrow interval from limited direct evidence. Full assumptions and results are given in Supplementary Methods: Precision planning and Supplementary Table 12.

### Reproducibility

Data preparation and analysis use R, and the manuscript is generated with Quarto and knitr. The repository contains curated extractions, mapping rules, analysis scripts and recorded package versions. Access arrangements are described in the Data and Code Availability statement.

## Results

### Evidence remains concentrated in a few regions

We identified 166 post-mortem studies of Parkinson’s disease published between 1963 and 2025. Of these, 152 contributed to the meta-analysis, representing 146 neuronal populations in 85 brain regions. Fourteen studies lacked comparable quantitative PD and control values (Supplementary Figure 2; Supplementary Table 3, Table 4). Less precise estimates were associated with greater neuronal loss in the study–population model (standard-error slope -0.80 on the log-ratio scale; 95% credible interval, -1.29 to -0.29; Supplementary Figure 3).

Research focused on the substantia nigra (SN) (66 studies), locus coeruleus (LC) (20), basal nucleus (BN) (14) and ventral tegmental area (VTA) (10). Among broader anatomical groups, the midbrain appeared in 79 studies, compared with 22 for the pons, 16 for the medulla, ten for the cortex and three for the cerebellum (Fig. 1a). Together, the studies covered only 18% of countable brain atlas labels (109 of 597; Supplementary Figure 1). Five studies examined enteric populations.

Evidence was also concentrated in a few neuronal classes. Dopaminergic (DA) neurons were examined in 75 studies, cholinergic (Ch) neurons in 23 and noradrenergic (NA) neurons in 19. GABAergic (GABA) and serotonergic (5-HT) neurons appeared in ten and five studies, respectively, while glutamatergic neurons and Purkinje cells each appeared in three (Fig. 1b). A further ten studies reported tyrosine hydroxylase-positive (TH+) counts and seven reported neuromelanin-pigmented (NM) counts without assignment to a transmitter class. Twenty-one studies reported neurons without a specified type, predominantly using general histological stains such as cresyl violet, Nissl, and haematoxylin and eosin (H&E) (Fig. 1b). Glycinergic, histaminergic and nitrergic populations were each examined in one study.

This concentration persisted across the publication record. By 2025, SN DA neurons had been examined in 62 studies, compared with 19 for LC NA, 12 for BN Ch and nine for VTA DA neurons (Fig. 1c). Of the 158 region–cell populations recorded in the review, 127 (80%) appeared in only one study and 31 in two or more. Six decades of counting therefore provided repeated observations for a few populations while leaving most without replication.

### Substantial estimated loss extends beyond the best-studied populations

Estimates across all 146 modelled populations show a wide range of neuronal deficits (Fig. 2a,b). Among populations studied more than once, SN DA and LC NA neurons had the largest estimated losses: 67% (95% credible interval, 63–70%) and 64% (58–68%), respectively. Several populations represented by one study also had large estimated losses, including DA neurons of the myenteric nerve plexus (Myenteric) (64%, 41–79%) and substance P-positive (SP+) neurons in the medullary reticular formation (Med RF) (61%, 41–75%) and median raphe (MnR) (60%, 44–72%). Two studies supported an estimated 48% loss in parafascicular nucleus of thalamus (Pf) neurons (31–60%; Fig. 2a).

**Figure 2:**
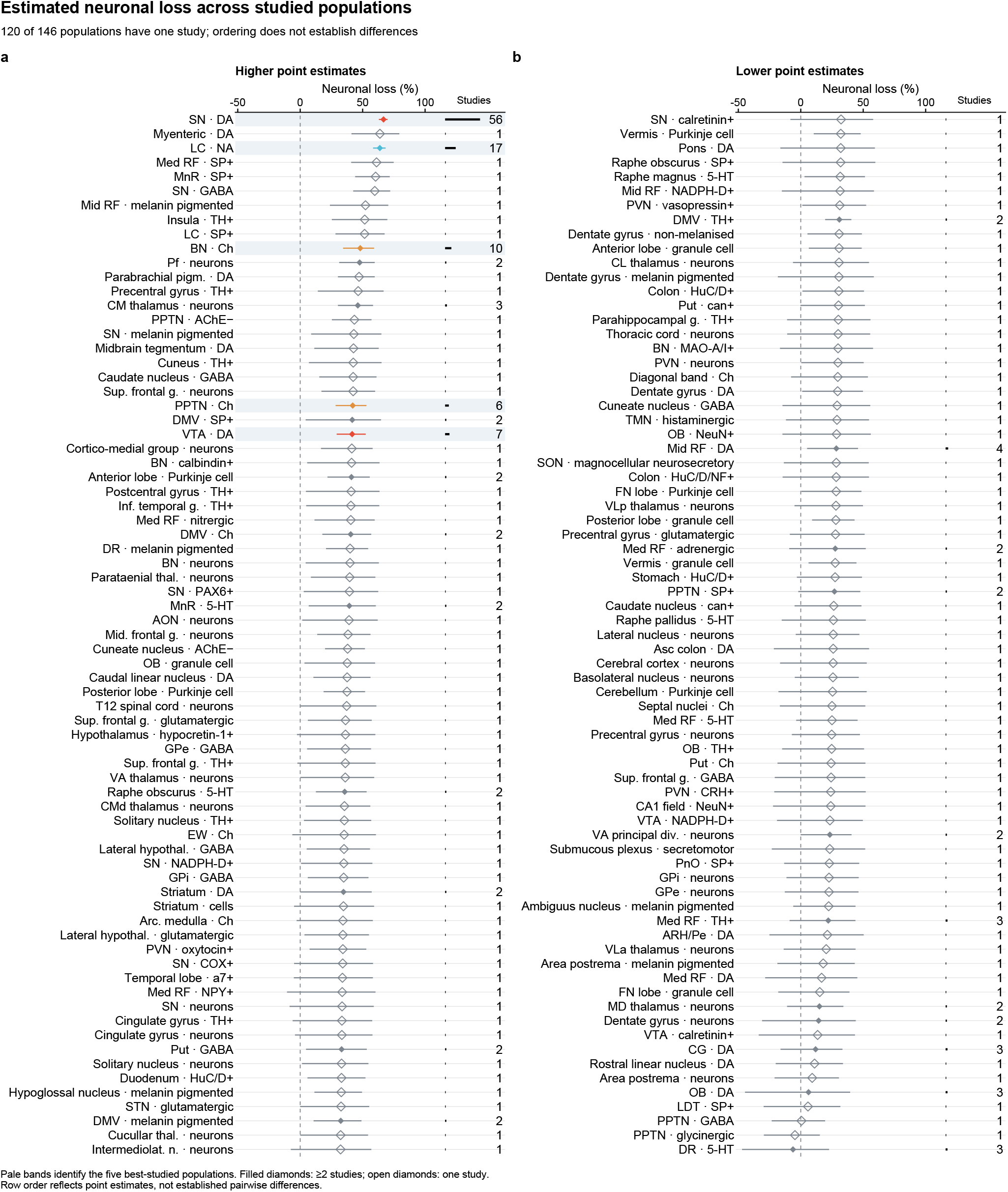
Estimated neuronal loss and 95% credible intervals across all 146 fitted neuronal populations. Pale bands highlight the five most-studied populations, coloured by transmitter class. Filled diamonds denote two or more studies; open diamonds denote one. Bars and numbers at the right show the number of contributing studies. Populations are ordered by point estimate, without implying established pairwise differences. Cell labels retain source-study marker definitions. Supplementary Table 11 provides the complete label key and abbreviations.

Some population estimates combined opposing observations. The two studies of intrinsic striatal TH+ neurons reported approximately 56% higher and 78% lower counts in PD than in controls (Porritt et al. 2006; Huot et al. 2007). Partial pooling drew the combined estimate towards the shared distribution, so it should be interpreted alongside the disagreement between studies.

The olfactory bulb (OB) DA estimate was imprecise: 6% loss, with an interval spanning both higher and lower counts in PD (-45 to 40%; Fig. 2b). All three contributing studies reported 13–108% higher counts in PD than in controls (Huisman et al. 2004, 2008; Mundiñano et al. 2011), but partial pooling shifted the population estimate towards loss. A separate study reported lower calretinin-positive interneuron density alongside unchanged TH+ density (Cave et al. 2016). These findings indicate uncertainty in the cellular pattern underlying olfactory dysfunction, which can precede motor symptoms in PD (Ross et al. 2008).

### Population comparisons constrain proposed explanations of vulnerability

Substantial loss extended beyond the pigmented midbrain populations that shaped early accounts of selective vulnerability (Hirsch et al. 1988). LC NA and BN Ch neurons showed substantial loss outside the DA class, while estimates within the DA class varied: 42% (29–52%) in VTA, 12% (-16 to 33%) in central grey substance of midbrain (CG), and 6% (-45 to 40%) in OB neurons (Fig. 3a). Dopaminergic identity is therefore not necessary for substantial loss, limiting the scope of explanations based on dopamine and its reactive metabolites (Masato et al. 2019). Similarly, BN Ch neurons showed 48% (34–59%) loss outside the classical pigmented catecholaminergic populations (Fig. 3b). Together with affected pigmented SN DA and LC NA populations, this extends the observed deficits beyond those emphasised in neuromelanin-based accounts (Whitehouse et al. 1983; Filimontseva et al. 2025).

**Figure 3:**
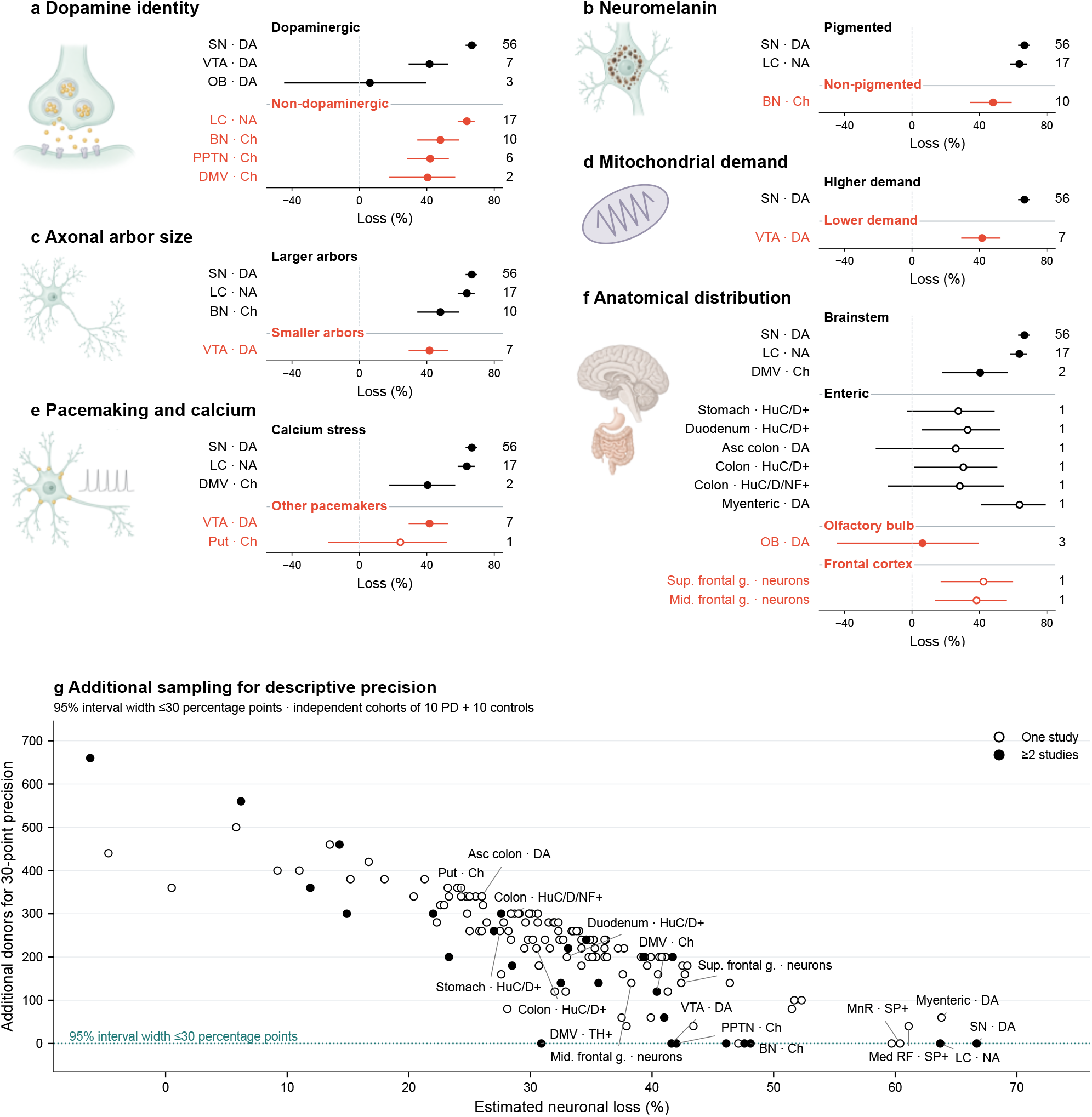
Neuronal loss and additional sampling across populations. Panels a–f show pooled loss and 95% credible intervals; panel g shows additional sampling needed for a 30-percentage-point precision target. Full legend follows. **a–f,** Neuronal loss across populations grouped by dopamine identity (a), neuromelanin (b), axonal arbor size (c), mitochondrial demand (d), pacemaking and calcium stress (e), and anatomical location (f). Points and bars show pooled loss and 95% credible intervals; negative values indicate higher counts or densities in PD. Numbers at the right show contributing studies. Open points denote one study and filled points at least two. Black and red distinguish reference and comparison groups; colour does not indicate rejection of a hypothesis. Intervals spanning zero leave loss unresolved. “Smaller arbors” and “Lower demand” describe VTA relative to SN. “Other pacemakers” does not imply absence of calcium signalling, and “Non-pigmented” follows the conventional population classification. Results and Supplementary Methods: Biological annotations describe the human and non-human evidence underlying these labels. The properties were not measured alongside counts in the analysed donors, and anatomical location does not establish temporal spread. Illustrations are schematic; labels follow Supplementary Table 11. **g,** Estimated neuronal loss against additional donors needed for a conditional 95% interval width of at most 30 percentage points, approximately ±15 points. All 146 fitted populations are shown, regardless of the direction of estimated loss. Open points identify populations represented by one study and filled points those represented by multiple studies. Points on the dotted zero line already meet the precision target under the calculation. Labels identify selected populations; precision alone does not establish replication or biological importance. Planning assumes independent cohorts of 10 PD and 10 control donors, a fixed log-ratio posterior centre and a fixed cohort variance. New data may change both the effect estimate and heterogeneity. The calculations describe conditional precision requirements rather than assurance probabilities or detection power, and do not establish cell death. Full results and sensitivity analyses at widths of 20 and 40 points are given in Supplementary Table 12 and Supplementary Methods.

Human and chimpanzee cortical axon-density measurements and basal-forebrain neuron counts suggest extensive axonal arbors in Ch neurons (Raghanti et al. 2008; Raghanti et al. 2011; Wu et al. 2014). These population-derived estimates, considered alongside substantial BN Ch loss, are consistent with the proposed link between axonal extent and vulnerability (Fig. 3c). However, arbor size and maintenance demands were not measured in the counted donors, so the comparison does not establish a mechanistic relationship.

Calcium-linked stress during autonomous pacemaking has been proposed to explain neuronal vulnerability (Guzman et al. 2010; Sánchez-Padilla et al. 2014; Goldberg et al. 2012). Human counts show loss in SN DA, LC NA and DMV Ch populations associated with this mechanism in experimental models, but also in VTA DA neurons, whose pacemaking can persist during calcium-current blockade (Khaliq and Bean 2010). The comparison in Fig. 3e limits explanations based on calcium-dependent pacemaking alone, although the physiological evidence comes from rodents rather than the counted human donors.

The anatomical comparison identified affected brainstem and enteric populations. Gastrointestinal loss estimates ranged from 26% in ascending-colon DA neurons to 64% in myenteric DA neurons (Fig. 3f), although each population was represented by one study and several intervals spanned no loss. The OB provides a rostral comparison: its DA loss estimate remained uncertain, while counts suggested neuronal loss in the superior and middle frontal gyri (Halliday and McCann 2005). These findings concern regions relevant to Braak staging and brain-first/gut-first accounts (Braak et al. 2003; Borghammer and Van Den Berge 2019), but anatomical proximity and cross-sectional deficits cannot establish the direction or sequence of degeneration. Clinical information was available in 105 of 166 studies (63%), but inconsistent reporting prevented adjustment to a common level of clinical severity.

### Precision of population estimates

We chose a 95% interval-width target of 30 percentage points as a practical resolution for interpreting neuronal loss in broadly low, moderate or high terms, rather than as a formal classification threshold (Methods: Precision planning). Only 11 of 146 populations met this target. Eight were represented by multiple studies, including SN DA, LC NA, BN Ch, VTA DA and PPTN Ch. The remaining three—MnR SP+, parabrachial pigmented nucleus DA and SN GABA—each relied on one study, distinguishing a precise model estimate from independently replicated evidence.

For the 135 populations falling short of the target, the median conditional requirement was 240 additional donors (range 40–660). Myenteric DA neurons, reported by Singaram (1996), required an estimated 60 additional donors, while medullary reticular formation SP+ neurons, reported by Halliday, Li, et al. (1990), required 40 (Fig. 3g). These estimates describe the sampling needed to narrow uncertainty around the magnitude of loss under the assumptions specified in Methods: Precision planning and Supplementary Methods. Two populations met the narrower 20-point target and 26 met the broader 40-point target (Supplementary Table 12).

### Robustness to study methods and annotation

Adjustment for counting method changed the overall loss estimate from 33.3% to 32.7%. Excluding inferred neuron-type labels gave 36.8% loss, compared with 33.3% in the corresponding all-data analysis (Supplementary Table 13). Higher risk-of-bias checklist scores were associated with lower estimated loss (log-ratio slope 0.064 per point; 95% credible interval, 0.012–0.117; Supplementary Table 14). These associations cannot isolate the contribution of individual methodological features.

## Discussion

Selective vulnerability has increasingly served as a broad explanation for neuronal loss rather than a hypothesis tested against explicit predictions. The “regional selectivity” described in the early 1990s has expanded to encompass anatomical, chemical and molecular features, yet the evidence remains concentrated in a few neuronal populations. Funnel asymmetry and fragmented sampling strengthen the case for quantitative synthesis beyond the earlier narrative review (Giguère et al. 2018), while also limiting what the pooled evidence can establish.

Six decades of counting have covered only 18% of countable brain atlas labels, and most recorded populations have been examined only once. Repeated study has established degeneration in familiar targets without establishing how exceptional those targets are. The resulting map reflects both biological vulnerability and scientific attention, limiting claims about the distribution of selectivity across the brain. Nevertheless, substantial LC NA and BN Ch loss shows that dopaminergic identity is not necessary for marked degeneration; BN Ch loss also extends the deficits beyond classically neuromelanin-containing populations. These features may contribute to vulnerability, but neither is necessary for substantial loss across all affected populations.

Targeted replication could establish whether several less-studied populations consistently show substantial loss. Estimated losses were 64% in myenteric DA neurons (95% credible interval, 41–79%; Singaram 1996), 61% in medullary reticular formation SP+ neurons (41–75%; Halliday, Li, et al. 1990), 60% in SN GABA neurons (42–72%; Martirosyan et al. 2024) and 60% in median raphe SP+ neurons (44–72%; Gai et al. 1991). Conditional estimates suggest that 60 additional donors for myenteric DA and 40 for medullary reticular formation SP+ neurons would bring these two populations to the precision target of a 95% credible interval width $ $30 percentage points. Replication would strengthen evidence for these deficits, although establishing selectivity would also require comparisons with potentially spared populations.

Post-mortem counts leave the origins of these deficits unresolved. The Fearnley–Lees reanalysis found lower counts in PD but did not establish a steeper age-associated decline. Differences established before the sampled ages therefore remain compatible with the data (Fearnley and Lees 1991). Earlier degeneration, developmental differences and age-related failure of compensation remain possibilities that cross-sectional counts cannot distinguish (Schwamborn 2018). Genetic evidence implicating enteric neurons and oligodendrocytes identifies further targets for investigation, without implying that their contribution involves cell death (Bryois et al. 2020). Stability of the aggregate estimate across counting-method and annotation analyses does not resolve these temporal uncertainties. A convincing account must explain both where neuronal deficits occur and how they arise.

This synthesis supports substantial loss beyond SN DA neurons, while leaving regional and cellular selectivity incompletely characterised. Progress requires precise, replicated estimates across affected and potentially spared populations, followed by direct comparisons that test explicit mechanistic predictions. New counting should therefore address the uncertainties that limit interpretation, rather than simply identify further populations with neuronal loss.

## Acknowledgements

Nathan Skene is funded by the UK Dementia Research Institute and UKRI Future Leaders Fellowship [grant number MR/T04327X/1].

JPTH is a National Institute for Health and Care Research (NIHR) Senior Investigator (NIHR203807). JPTH was supported by NIHR Bristol Biomedical Research Centre at University Hospitals Bristol and Weston NHS Foundation Trust and the University of Bristol and is a member of the MRC Integrative Epidemiology Unit at the University of Bristol. JPTH and SD were supported by the NIHR Applied Research Collaboration West (ARC West) at University Hospitals Bristol and Weston NHS Foundation Trust. The views expressed in this article are those of the authors and do not necessarily represent those of the NHS, the NIHR, MRC, or the Department of Health and Social Care.

Samuel Burke is funded by the FRQS doctoral scholarship, Brain Canada and ASAP.

## Author Contributions

Will Lunt and Jay A. Moore contributed equally to this work and are joint first authors.

Investigation: WL, MS, JC and JAM screened records; WL, MS, JC, JAM and SB assessed full-text eligibility and risk of bias; WL, MS, JC, JS and JAM extracted data. Data curation: WL, MS, JC, JS and JAM. Validation: JAM and SB. JAM verified extractions and corrected risk-of-bias assessments against source records; SB adjudicated screening disagreements. Supervision and funding acquisition: NGS. Writing, review and editing: all authors.

## Competing Interests

The authors declare no competing interests.

## Data and Code Availability

Curated study extractions, analysis datasets, mapping rules, code and manuscript sources are maintained in the SelectiveVulnerabilityMetaAnalysis repository. The reproduction guide provides separate instructions for regenerating the manuscript from supplied fitted results and refitting models from curated inputs, including dependencies, model settings and verification commands. Versioned bundles record source revisions and file checksums, and the website uses the same analysis outputs. The review is registered with PROSPERO (CRD42021265515).

## Ethics Statement

This study is a systematic review and meta-analysis of previously published data from post-mortem studies. No new human tissue was collected or analysed. Ethics approval was not required.

## Supplementary Methods

### Review records and additional extraction details

We also excluded cohorts explicitly labelled young-onset PD. The short-duration striatal group in Porritt et al. (2006), recorded as “PD (short; < 9 years)”, remained in the extraction database but was excluded from the meta-analysis.

Search strategies and update exports are archived in the study repository and listed in Supplementary Table 6. MEDLINE, EMBASE and Web of Science exports dated 18 December 2025 match the recorded update totals. The strategy document was modified on 21 December 2025, but the exact executed search histories remain unverified. Twenty-six retrieval records could not be assigned a final disposition because of duplicate identifiers and outdated access decisions (Supplementary Figure 2). Pre-consensus screening decisions were not systematically retained, so the records do not establish how many independent assessments were made for every full text or risk-of-bias rating.

Risk-of-bias scores counted affirmative responses to eight adapted checklist items, omitting missing values, and ranged from 1 to 8. In 2025, JAM checked selected assessments against source records and revised them where needed. This was a targeted verification exercise rather than a new blinded assessment by two reviewers.

Checking the Fearnley–Lees source corrected the extracted ventromedial label to total caudal substantia nigra (Fearnley and Lees 1991). We also unified inconsistent marker labels before calculating group summaries.

We accessed the shared extraction database on 21 January 2026. In addition to the fields listed in the main Methods, clinical data included disease duration, age at onset, Braak PD and neurofibrillary-tangle stage, Hoehn-Yahr stage, UPDRS, MMSE, Lewy body scores and dementia status. Availability required at least one recorded value for these disease or cognitive variables; age at death, sex and derived rescaled fields did not qualify. Such information was available in 105 of 166 papers (63%), but inconsistent reporting prevented adjustment to a common disease stage or duration.

### Missing variability and group summaries

We derived standard deviations from reported variability measures where possible and treated values greater than ten times the absolute group mean as missing. To impute missing values, we first used the median coefficient of variation (standard deviation divided by mean) from the same paper. If unavailable, we used the median for the same region–neuron-type combination, followed by the dataset-wide 75th percentile. Multiplying the selected coefficient by the group mean gave the imputed standard deviation. The curated extraction table identifies all imputations. The analysis included 22 effects from 11 studies with at least one imputed standard deviation.

Where individual observations were available, we calculated group summaries within each recorded diagnostic group, region, neuron type and marker. When preparing comparisons, we combined repeated grouped records by averaging means, rounding the average sample size, and taking the square root of the average squared standard deviation. This affected 19 selected effects from 17 studies and combined alternative anatomical or marker-defined summaries (Supplementary Table 15). The procedure neither reconstructed the variance of the averaged measurements nor pooled independent cohorts by sample size. We treated the independent PD faller and nonfaller groups differently: these were pooled using their sample sizes, exact published means and standard errors, including variation between group means (Karachi et al. 2010). Their shared control group was retained once.

### Selection of measurements within studies

When a study reported several eligible measurements of the same population, we selected one using the following order, introduced in April 2026:

1. Counting method: stereology, then computer-assisted counting, then manual counting.
2. Marker: the marker preferred for that neuron type in the predefined mapping, followed by surrogate stains. For example, ChAT took priority over cresyl violet for Ch neurons, and TH or DBH over neuromelanin alone for NA neurons. These selection rules do not establish marker specificity in every anatomical setting.
3. Sample size: the larger combined PD and control sample broke remaining ties.

Supplementary Table 16 lists the complete marker priorities. Unlisted combinations receive surrogate priority.

### Effect size derivation

We calculated log ratios of mean neuronal counts or densities in PD relative to controls, with their sampling variances, using metafor::escalc(measure = “ROM”) (Viechtbauer 2010). PD and control sample sizes, means and standard deviations were supplied as the first and second groups, respectively. We used the function’s default correction and variance settings.

### Bayesian model

We fitted the model with brms using yi | se(sei, sigma = FALSE) ∼ 1 + (1 | study_id) + (1 | region_cell). The study and population effects are separate normally distributed random intercepts. Here, yi is the corrected log ratio, sei its estimated standard error, study_id the publication identifier (PMID), and region_cell the region–neuron-type population. Sampling variances were treated as fixed, with no additional residual variance estimated. Population point estimates were calculated as 100[1 − exp (*θ̄*), where *θ̄* is the posterior mean population log ratio. We transformed the 2.5th and 97.5th posterior percentiles to obtain 95% credible intervals, reversing their order because percentage loss decreases as the log ratio increases.

The intercept prior was Student-t(3, −0.5, 2.5), and study and population standard deviations had half-Student-t(3, 0, 2.5) priors. We ran four chains of 4,000 iterations, each with 2,000 warmup iterations, retaining 8,000 draws. Sampler settings were adapt_delta = 0.95, max_treedepth = 12 and seed = 42. The study repository provides convergence diagnostics with the fitted results. Model fitting used brms 2.23.0, rstan 2.32.7 and StanHeaders 2.32.10; posterior summaries used posterior 1.7.0.

The grand mean summarises the sampled population labels; it does not estimate the proportion of all neurons lost in PD.

### Risk of bias

We added the adapted checklist score as a continuous predictor in the Bayesian model and summarised its slope and 95% credible interval (Supplementary Table 14).

### Counting method

We entered counting method as a three-level fixed factor, using manual counting as the reference and retaining study and population random intercepts. Source studies were classified as manual, computer-assisted or stereological. These categories describe reported practice; computer assistance alone does not establish a different counting principle. The model used the sampler settings specified above. Coefficient priors and diagnostics accompany the fitted output in the repository.

Supplementary Table 17 reports model-adjusted estimates. Because counting method varied with publication year, checklist score and the populations studied, these comparisons describe associations rather than causal effects of counting technique.

### Anatomical harmonisation and unresolved labels

We mapped brain-region labels to the Allen Brain Atlas ontology (Hawrylycz et al. 2012). The following source checks informed the anatomical and measurement decisions summarised in the main Methods:

- SNc and unspecified SN DA measurements were combined.
- Paranigral DA measurements were combined with VTA measurements because the paranigral nucleus is an A10 subnucleus (Javoy-Agid et al. 1984; McRitchie et al. 1997).
- Whole-midbrain DA counts from German et al. (1989) were excluded because they combine A8, A9 and A10 populations, including the SN.
- Nigral m-calpain measurements from Mouatt-Prigent et al. (1996) were excluded because they report marker-upregulation density rather than neuron number.

Other corrections to extracted neuron-type and anatomical labels are documented in Supplementary Table 5 and the mapping files.

Some labels remain heterogeneous or overlapping:

- Nigral marker-defined labels, including calbindin-positive, calretinin-positive, NADPH-d-negative and COX-positive neurons, can describe subsets of the broader DA population.
- We retained substance P (SP)-positive and TH+ labels separately. SP immunoreactivity alone does not establish catecholaminergic identity; published brainstem studies distinguish SP-containing from monoamine-defined populations (Halliday, Blumbergs, et al. 1990).
- Whole-striatum labels retain measurements from papers that did not distinguish caudate from putamen (Put).
- Dorsal motor nucleus labels retain heterogeneous markers, including TH, substance P and pigmentation. These may identify catecholaminergic or co-expressing subsets, so the labels should not be treated as independent populations.

### Biological annotations

Figure 3 presents descriptive comparisons relevant to proposed mechanisms. Transmitter classes follow the curated population definitions. SN DA and LC NA represent the classical pigmented catecholaminergic groups, with BN Ch providing a comparison outside these groups (Hirsch et al. 1988; Whitehouse et al. 1983; Filimontseva et al. 2025). Pigment content varies within populations, and historical labels based on pigmented cells do not confirm neuromelanin chemically. The comparison therefore concerns conventional population classifications, not a pigment-dose response.

Axonal arbor size describes the total extent and branching of an axon. Reconstructions of rat SN DA and mouse basal forebrain Ch neurons provide direct evidence of extensive arbors (Matsuda et al. 2009; Wu et al. 2014). Human and non-human primate evidence comes from measurements of cortical Ch axon-length density in humans, chimpanzees and macaques (Raghanti et al. 2008) and nucleus-basalis neuron counts across primates (Raghanti et al. 2011). Combining published axon density, cortical volume and neuron counts, Wu et al. (2014) estimated mean total Ch axon lengths of approximately 107 m in humans and 31 m in chimpanzees. These estimates assume that sampled cortical fibres arise from the specified source population and that sampled densities extend across the target volume. They are not direct reconstructions of individual axons or measurements from the PD donors in this review. Reconstructions of 132 LC neurons from four mice also show extensive axons (mean 34.20 cm), but fewer branches per unit length than in the comparison set of other central neurons (Su et al. 2026). Thus, large total extent need not imply dense branching. In vivo mouse labelling supports larger SN than VTA striatal arbors, estimated from labelled fibre coverage normalised by labelled neuron counts (Giguère et al. 2019). This measures the striatal axonal domain rather than complete single-neuron length. In mouse neuronal cultures, SN DA neurons had more complex arbors, higher basal respiration and less respiratory reserve (Pacelli et al. 2015). Together, these findings motivate the prediction of greater loss with greater arbor-maintenance demand. We applied no common quantitative threshold and did not infer metabolic demand or protein clearance from observed loss.

The pacemaking comparison distinguishes autonomous activity from calcium-linked mitochondrial stress. Mouse experiments provide evidence of such stress in SN DA, LC NA and DMV Ch neurons (Guzman et al. 2010; Sánchez-Padilla et al. 2014; Goldberg et al. 2012). Mouse VTA DA neurons provide a comparison in which autonomous firing persists during calcium-current blockade (Khaliq and Bean 2010). Rat striatal Ch interneurons provide another example of autonomous firing (Bennett and Wilson 1999), mapped at class level to human Put Ch neurons. Neither comparison establishes absence of calcium signalling or oxidative stress. The proposed mechanism concerns calcium-associated stress rather than spontaneous firing alone.

Anatomical groups identify brainstem, enteric, olfactory and frontal cortical populations relevant to staging and body-first/brain-first accounts (Braak et al. 2003; Borghammer and Van Den Berge 2019), without assigning a sequence of neuronal loss. Across panels, the annotated properties were not measured alongside counts in the analysed donors, and missing evidence was not coded as absence. Every selected estimate is shown with its study count and credible interval. An interval spanning zero leaves loss unresolved; it does not establish preservation. Populations recur across panels because the proposed mechanisms overlap, not because they provide independent confirmations.

### Precision planning

The main target was a total conditional 95% credible-interval width of 30 percentage points for mean percentage loss, approximately ±15 points. This defines a working resolution for descriptive mapping, rather than a validated standard for a well-characterised population. A precisely estimated small or absent reduction can also meet the target, so all 146 fitted populations were included regardless of the direction of estimated loss. We repeated the calculation at widths of 20 and 40 percentage points to assess sensitivity to the chosen resolution (Supplementary Table 12).

We used the known-variance normal-mean procedure in SampleSizeMeans 1.2.3 (Joseph and Bélisle 2023). Let and denote the mean and variance of the existing posterior for a population’s log PD-to-control ratio. Each new independent cohort comprises 10 PD and 10 control donors and contributes one log-ratio estimate. Its planning variance is *A* = E(*τ*^2^) + *c*/10, where *τ^2^* is the fitted between-study variance and is the sum of squared median PD and control coefficients of variation in the analysed data. The global sampling-variance approximation is held fixed across populations.

For a target percentage-point width *w*, the corresponding log-ratio width at the fixed mean is ℓ = 2 asinh {*w*/(200*e*^*m*^)} . We supplied this length, cohort precision 1/*A*, prior-equivalent cohort count *A*/*v*0 and credibility level 0.95 to SampleSizeMeans::mu.varknown. We rounded upwards to whole cohorts and multiplied by 20 for additional donors. After cohorts, the conditional variance is *v*_*k*_ = (1/*v*_0_ + *k*/*A*)−1 and the transformed width is 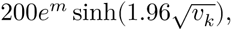, using the exact normal quantile in code. Intervals are asymmetric after transformation. We verified that each reported design meets the requested width and that one fewer cohort does not, unless the target is already met at zero additional cohorts.

The calculation fixes the future interval centre at the current posterior mean and treats cohort variance as known. Because new data may change both the mean and heterogeneity, the results are conditional sampling scenarios rather than expected widths, assurance probabilities or guaranteed recruitment requirements. The shared variance approximation may also be unsuitable for a particular protocol. Partial pooling can yield narrow intervals for populations represented by one study; meeting the target therefore establishes neither replication nor freedom from bias, and does not demonstrate cell death. The criterion concerns one population’s mean, rather than a pairwise difference or a sequential stopping rule. Donor requirements cannot be summed across populations because one tissue collection can supply several measurements. Code and full calculations are retained in the repository.

### Small-study effects

We added effect-size standard error as a continuous predictor to the study–population Bayesian model (Egger et al. 1997) and summarised its slope with a 95% credible interval. A negative slope on the log-ratio scale indicates greater reported loss in less precise studies. The contour-enhanced funnel plot shows the observed estimates and fitted relationship with precision (Supplementary Figure 3). This association cannot distinguish publication bias from other causes of funnel asymmetry.

### Inferred neuron-type labels

We compared the full dataset with the subset excluding inferred neuron-type labels, fitting both with the same frequentist REML population specification. These estimates are reported with confidence intervals rather than Bayesian credible intervals (Supplementary Table 13).

## Supplementary Figures

**Supplementary Figure 1:**
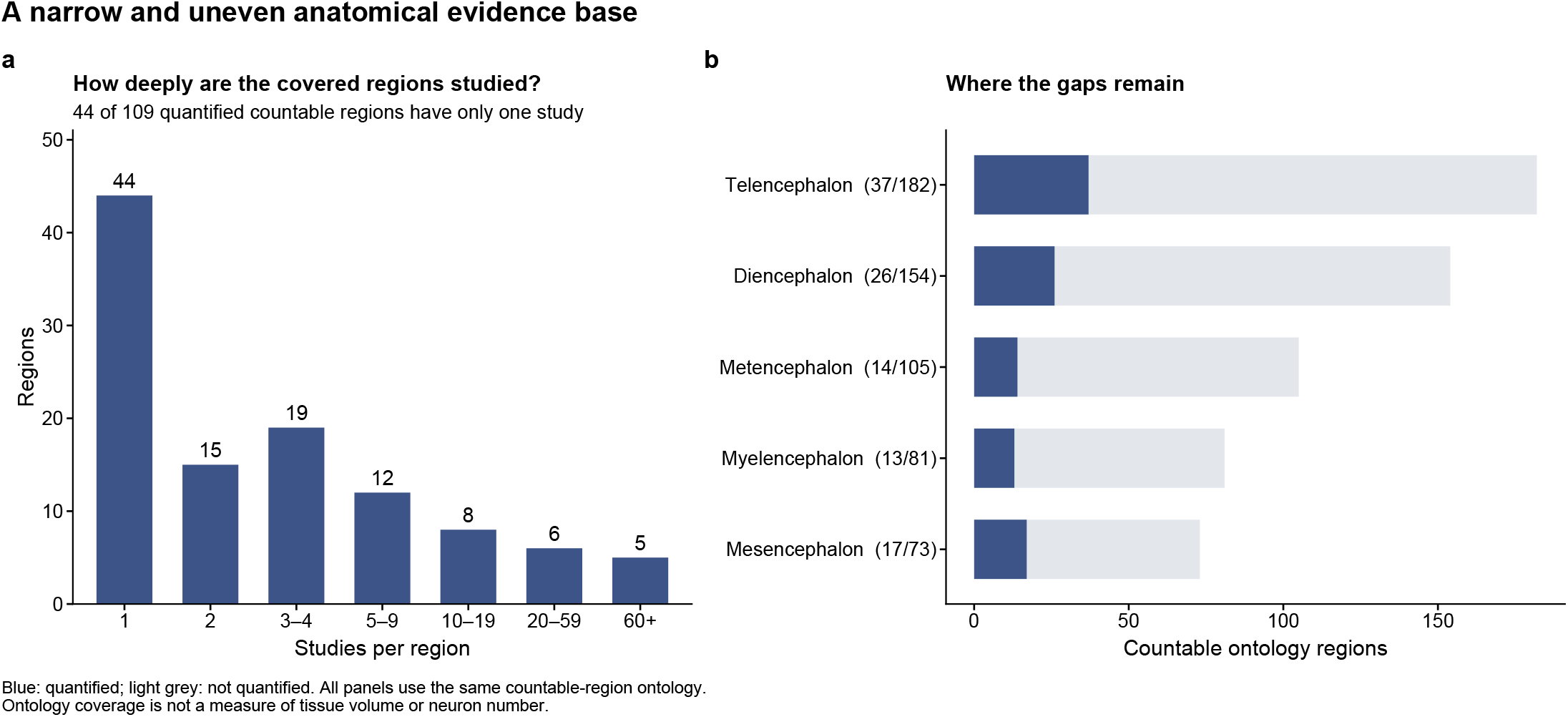
Anatomical coverage and replication in the PD neuron-count literature. (a) Study counts for 109 quantified countable Allen Brain Atlas labels, of which 44 have one study. (b) Quantified (blue) and total (grey) countable labels in five anatomical divisions. The aggregate Top-Level Structures and Grey Matter entries, each represented by one quantified entry, are omitted from this panel but retained in the overall coverage and replication totals. Non-neuronal ontology entries are excluded throughout. Coverage measures atlas labels rather than tissue volume or neuron number.

**Supplementary Figure 2:**
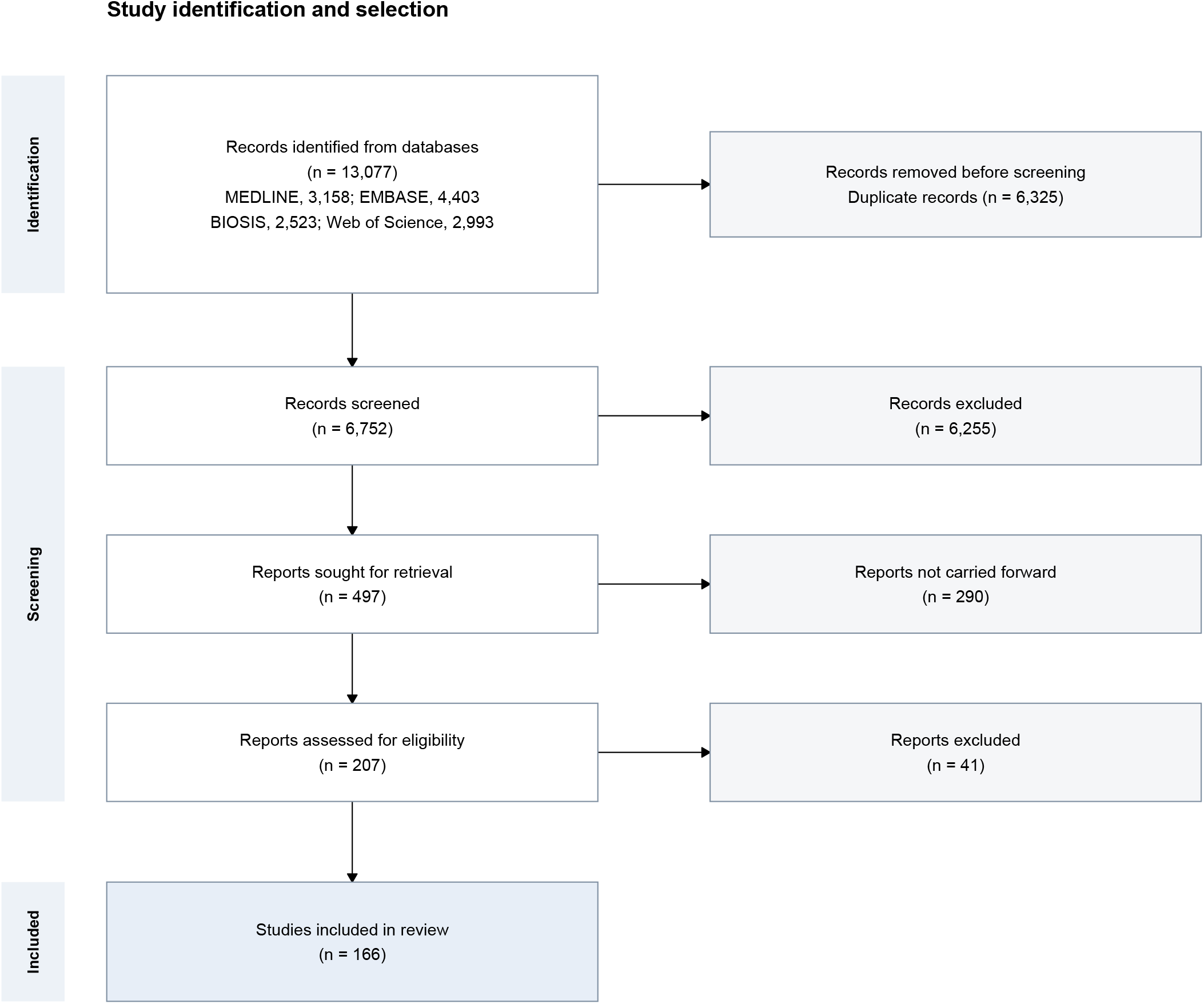
Neuropathological study retrieval process. The PRISMA flow diagram shows 13,077 records identified through systematic review, 6,752 records screened and 166 eligible studies included. Database totals combine the original and update rows in the screening workbook, correcting its combined BIOSIS total. The number of reports not carried forward is calculated as reports sought minus reports proceeding to eligibility assessment. It includes recorded retrieval-stage exclusions and the unresolved workbook discrepancy. Original counts and unresolved dispositions are documented in the source audit; no exclusion reason has been inferred.

**Supplementary Figure 3:**
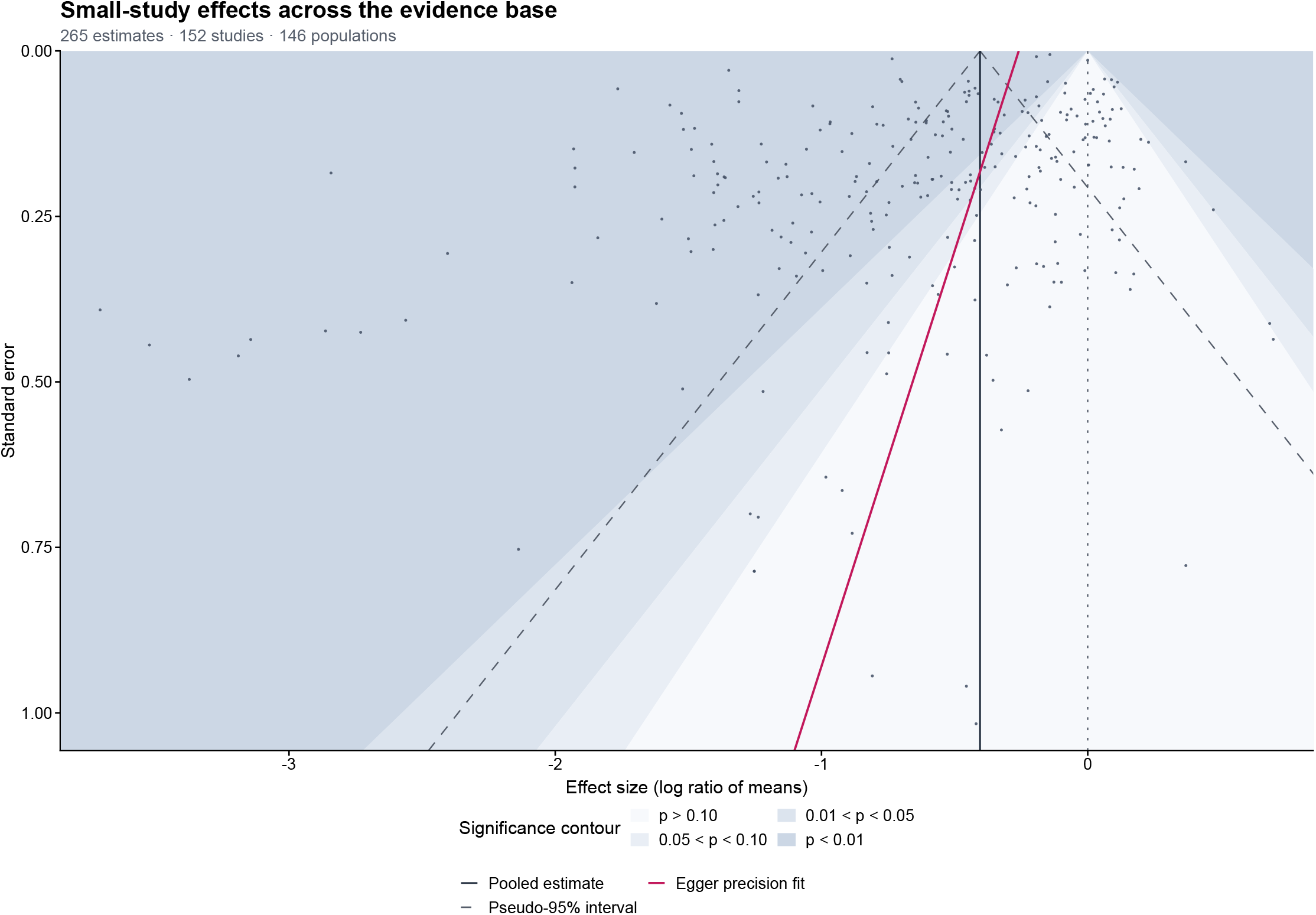
Small-study effects across 265 estimates from 152 studies and 146 populations. Points show log ratios of means against standard error, with more precise estimates at the top. Shading marks two-sided significance contours around the null (dotted line). The solid dark line marks the study–population model pooled estimate (-0.404 log ratio of means; 33.3% loss), with a dashed pseudo-95% interval. The crimson line shows the Bayesian Egger-style fit: slope -0.80 (95% CrI -1.29 to -0.29; posterior probability of a positive slope 0.002). Its intercept at zero standard error corresponds to 22.8% loss (95% CrI 12.1 to 32.5%). Asymmetry describes an association between precision and effect size; the intercept is not a bias-corrected estimate.

## Supplementary Tables

**Table 1:**
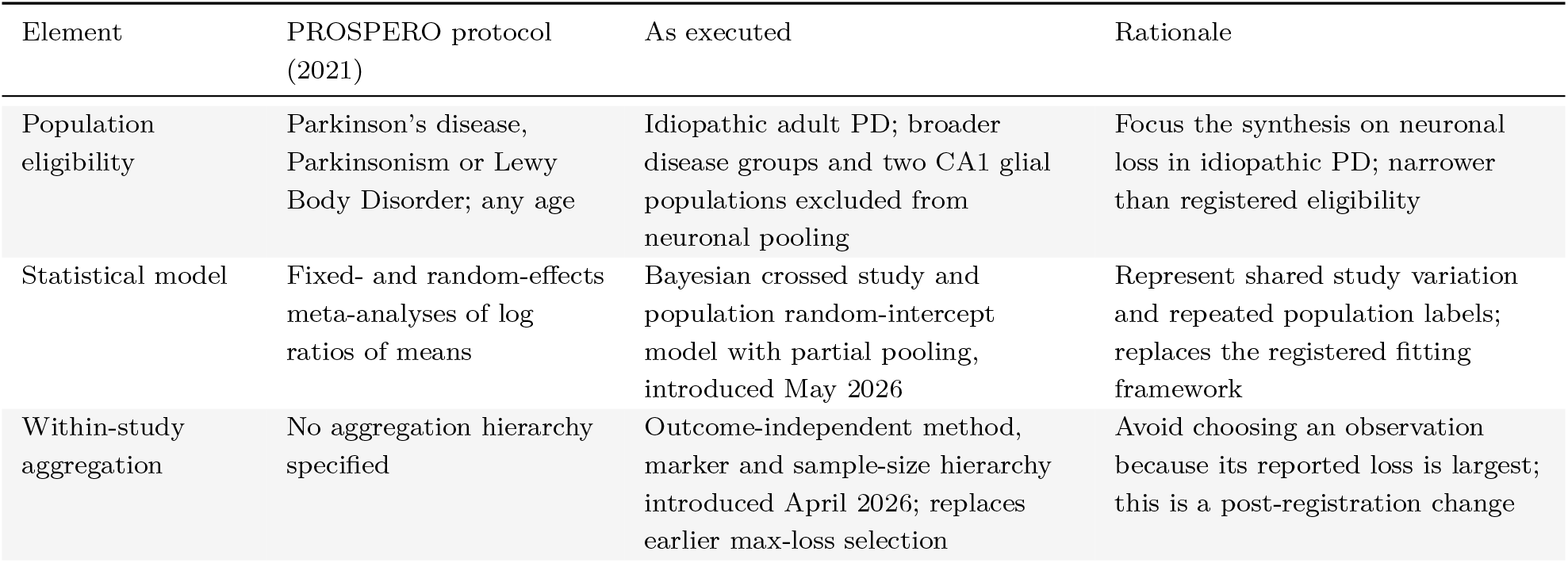
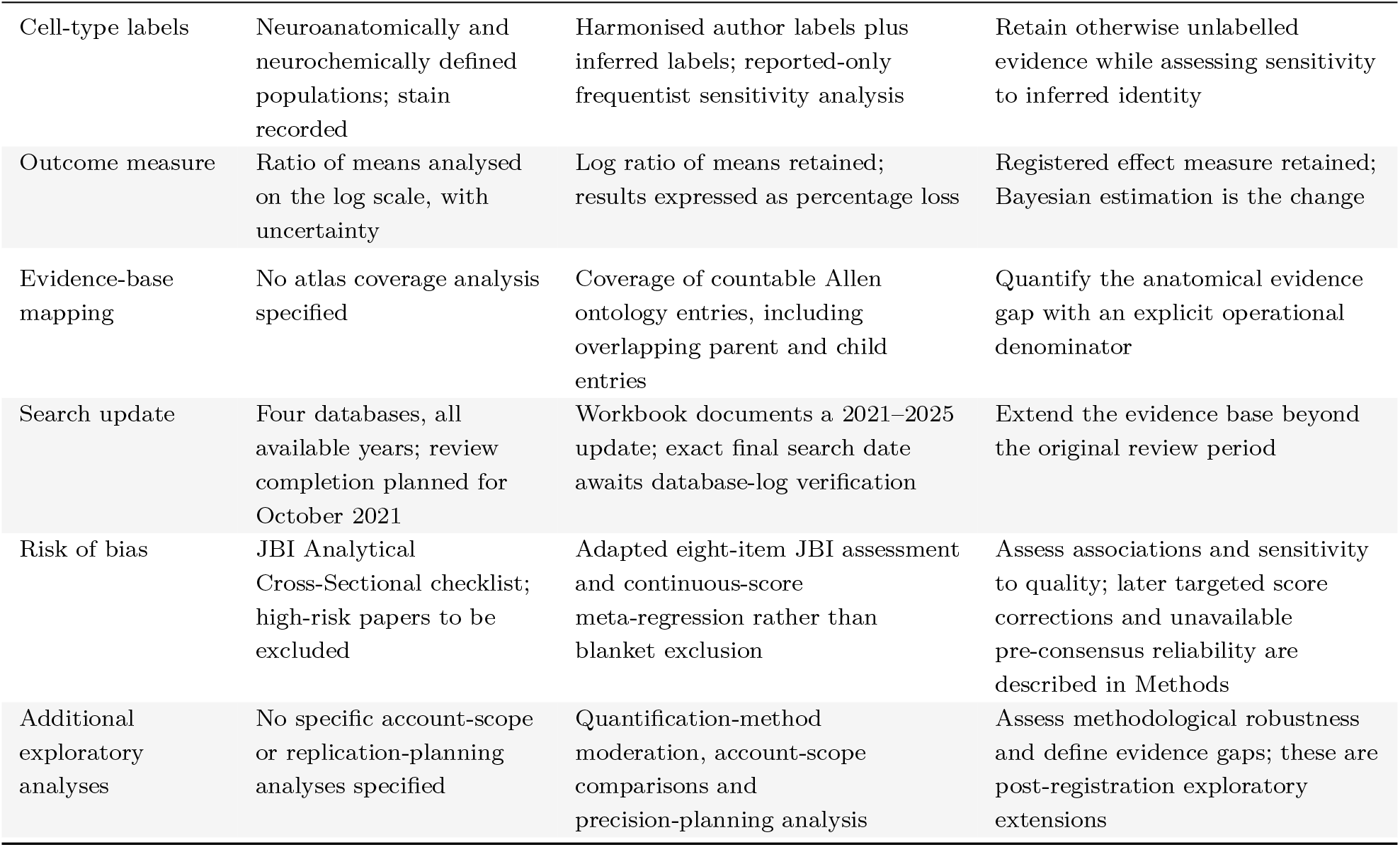
Changes from the registered protocol (PROSPERO CRD42021265515, registered 13 July 2021). The saved registration and repository history distinguish registered plans from later decisions.

| Element | PROSPERO protocol (2021) | As executed | Rationale |
| --- | --- | --- | --- |
| Population eligibility | Parkinson’s disease, Parkinsonism or Lewy Body Disorder; any age | Idiopathic adult PD; broader disease groups and two CA1 glial populations excluded from neuronal pooling | Focus the synthesis on neuronal loss in idiopathic PD; narrower than registered eligibility |
| Statistical model | Fixed- and random-effects meta-analyses of log ratios of means | Bayesian crossed study and population random-intercept model with partial pooling, introduced May 2026 | Represent shared study variation and repeated population labels; replaces the registered fitting framework |
| Within-study aggregation | No aggregation hierarchy specified | Outcome-independent method, marker and sample-size hierarchy introduced April 2026; replaces earlier max-loss selection | Avoid choosing an observation because its reported loss is largest; this is a post-registration change |
| Cell-type labels | Neuroanatomically and neurochemically defined populations; stain recorded | Harmonised author labels plus inferred labels; reported-only frequentist sensitivity analysis | Retain otherwise unlabelled evidence while assessing sensitivity to inferred identity |
| Outcome measure | Ratio of means analysed on the log scale, with uncertainty | Log ratio of means retained; results expressed as percentage loss | Registered effect measure retained; Bayesian estimation is the change |
| Evidence-base mapping | No atlas coverage analysis specified | Coverage of countable Allen ontology entries, including overlapping parent and child entries | Quantify the anatomical evidence gap with an explicit operational denominator |
| Search update | Four databases, all available years; review completion planned for October 2021 | Workbook documents a 2021–2025 update; exact final search date awaits database-log verification | Extend the evidence base beyond the original review period |
| Risk of bias | JBI Analytical Cross-Sectional checklist; high-risk papers to be excluded | Adapted eight-item JBI assessment and continuous-score meta-regression rather than blanket exclusion | Assess associations and sensitivity to quality; later targeted score corrections and unavailable pre-consensus reliability are described in Methods |
| Additional exploratory analyses | No specific account-scope or replication-planning analyses specified | Quantification-method moderation, account-scope comparisons and precision-planning analysis | Assess methodological robustness and define evidence gaps; these are post-registration exploratory extensions |

**Table 2:**
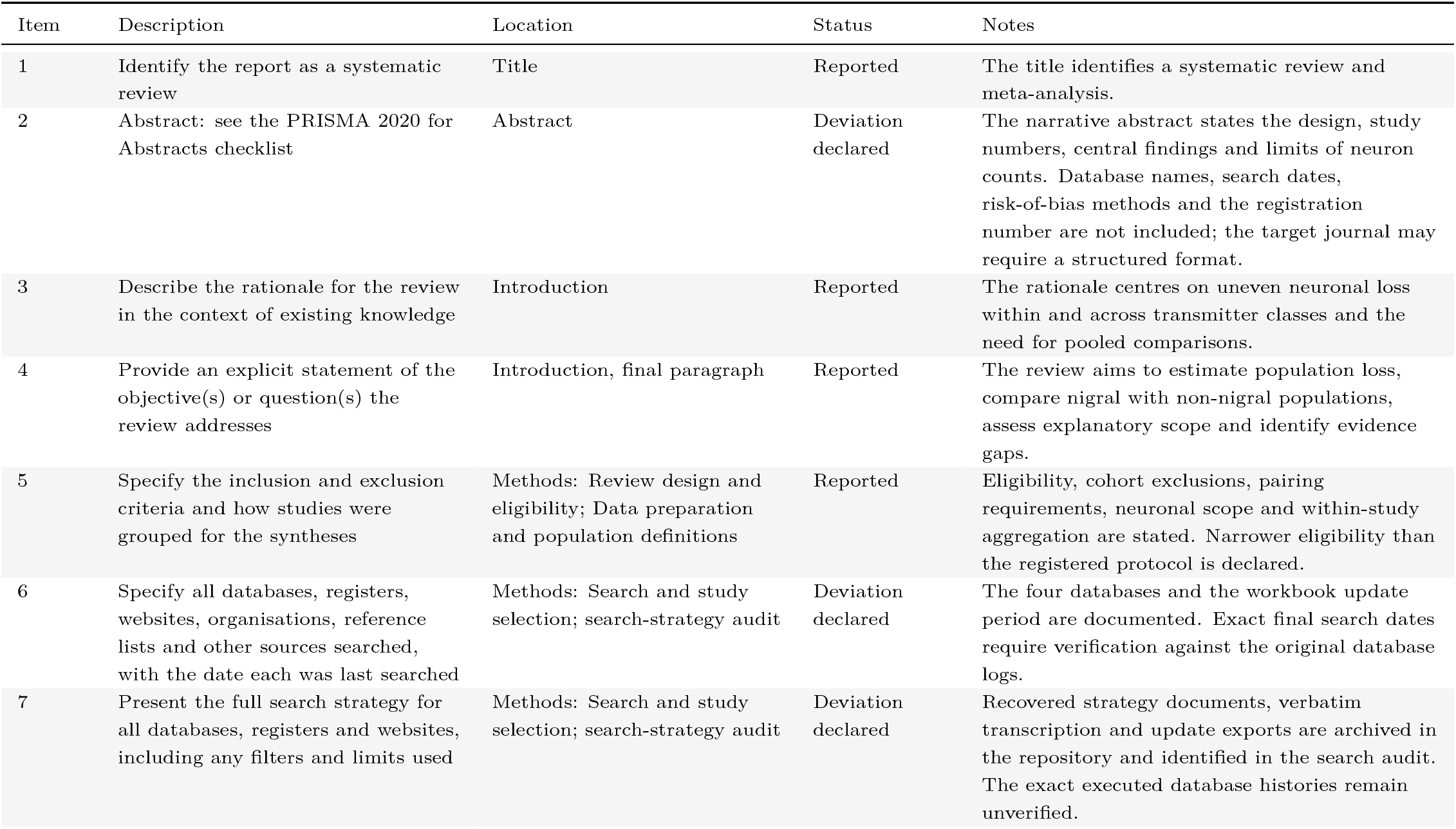
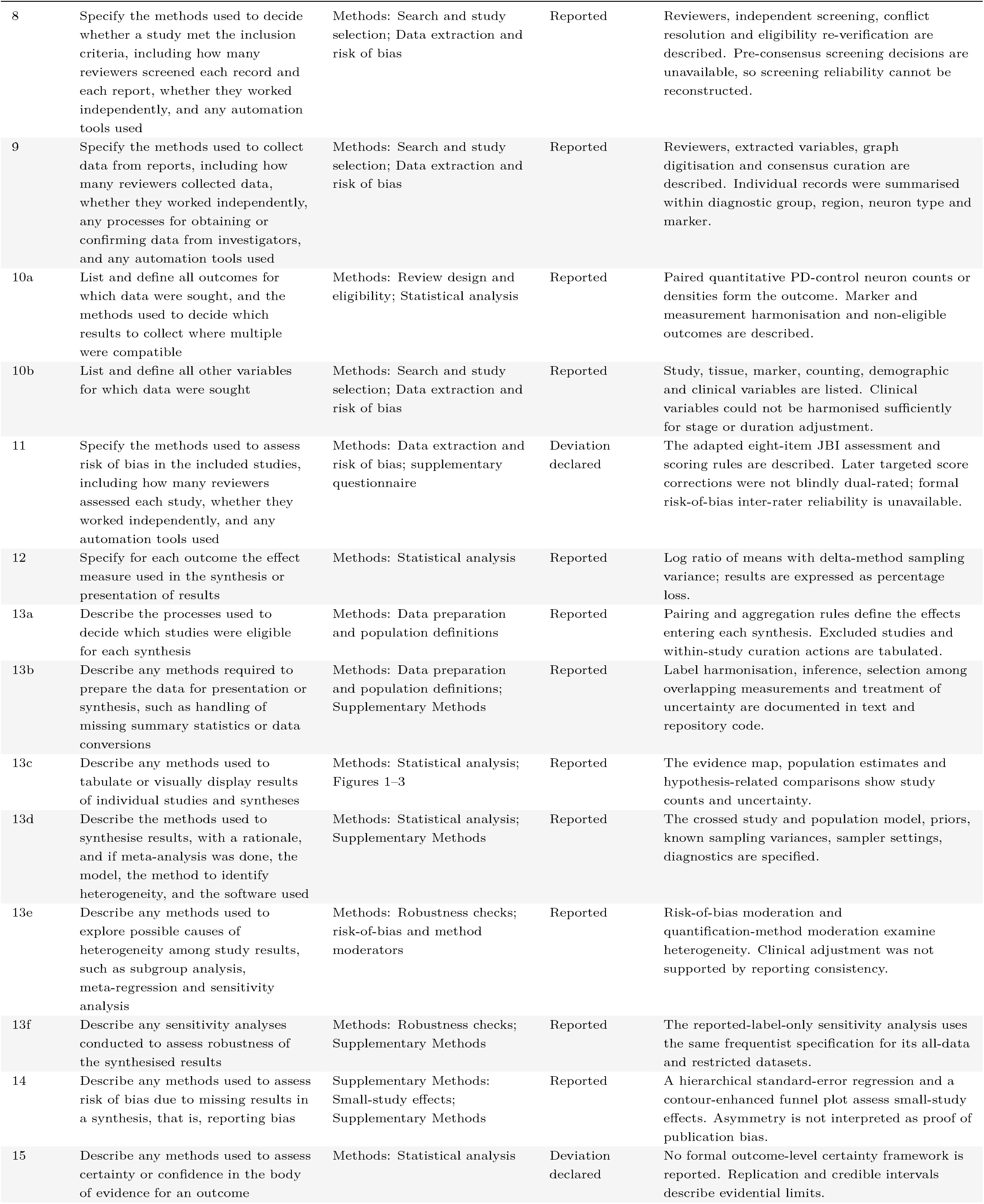
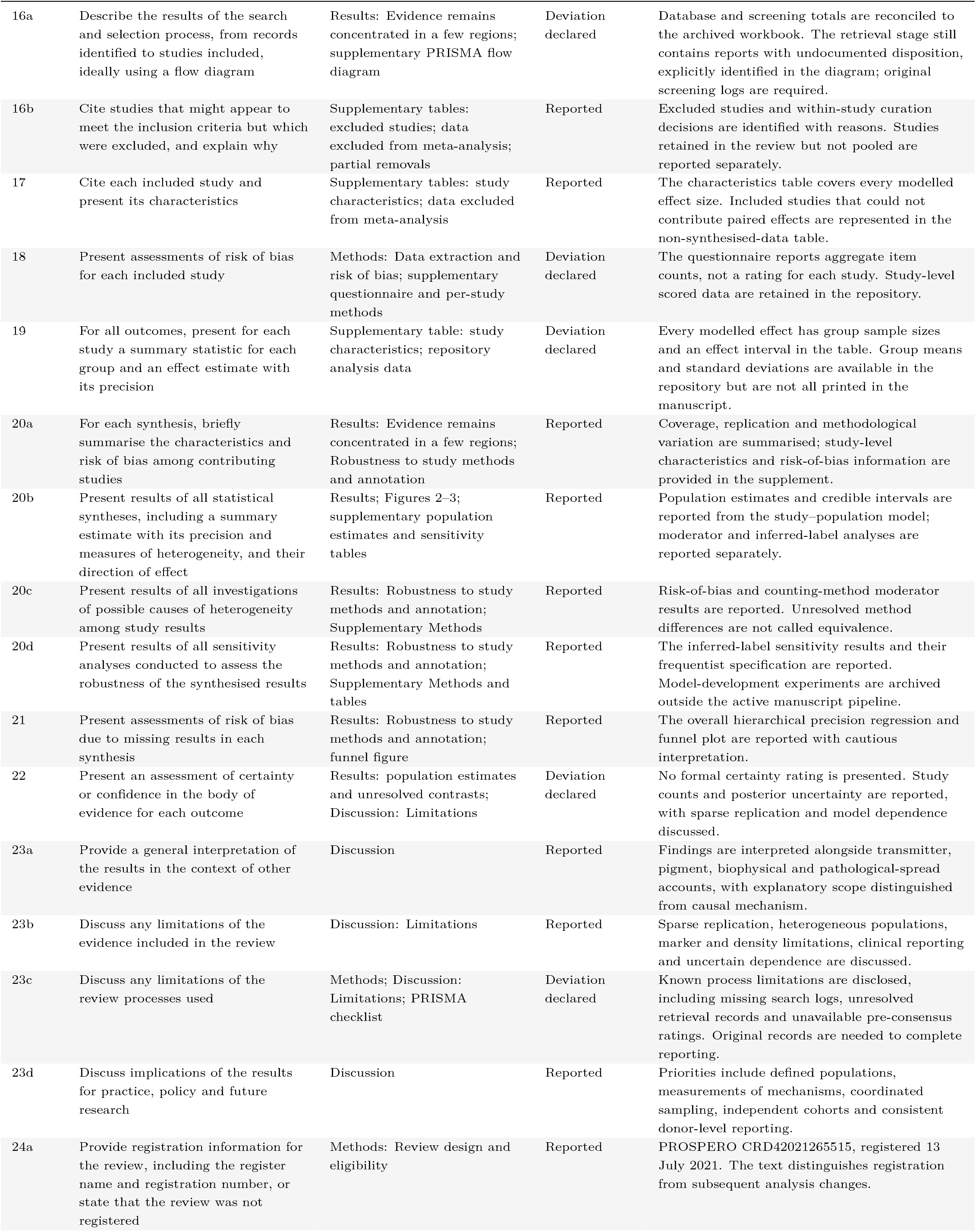
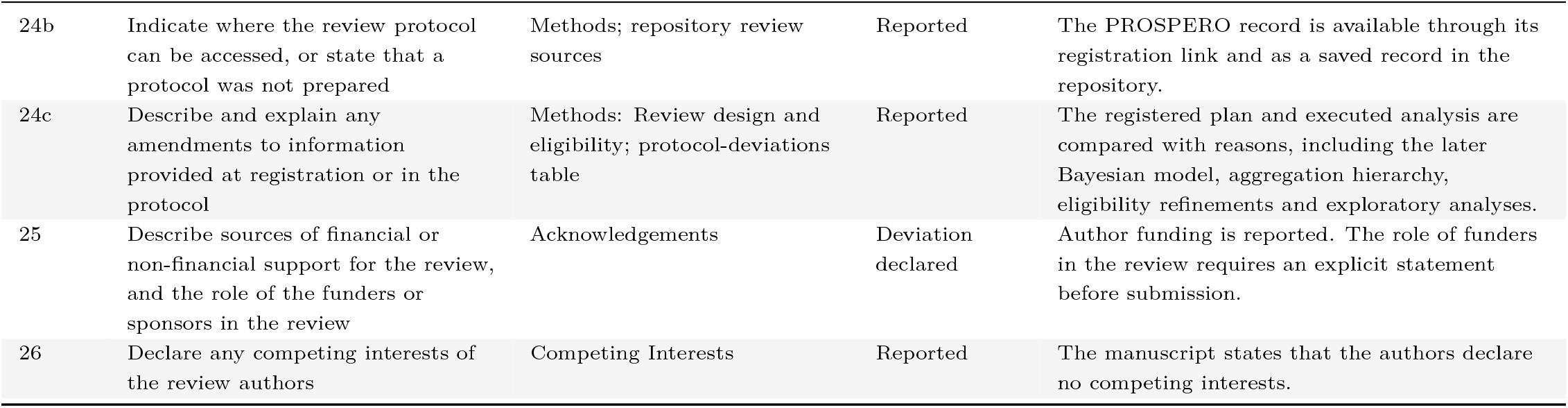
PRISMA 2020 checklist. Items are mapped to the revised manuscript and assessed against available records. Outstanding items concern search details, retrieval records and aspects of reviewer independence. Reporting inter-rater kappa is not itself a PRISMA requirement.

| Item | Description | Location | Status | Notes |
| --- | --- | --- | --- | --- |
| 1 | Identify the report as a systematic review | Title | Reported | The title identifies a systematic review and meta-analysis. |
| 2 | Abstract: see the PRISMA 2020 for Abstracts checklist | Abstract | Deviation declared | The narrative abstract states the design, study numbers, central findings and limits of neuron counts. Database names, search dates, risk-of-bias methods and the registration number are not included; the target journal may require a structured format. |
| 3 | Describe the rationale for the review in the context of existing knowledge | Introduction | Reported | The rationale centres on uneven neuronal loss within and across transmitter classes and the need for pooled comparisons. |
| 4 | Provide an explicit statement of the objective(s) or question(s) the review addresses | Introduction, final paragraph | Reported | The review aims to estimate population loss, compare nigral with non-nigral populations, assess explanatory scope and identify evidence gaps. |
| 5 | Specify the inclusion and exclusion criteria and how studies were grouped for the syntheses | Methods: Review design and eligibility; Data preparation and population definitions | Reported | Eligibility, cohort exclusions, pairing requirements, neuronal scope and within-study aggregation are stated. Narrower eligibility than the registered protocol is declared. |
| 6 | Specify all databases, registers, websites, organisations, reference lists and other sources searched, with the date each was last searched | Methods: Search and study selection; search-strategy audit | Deviation declared | The four databases and the workbook update period are documented. Exact final search dates require verification against the original database logs. |
| 7 | Present the full search strategy for all databases, registers and websites, including any filters and limits used | Methods: Search and study selection; search-strategy audit | Deviation declared | Recovered strategy documents, verbatim transcription and update exports are archived in the repository and identified in the search audit. The exact executed database histories remain unverified. |
| 8 | Specify the methods used to decide whether a study met the inclusion criteria, including how many reviewers screened each record and each report, whether they worked independently, and any automation tools used | Methods: Search and study selection; Data extraction and risk of bias | Reported | Reviewers, independent screening, conflict resolution and eligibility re-verification are described. Pre-consensus screening decisions are unavailable, so screening reliability cannot be reconstructed. |
| 9 | Specify the methods used to collect data from reports, including how many reviewers collected data, whether they worked independently, any processes for obtaining or confirming data from investigators, and any automation tools used | Methods: Search and study selection; Data extraction and risk of bias | Reported | Reviewers, extracted variables, graph digitisation and consensus curation are described. Individual records were summarised within diagnostic group, region, neuron type and marker. |
| 10a | List and define all outcomes for which data were sought, and the methods used to decide which results to collect where multiple were compatible | Methods: Review design and eligibility; Statistical analysis | Reported | Paired quantitative PD-control neuron counts or densities form the outcome. Marker and measurement harmonisation and non-eligible outcomes are described. |
| 10b | List and define all other variables for which data were sought | Methods: Search and study selection; Data extraction and risk of bias | Reported | Study, tissue, marker, counting, demographic and clinical variables are listed. Clinical variables could not be harmonised sufficiently for stage or duration adjustment. |
| 11 | Specify the methods used to assess risk of bias in the included studies, including how many reviewers assessed each study, whether they worked independently, and any automation tools used | Methods: Data extraction and risk of bias; supplementary questionnaire | Deviation declared | The adapted eight-item JBI assessment and scoring rules are described. Later targeted score corrections were not blindly dual-rated; formal risk-of-bias inter-rater reliability is unavailable. |
| 12 | Specify for each outcome the effect measure used in the synthesis or presentation of results | Methods: Statistical analysis | Reported | Log ratio of means with delta-method sampling variance; results are expressed as percentage loss. |
| 13a | Describe the processes used to decide which studies were eligible for each synthesis | Methods: Data preparation and population definitions | Reported | Pairing and aggregation rules define the effects entering each synthesis. Excluded studies and within-study curation actions are tabulated. |
| 13b | Describe any methods required to prepare the data for presentation or synthesis, such as handling of missing summary statistics or data conversions | Methods: Data preparation and population definitions; Supplementary Methods | Reported | Label harmonisation, inference, selection among overlapping measurements and treatment of uncertainty are documented in text and repository code. |
| 13c | Describe any methods used to tabulate or visually display results of individual studies and syntheses | Methods: Statistical analysis; Figures 1–3 | Reported | The evidence map, population estimates and hypothesis-related comparisons show study counts and uncertainty. |
| 13d | Describe the methods used to synthesise results, with a rationale, and if meta-analysis was done, the model, the method to identify heterogeneity, and the software used | Methods: Statistical analysis; Supplementary Methods | Reported | The crossed study and population model, priors, known sampling variances, sampler settings, diagnostics are specified. |
| 13e | Describe any methods used to explore possible causes of heterogeneity among study results, such as subgroup analysis, meta-regression and sensitivity analysis | Methods: Robustness checks; risk-of-bias and method moderators | Reported | Risk-of-bias moderation and quantification-method moderation examine heterogeneity. Clinical adjustment was not supported by reporting consistency. |
| 13f | Describe any sensitivity analyses conducted to assess robustness of the synthesised results | Methods: Robustness checks; Supplementary Methods | Reported | The reported-label-only sensitivity analysis uses the same frequentist specification for its all-data and restricted datasets. |
| 14 | Describe any methods used to assess risk of bias due to missing results in a synthesis, that is, reporting bias | Supplementary Methods: Small-study effects; Supplementary Methods | Reported | A hierarchical standard-error regression and a contour-enhanced funnel plot assess small-study effects. Asymmetry is not interpreted as proof of publication bias. |
| 15 | Describe any methods used to assess certainty or confidence in the body of evidence for an outcome | Methods: Statistical analysis | Deviation declared | No formal outcome-level certainty framework is reported. Replication and credible intervals describe evidential limits. |
| 16a | Describe the results of the search and selection process, from records identified to studies included, ideally using a flow diagram | Results: Evidence remains concentrated in a few regions; supplementary PRISMA flow diagram | Deviation declared | Database and screening totals are reconciled to the archived workbook. The retrieval stage still contains reports with undocumented disposition, explicitly identified in the diagram; original screening logs are required. |
| 16b | Cite studies that might appear to meet the inclusion criteria but which were excluded, and explain why | Supplementary tables: excluded studies; data excluded from meta-analysis; partial removals | Reported | Excluded studies and within-study curation decisions are identified with reasons. Studies retained in the review but not pooled are reported separately. |
| 17 | Cite each included study and present its characteristics | Supplementary tables: study characteristics; data excluded from meta-analysis | Reported | The characteristics table covers every modelled effect size. Included studies that could not contribute paired effects are represented in the non-synthesised-data table. |
| 18 | Present assessments of risk of bias for each included study | Methods: Data extraction and risk of bias; supplementary questionnaire and per-study methods | Deviation declared | The questionnaire reports aggregate item counts, not a rating for each study. Study-level scored data are retained in the repository. |
| 19 | For all outcomes, present for each study a summary statistic for each group and an effect estimate with its precision | Supplementary table: study characteristics; repository analysis data | Deviation declared | Every modelled effect has group sample sizes and an effect interval in the table. Group means and standard deviations are available in the repository but are not all printed in the manuscript. |
| 20a | For each synthesis, briefly summarise the characteristics and risk of bias among contributing studies | Results: Evidence remains concentrated in a few regions; Robustness to study methods and annotation | Reported | Coverage, replication and methodological variation are summarised; study-level characteristics and risk-of-bias information are provided in the supplement. |
| 20b | Present results of all statistical syntheses, including a summary estimate with its precision and measures of heterogeneity, and their direction of effect | Results; Figures 2–3; supplementary population estimates and sensitivity tables | Reported | Population estimates and credible intervals are reported from the study–population model; moderator and inferred-label analyses are reported separately. |
| 20c | Present results of all investigations of possible causes of heterogeneity among study results | Results: Robustness to study methods and annotation; Supplementary Methods | Reported | Risk-of-bias and counting-method moderator results are reported. Unresolved method differences are not called equivalence. |
| 20d | Present results of all sensitivity analyses conducted to assess the robustness of the synthesised results | Results: Robustness to study methods and annotation; Supplementary Methods and tables | Reported | The inferred-label sensitivity results and their frequentist specification are reported. Model-development experiments are archived outside the active manuscript pipeline. |
| 21 | Present assessments of risk of bias due to missing results in each synthesis | Results: Robustness to study methods and annotation; funnel figure | Reported | The overall hierarchical precision regression and funnel plot are reported with cautious interpretation. |
| 22 | Present an assessment of certainty or confidence in the body of evidence for each outcome | Results: population estimates and unresolved contrasts; Discussion: Limitations | Deviation declared | No formal certainty rating is presented. Study counts and posterior uncertainty are reported, with sparse replication and model dependence discussed. |
| 23a | Provide a general interpretation of the results in the context of other evidence | Discussion | Reported | Findings are interpreted alongside transmitter, pigment, biophysical and pathological-spread accounts, with explanatory scope distinguished from causal mechanism. |
| 23b | Discuss any limitations of the evidence included in the review | Discussion: Limitations | Reported | Sparse replication, heterogeneous populations, marker and density limitations, clinical reporting and uncertain dependence are discussed. |
| 23c | Discuss any limitations of the review processes used | Methods; Discussion: Limitations; PRISMA checklist | Deviation declared | Known process limitations are disclosed, including missing search logs, unresolved retrieval records and unavailable pre-consensus ratings. Original records are needed to complete reporting. |
| 23d | Discuss implications of the results for practice, policy and future research | Discussion | Reported | Priorities include defined populations, measurements of mechanisms, coordinated sampling, independent cohorts and consistent donor-level reporting. |
| 24a | Provide registration information for the review, including the register name and registration number, or state that the review was not registered | Methods: Review design and eligibility | Reported | PROSPERO CRD42021265515, registered 13 July 2021. The text distinguishes registration from subsequent analysis changes. |
| 24b | Indicate where the review protocol can be accessed, or state that a protocol was not prepared | Methods; repository review sources | Reported | The PROSPERO record is available through its registration link and as a saved record in the repository. |
| 24c | Describe and explain any amendments to information provided at registration or in the protocol | Methods: Review design and eligibility; protocol-deviations table | Reported | The registered plan and executed analysis are compared with reasons, including the later Bayesian model, aggregation hierarchy, eligibility refinements and exploratory analyses. |
| 25 | Describe sources of financial or non-financial support for the review, and the role of the funders or sponsors in the review | Acknowledgements | Deviation declared | Author funding is reported. The role of funders in the review requires an explicit statement before submission. |
| 26 | Declare any competing interests of the review authors | Competing Interests | Reported | The manuscript states that the authors declare no competing interests. |

**Table 3:**
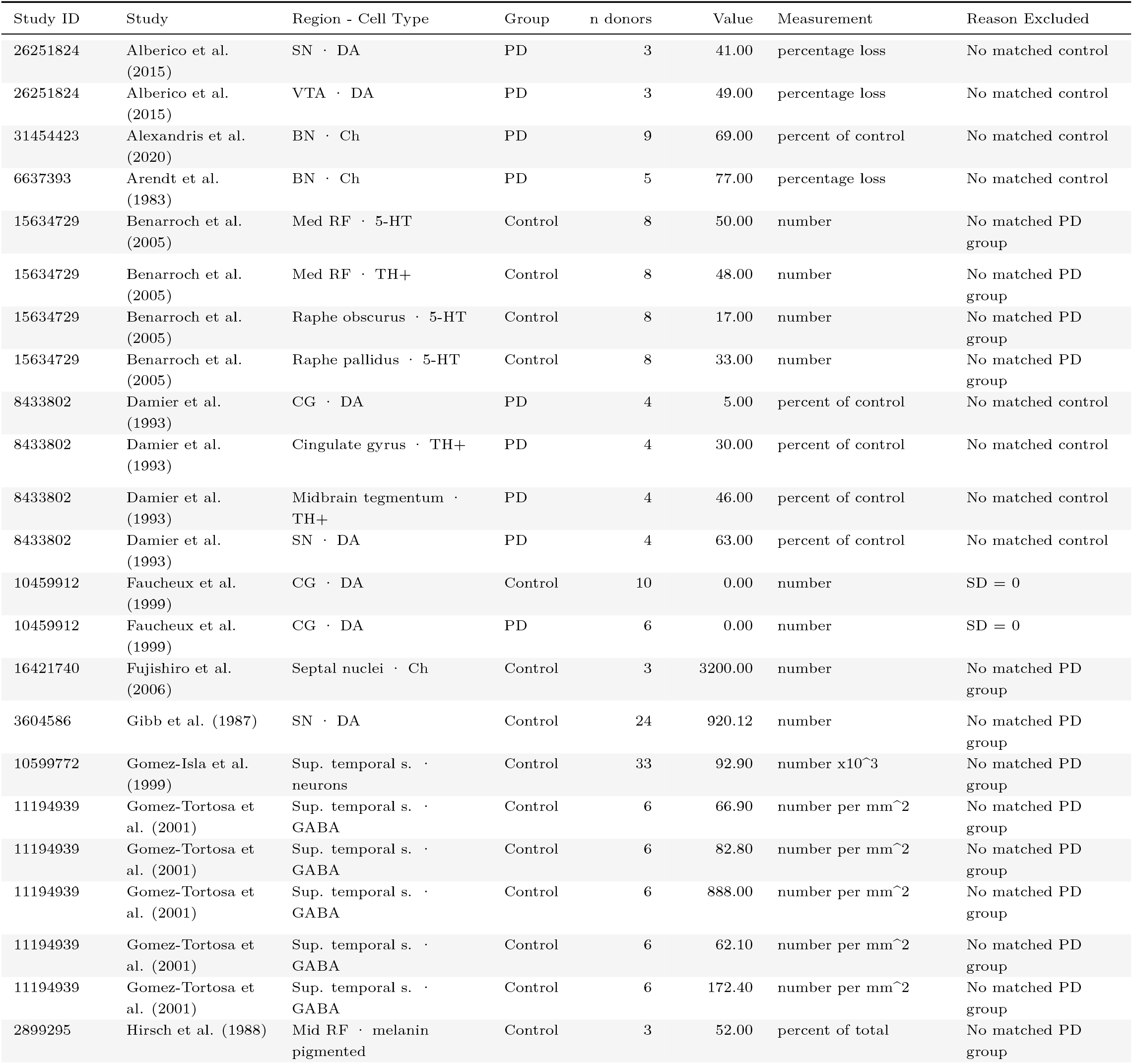
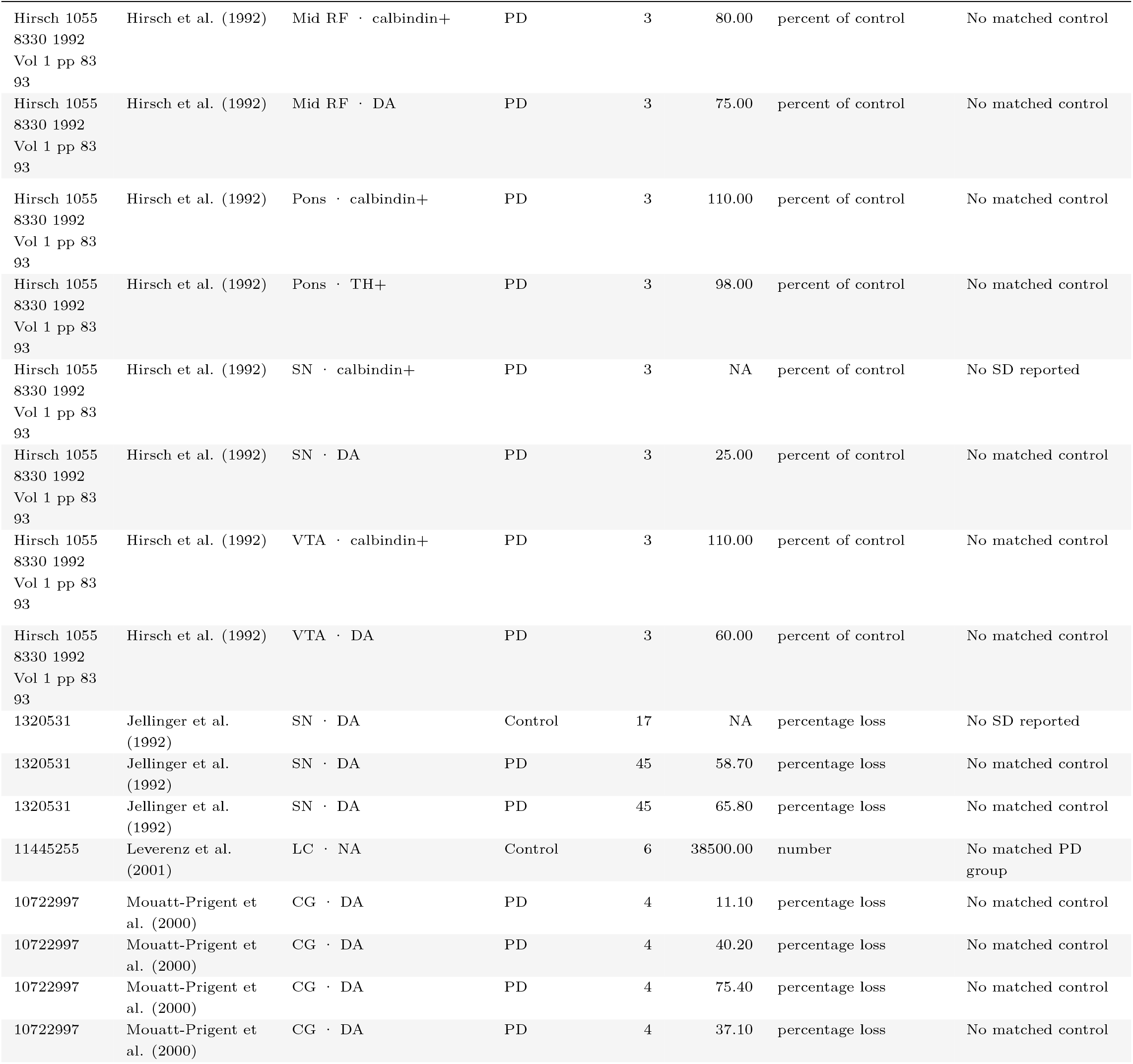
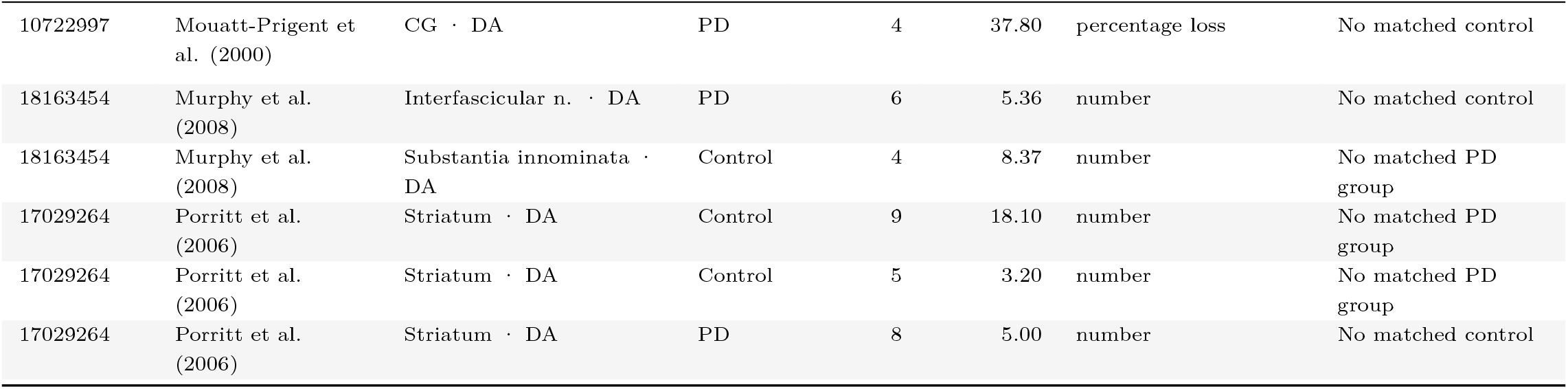
Observations omitted from effect-size calculation. Reasons include missing comparable PD and control measurements or unusable variability estimates. Values retain their reported units; n denotes donors in the recorded group.

| Study ID | Study | Region - Cell Type | Group | n donors | Value | Measurement | Reason Excluded |
| --- | --- | --- | --- | --- | --- | --- | --- |
| 26251824 | Alberico et al. (2015) | SN · DA | PD | 3 | 41.00 | percentage loss | No matched control |
| 26251824 | Alberico et al. (2015) | VTA · DA | PD | 3 | 49.00 | percentage loss | No matched control |
| 31454423 | Alexandris et al. (2020) | BN · Ch | PD | 9 | 69.00 | percent of control | No matched control |
| 6637393 | Arendt et al. (1983) | BN · Ch | PD | 5 | 77.00 | percentage loss | No matched control |
| 15634729 | Benarroch et al. (2005) | Med RF · 5-HT | Control | 8 | 50.00 | number | No matched PD group |
| 15634729 | Benarroch et al. (2005) | Med RF · TH+ | Control | 8 | 48.00 | number | No matched PD group |
| 15634729 | Benarroch et al. (2005) | Raphe obscurus · 5-HT | Control | 8 | 17.00 | number | No matched PD group |
| 15634729 | Benarroch et al. (2005) | Raphe pallidus · 5-HT | Control | 8 | 33.00 | number | No matched PD group |
| 8433802 | Damier et al. (1993) | CG · DA | PD | 4 | 5.00 | percent of control | No matched control |
| 8433802 | Damier et al. (1993) | Cingulate gyrus · TH+ | PD | 4 | 30.00 | percent of control | No matched control |
| 8433802 | Damier et al. (1993) | Midbrain tegmentum · TH+ | PD | 4 | 46.00 | percent of control | No matched control |
| 8433802 | Damier et al. (1993) | SN · DA | PD | 4 | 63.00 | percent of control | No matched control |
| 10459912 | Faucheux et al. (1999) | CG · DA | Control | 10 | 0.00 | number | SD = 0 |
| 10459912 | Faucheux et al. (1999) | CG · DA | PD | 6 | 0.00 | number | SD = 0 |
| 16421740 | Fujishiro et al. (2006) | Septal nuclei · Ch | Control | 3 | 3200.00 | number | No matched PD group |
| 3604586 | Gibb et al. (1987) | SN · DA | Control | 24 | 920.12 | number | No matched PD group |
| 10599772 | Gomez-Isla et al. (1999) | Sup. temporal s. · neurons | Control | 33 | 92.90 | number x10<sup>3</sup> | No matched PD group |
| 11194939 | Gomez-Tortosa et al. (2001) | Sup. temporal s. · GABA | Control | 6 | 66.90 | number per mm<sup>2</sup> | No matched PD group |
| 11194939 | Gomez-Tortosa et al. (2001) | Sup. temporal s. · GABA | Control | 6 | 82.80 | number per mm<sup>2</sup> | No matched PD group |
| 11194939 | Gomez-Tortosa et al. (2001) | Sup. temporal s. · GABA | Control | 6 | 888.00 | number per mm<sup>2</sup> | No matched PD group |
| 11194939 | Gomez-Tortosa et al. (2001) | Sup. temporal s. · GABA | Control | 6 | 62.10 | number per mm<sup>2</sup> | No matched PD group |
| 11194939 | Gomez-Tortosa et al. (2001) | Sup. temporal s. · GABA | Control | 6 | 172.40 | number per mm<sup>2</sup> | No matched PD group |
| 2899295 | Hirsch et al. (1988) | Mid RF · melanin pigmented | Control | 3 | 52.00 | percent of total | No matched PD group |
| Hirsch 1055<br>8330 1992<br>Vol 1 pp 83<br>93 | Hirsch et al. (1992) | Mid RF · calbindin+ | PD | 3 | 80.00 | percent of control | No matched control |
| Hirsch 1055<br>8330 1992<br>Vol 1 pp 83<br>93 | Hirsch et al. (1992) | Mid RF · DA | PD | 3 | 75.00 | percent of control | No matched control |
| Hirsch 1055<br>8330 1992<br>Vol 1 pp 83<br>93 | Hirsch et al. (1992) | Pons · calbindin+ | PD | 3 | 110.00 | percent of control | No matched control |
| Hirsch 1055<br>8330 1992<br>Vol 1 pp 83<br>93 | Hirsch et al. (1992) | Pons · TH+ | PD | 3 | 98.00 | percent of control | No matched control |
| Hirsch 1055<br>8330 1992<br>Vol 1 pp 83<br>93 | Hirsch et al. (1992) | SN · calbindin+ | PD | 3 | NA | percent of control | No SD reported |
| Hirsch 1055<br>8330 1992<br>Vol 1 pp 83<br>93 | Hirsch et al. (1992) | SN · DA | PD | 3 | 25.00 | percent of control | No matched control |
| Hirsch 1055<br>8330 1992<br>Vol 1 pp 83<br>93 | Hirsch et al. (1992) | VTA · calbindin+ | PD | 3 | 110.00 | percent of control | No matched control |
| Hirsch 1055<br>8330 1992<br>Vol 1 pp 83<br>93 | Hirsch et al. (1992) | VTA · DA | PD | 3 | 60.00 | percent of control | No matched control |
| 1320531 | Jellinger et al.<br>(1992) | SN · DA | Control | 17 | NA | percentage loss | No SD reported |
| 1320531 | Jellinger et al.<br>(1992) | SN · DA | PD | 45 | 58.70 | percentage loss | No matched control |
| 1320531 | Jellinger et al.<br>(1992) | SN · DA | PD | 45 | 65.80 | percentage loss | No matched control |
| 11445255 | Leverenz et al.<br>(2001) | LC · NA | Control | 6 | 38500.00 | number | No matched PD group |
| 10722997 | Mouatt-Prigent et al. (2000) | CG · DA | PD | 4 | 11.10 | percentage loss | No matched control |
| 10722997 | Mouatt-Prigent et al. (2000) | CG · DA | PD | 4 | 40.20 | percentage loss | No matched control |
| 10722997 | Mouatt-Prigent et al. (2000) | CG · DA | PD | 4 | 75.40 | percentage loss | No matched control |
| 10722997 | Mouatt-Prigent et al. (2000) | CG · DA | PD | 4 | 37.10 | percentage loss | No matched control |
| 10722997 | Mouatt-Prigent et al. (2000) | CG · DA | PD | 4 | 37.80 | percentage loss | No matched control |
| 18163454 | Murphy et al. (2008) | Interfascicular n. · DA | PD | 6 | 5.36 | number | No matched control |
| 18163454 | Murphy et al. (2008) | Substantia innominata · DA | Control | 4 | 8.37 | number | No matched PD group |
| 17029264 | Porritt et al. (2006) | Striatum · DA | Control | 9 | 18.10 | number | No matched PD group |
| 17029264 | Porritt et al. (2006) | Striatum · DA | Control | 5 | 3.20 | number | No matched PD group |
| 17029264 | Porritt et al. (2006) | Striatum · DA | PD | 8 | 5.00 | number | No matched control |

**Table 4:**
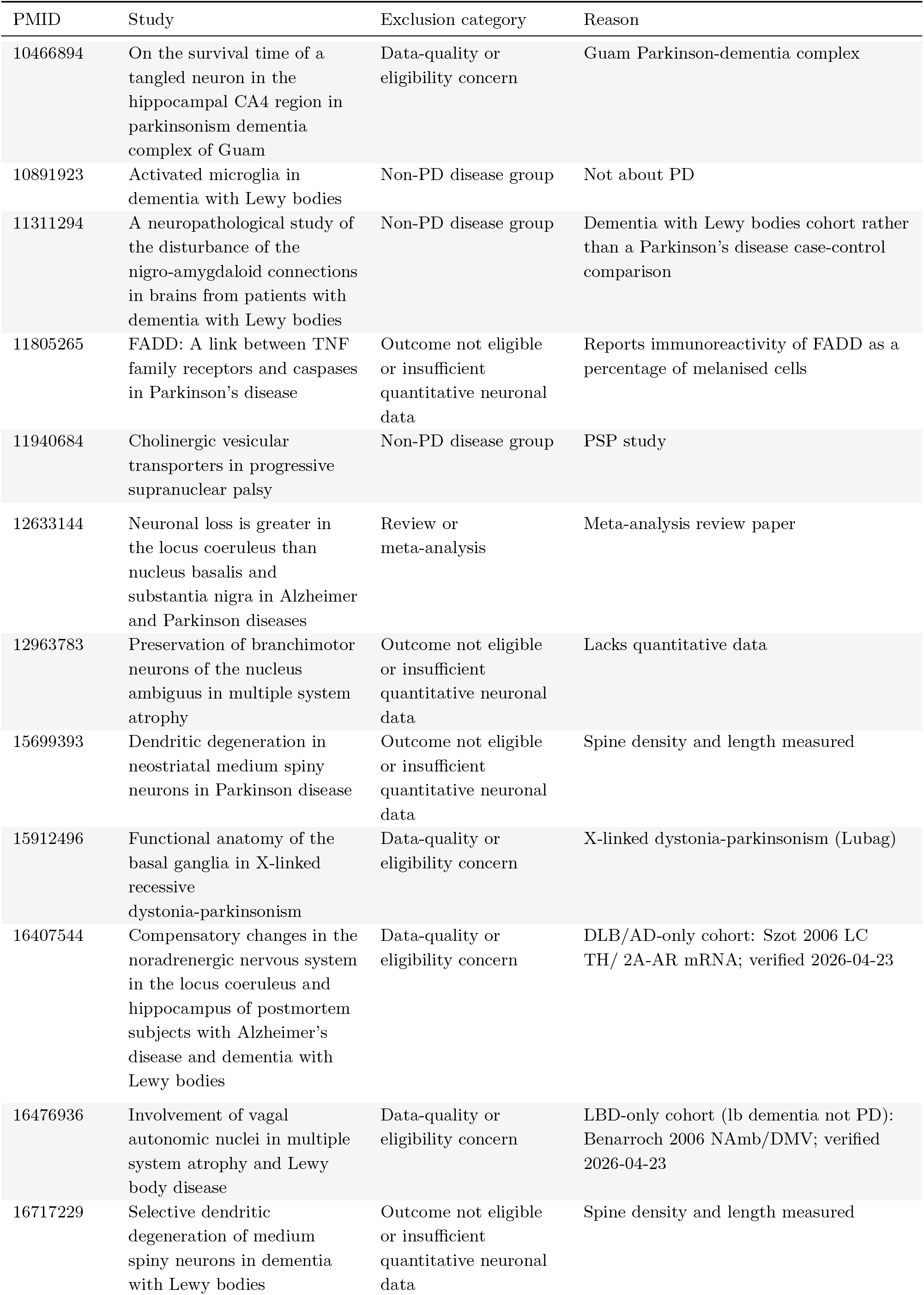
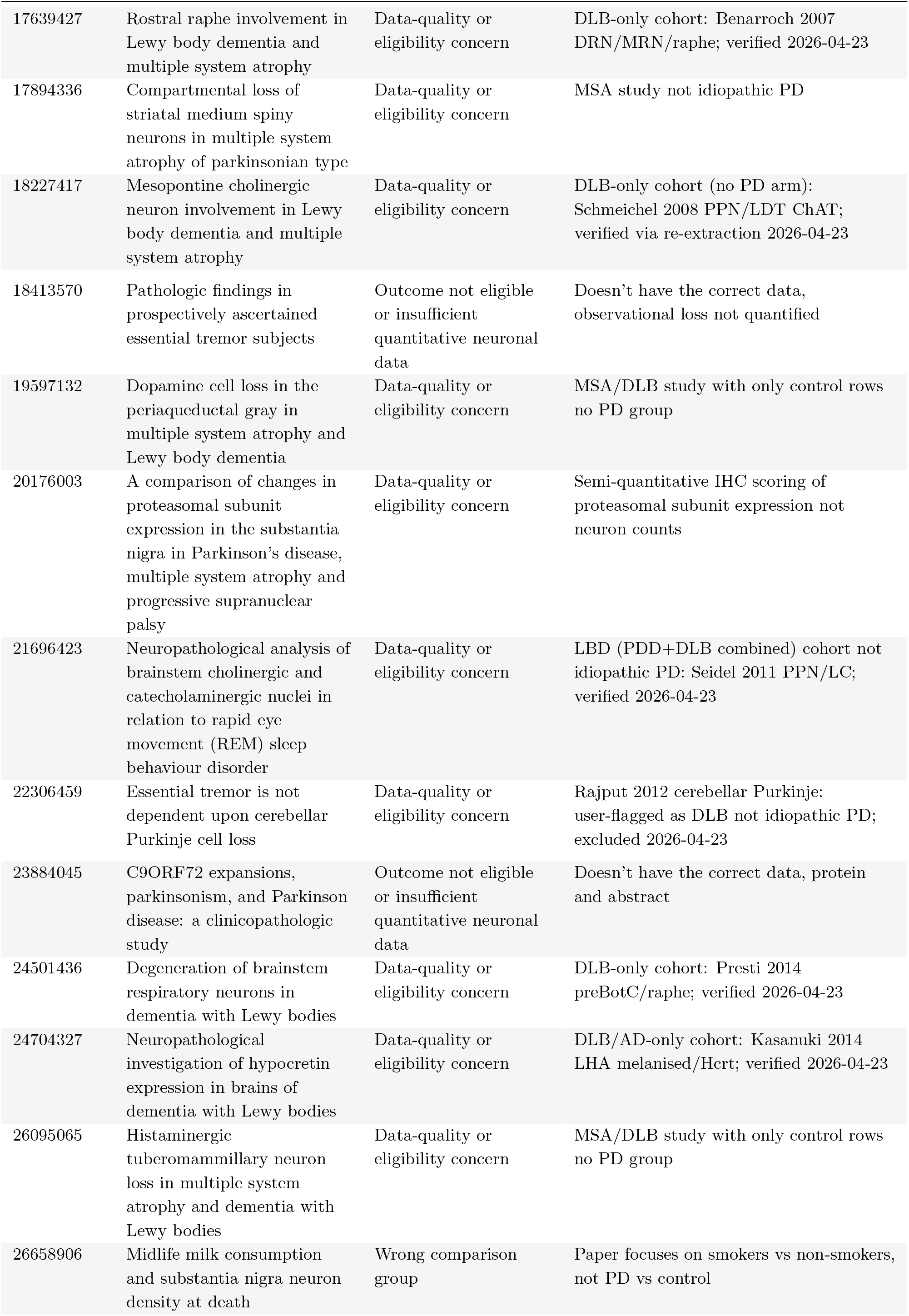
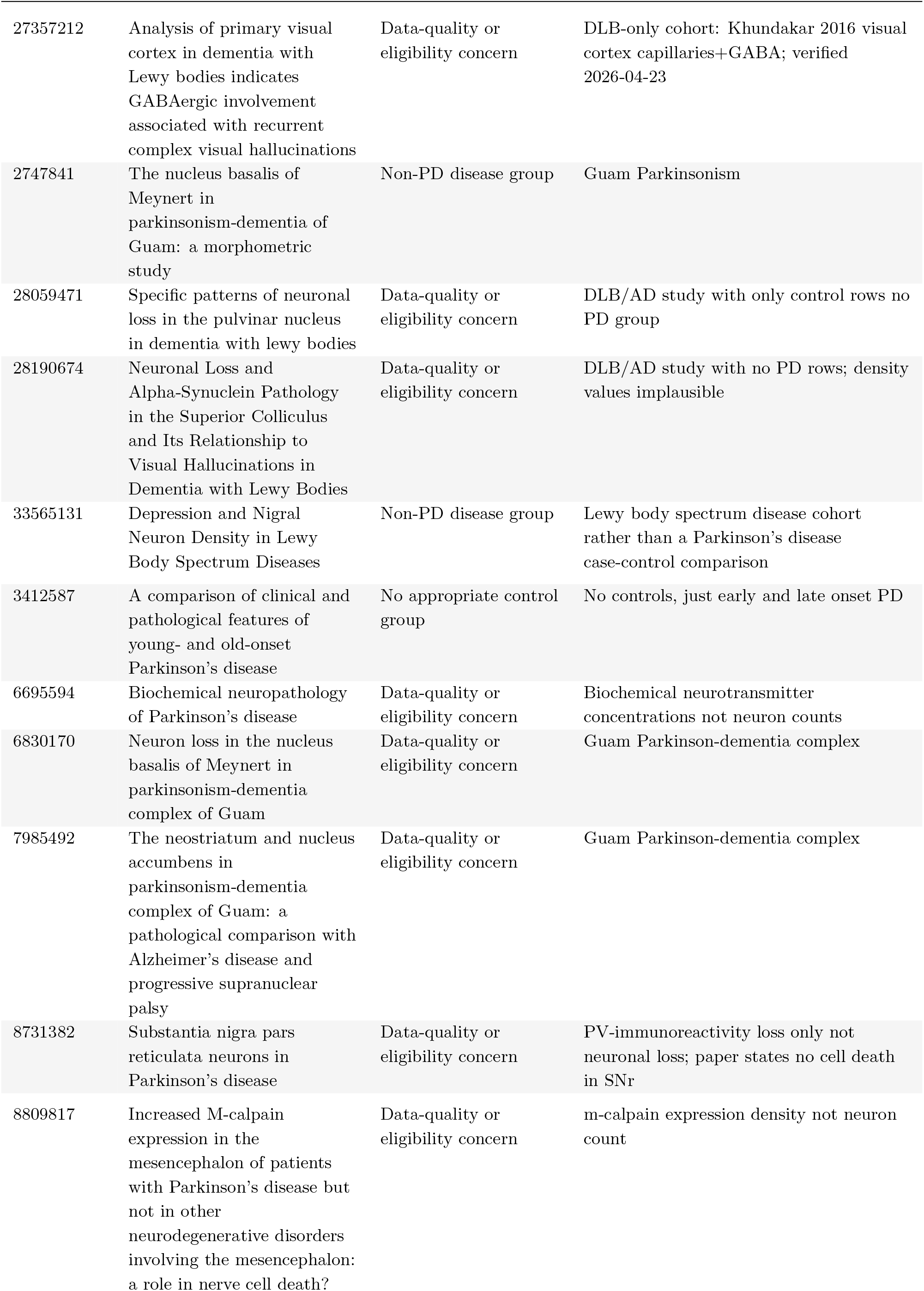
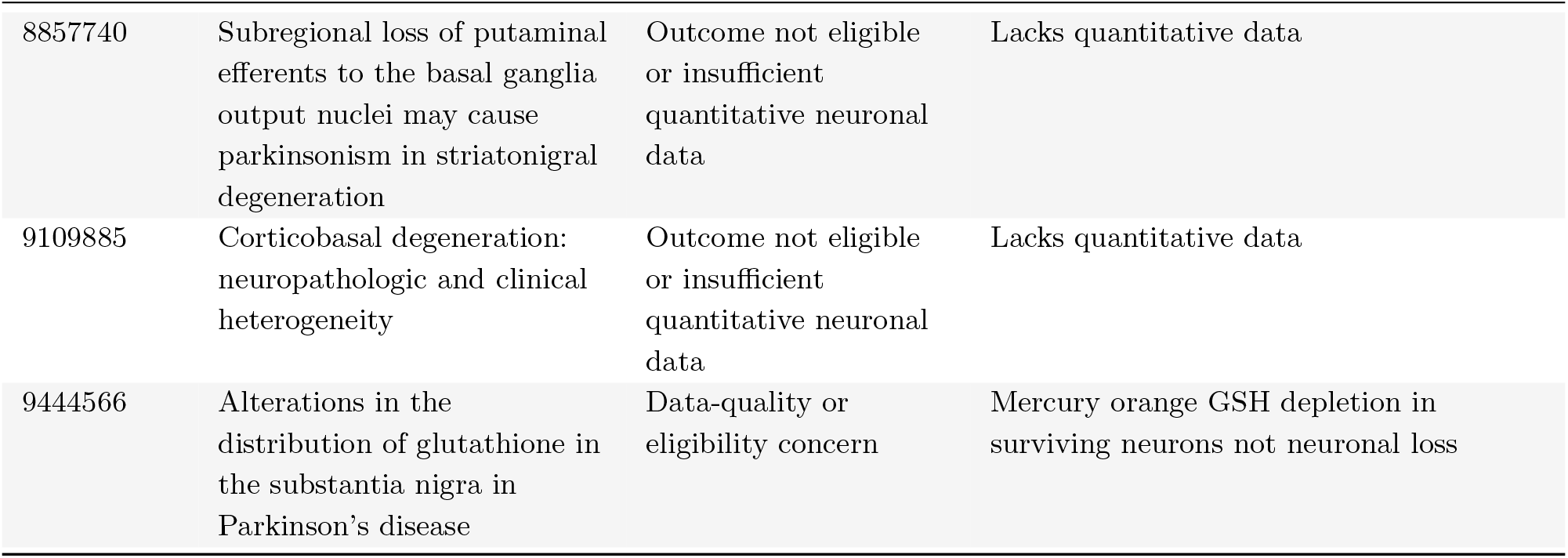
Studies excluded after initial full-text inclusion. Titles, exclusion categories and source-specific reasons are shown. Supplementary Table 5 records exclusions and corrections within studies.

| PMID | Study | Exclusion category | Reason |
| --- | --- | --- | --- |
| 10466894 | On the survival time of a tangled neuron in the hippocampal CA4 region in parkinsonism dementia complex of Guam | Data-quality or eligibility concern | Guam Parkinson-dementia complex |
| 10891923 | Activated microglia in dementia with Lewy bodies | Non-PD disease group | Not about PD |
| 11311294 | A neuropathological study of the disturbance of the nigro-amygdaloid connections in brains from patients with dementia with Lewy bodies | Non-PD disease group | Dementia with Lewy bodies cohort rather than a Parkinson's disease case-control comparison |
| 11805265 | FADD: A link between TNF family receptors and caspases in Parkinson's disease | Outcome not eligible or insufficient quantitative neuronal data | Reports immunoreactivity of FADD as a percentage of melanised cells |
| 11940684 | Cholinergic vesicular transporters in progressive supranuclear palsy | Non-PD disease group | PSP study |
| 12633144 | Neuronal loss is greater in the locus coeruleus than nucleus basalis and substantia nigra in Alzheimer and Parkinson diseases | Review or meta-analysis | Meta-analysis review paper |
| 12963783 | Preservation of branchiomotor neurons of the nucleus ambiguus in multiple system atrophy | Outcome not eligible or insufficient quantitative neuronal data | Lacks quantitative data |
| 15699393 | Dendritic degeneration in neostriatal medium spiny neurons in Parkinson disease | Outcome not eligible or insufficient quantitative neuronal data | Spine density and length measured |
| 15912496 | Functional anatomy of the basal ganglia in X-linked recessive dystonia-parkinsonism | Data-quality or eligibility concern | X-linked dystonia-parkinsonism (Lubag) |
| 16407544 | Compensatory changes in the noradrenergic nervous system in the locus coeruleus and hippocampus of postmortem subjects with Alzheimer's disease and dementia with Lewy bodies | Data-quality or eligibility concern | DLB/AD-only cohort: Szot 2006 LC TH/ 2A-AR mRNA; verified 2026-04-23 |
| 16476936 | Involvement of vagal autonomic nuclei in multiple system atrophy and Lewy body disease | Data-quality or eligibility concern | LBD-only cohort (lb dementia not PD): Benarroch 2006 NAmb/DMV; verified 2026-04-23 |
| 16717229 | Selective dendritic degeneration of medium spiny neurons in dementia with Lewy bodies | Outcome not eligible or insufficient quantitative neuronal data | Spine density and length measured |
| 17639427 | Rostral raphe involvement in Lewy body dementia and multiple system atrophy | Data-quality or eligibility concern | DLB-only cohort: Benarroch 2007 DRN/MRN/raphe; verified 2026-04-23 |
| 17894336 | Compartmental loss of striatal medium spiny neurons in multiple system atrophy of parkinsonian type | Data-quality or eligibility concern | MSA study not idiopathic PD |
| 18227417 | Mesopontine cholinergic neuron involvement in Lewy body dementia and multiple system atrophy | Data-quality or eligibility concern | DLB-only cohort (no PD arm): Schmeichel 2008 PPN/LDT ChAT; verified via re-extraction 2026-04-23 |
| 18413570 | Pathologic findings in prospectively ascertained essential tremor subjects | Outcome not eligible or insufficient quantitative neuronal data | Doesn't have the correct data, observational loss not quantified |
| 19597132 | Dopamine cell loss in the periaqueductal gray in multiple system atrophy and Lewy body dementia | Data-quality or eligibility concern | MSA/DLB study with only control rows no PD group |
| 20176003 | A comparison of changes in proteasomal subunit expression in the substantia nigra in Parkinson's disease, multiple system atrophy and progressive supranuclear palsy | Data-quality or eligibility concern | Semi-quantitative IHC scoring of proteasomal subunit expression not neuron counts |
| 21696423 | Neuropathological analysis of brainstem cholinergic and catecholaminergic nuclei in relation to rapid eye movement (REM) sleep behaviour disorder | Data-quality or eligibility concern | LBD (PDD+DLB combined) cohort not idiopathic PD: Seidel 2011 PPN/LC; verified 2026-04-23 |
| 22306459 | Essential tremor is not dependent upon cerebellar Purkinje cell loss | Data-quality or eligibility concern | Rajput 2012 cerebellar Purkinje: user-flagged as DLB not idiopathic PD; excluded 2026-04-23 |
| 23884045 | C9ORF72 expansions, parkinsonism, and Parkinson disease: a clinicopathologic study | Outcome not eligible or insufficient quantitative neuronal data | Doesn't have the correct data, protein and abstract |
| 24501436 | Degeneration of brainstem respiratory neurons in dementia with Lewy bodies | Data-quality or eligibility concern | DLB-only cohort: Presti 2014 preBotC/raphe; verified 2026-04-23 |
| 24704327 | Neuropathological investigation of hypocretin expression in brains of dementia with Lewy bodies | Data-quality or eligibility concern | DLB/AD-only cohort: Kasanuki 2014 LHA melanised/Hcrt; verified 2026-04-23 |
| 26095065 | Histaminergic tuberomammillary neuron loss in multiple system atrophy and dementia with Lewy bodies | Data-quality or eligibility concern | MSA/DLB study with only control rows no PD group |
| 26658906 | Midlife milk consumption and substantia nigra neuron density at death | Wrong comparison group | Paper focuses on smokers vs non-smokers, not PD vs control |
| 27357212 | Analysis of primary visual cortex in dementia with Lewy bodies indicates GABAergic involvement associated with recurrent complex visual hallucinations | Data-quality or eligibility concern | DLB-only cohort: Khundakar 2016 visual cortex capillaries+GABA; verified 2026-04-23 |
| 2747841 | The nucleus basalis of Meynert in parkinsonism-dementia of Guam: a morphometric study | Non-PD disease group | Guam Parkinsonism |
| 28059471 | Specific patterns of neuronal loss in the pulvinar nucleus in dementia with lewy bodies | Data-quality or eligibility concern | DLB/AD study with only control rows no PD group |
| 28190674 | Neuronal Loss and Alpha-Synuclein Pathology in the Superior Colliculus and Its Relationship to Visual Hallucinations in Dementia with Lewy Bodies | Data-quality or eligibility concern | DLB/AD study with no PD rows; density values implausible |
| 33565131 | Depression and Nigral Neuron Density in Lewy Body Spectrum Diseases | Non-PD disease group | Lewy body spectrum disease cohort rather than a Parkinson's disease case-control comparison |
| 3412587 | A comparison of clinical and pathological features of young- and old-onset Parkinson's disease | No appropriate control group | No controls, just early and late onset PD |
| 6695594 | Biochemical neuropathology of Parkinson's disease | Data-quality or eligibility concern | Biochemical neurotransmitter concentrations not neuron counts |
| 6830170 | Neuron loss in the nucleus basalis of Meynert in parkinsonism-dementia complex of Guam | Data-quality or eligibility concern | Guam Parkinson-dementia complex |
| 7985492 | The neostriatum and nucleus accumbens in parkinsonism-dementia complex of Guam: a pathological comparison with Alzheimer's disease and progressive supranuclear palsy | Data-quality or eligibility concern | Guam Parkinson-dementia complex |
| 8731382 | Substantia nigra pars reticulata neurons in Parkinson's disease | Data-quality or eligibility concern | PV-immunoreactivity loss only not neuronal loss; paper states no cell death in SNr |
| 8809817 | Increased M-calpain expression in the mesencephalon of patients with Parkinson's disease but not in other neurodegenerative disorders involving the mesencephalon: a role in nerve cell death? | Data-quality or eligibility concern | m-calpain expression density not neuron count |
| 8857740 | Subregional loss of putaminal efferents to the basal ganglia output nuclei may cause parkinsonism in striatonigral degeneration | Outcome not eligible or insufficient quantitative neuronal data | Lacks quantitative data |
| 9109885 | Corticobasal degeneration: neuropathologic and clinical heterogeneity | Outcome not eligible or insufficient quantitative neuronal data | Lacks quantitative data |
| 9444566 | Alterations in the distribution of glutathione in the substantia nigra in Parkinson's disease | Data-quality or eligibility concern | Mercury orange GSH depletion in surviving neurons not neuronal loss |

**Table 5:**
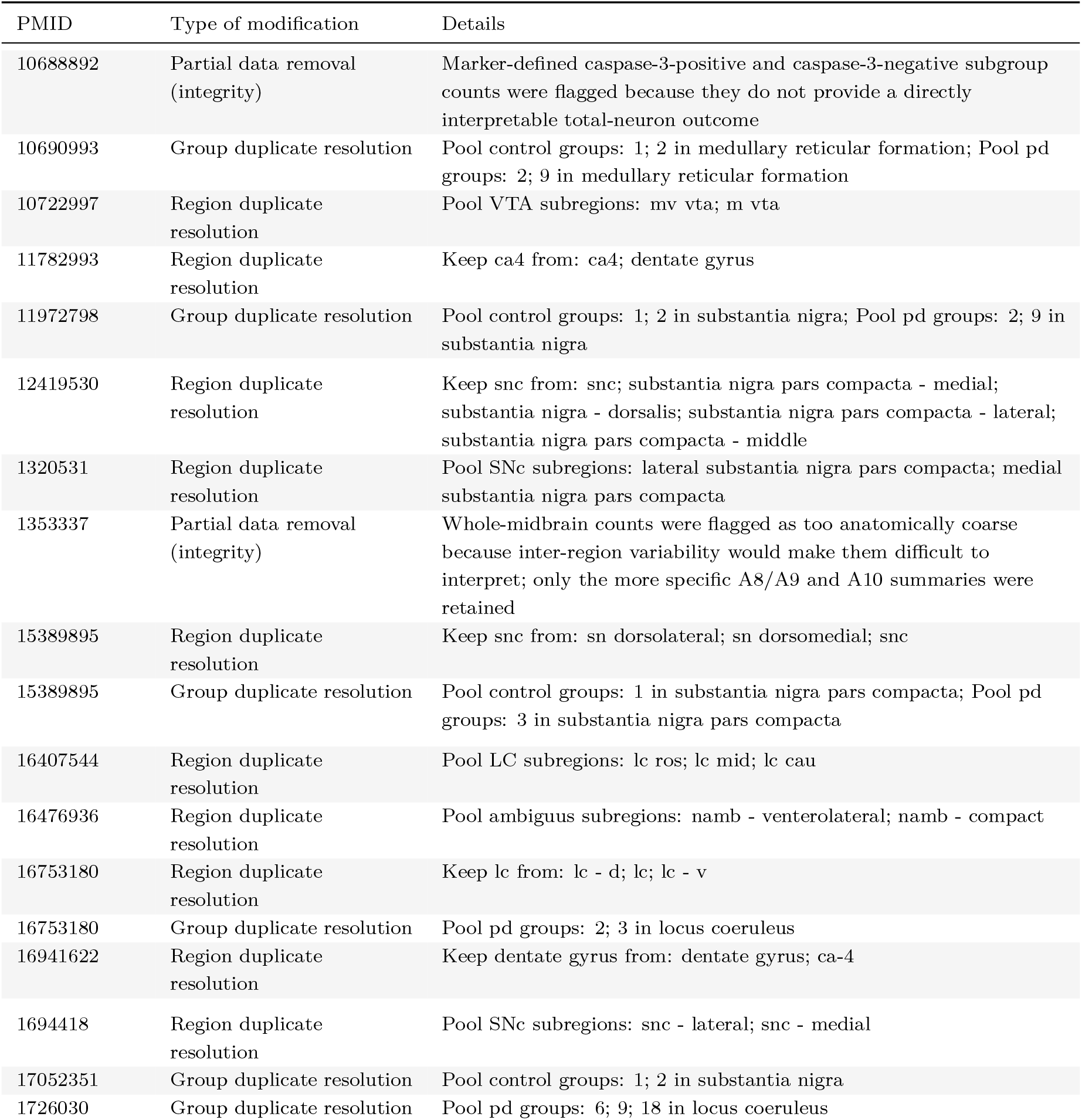
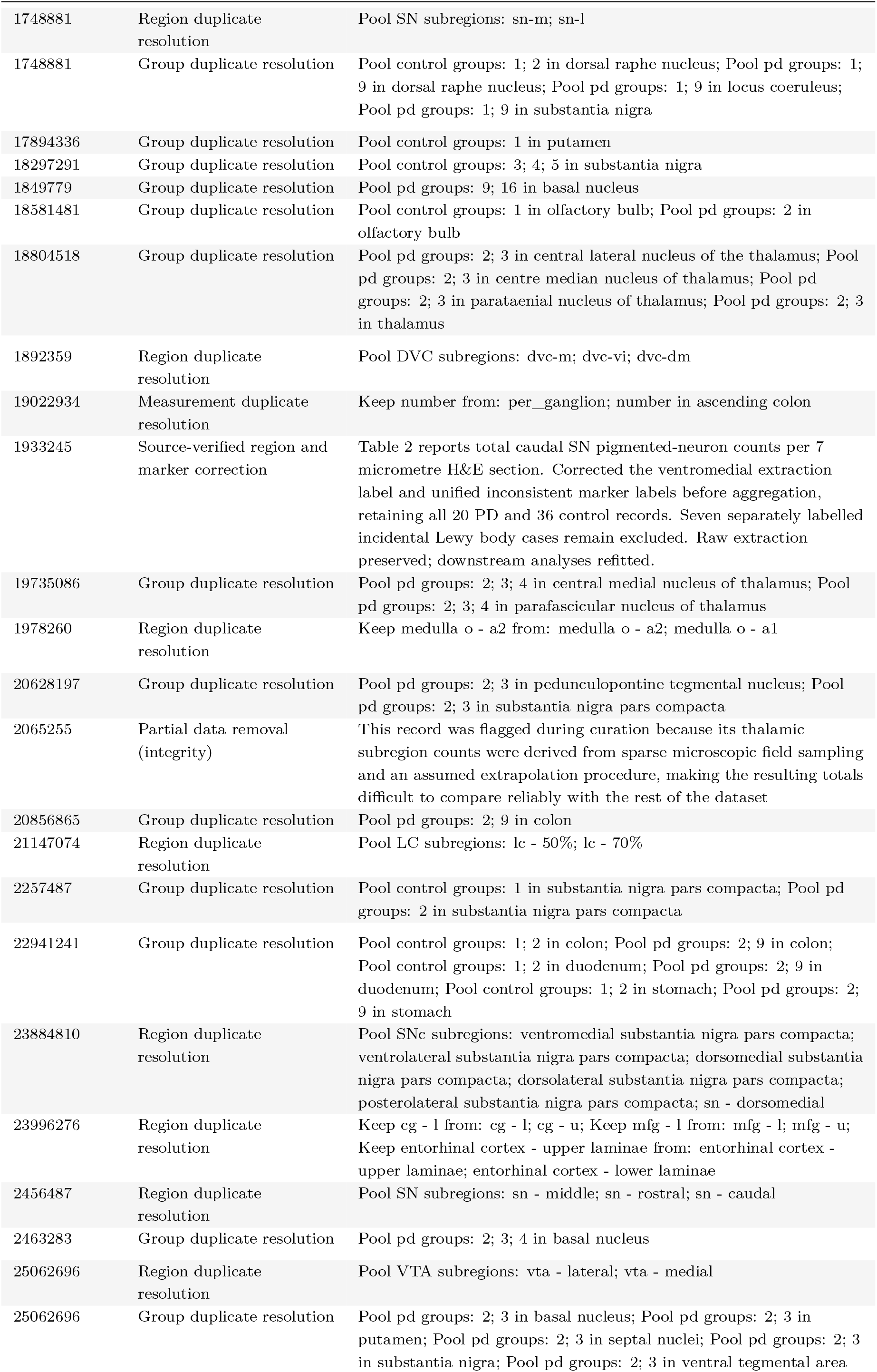
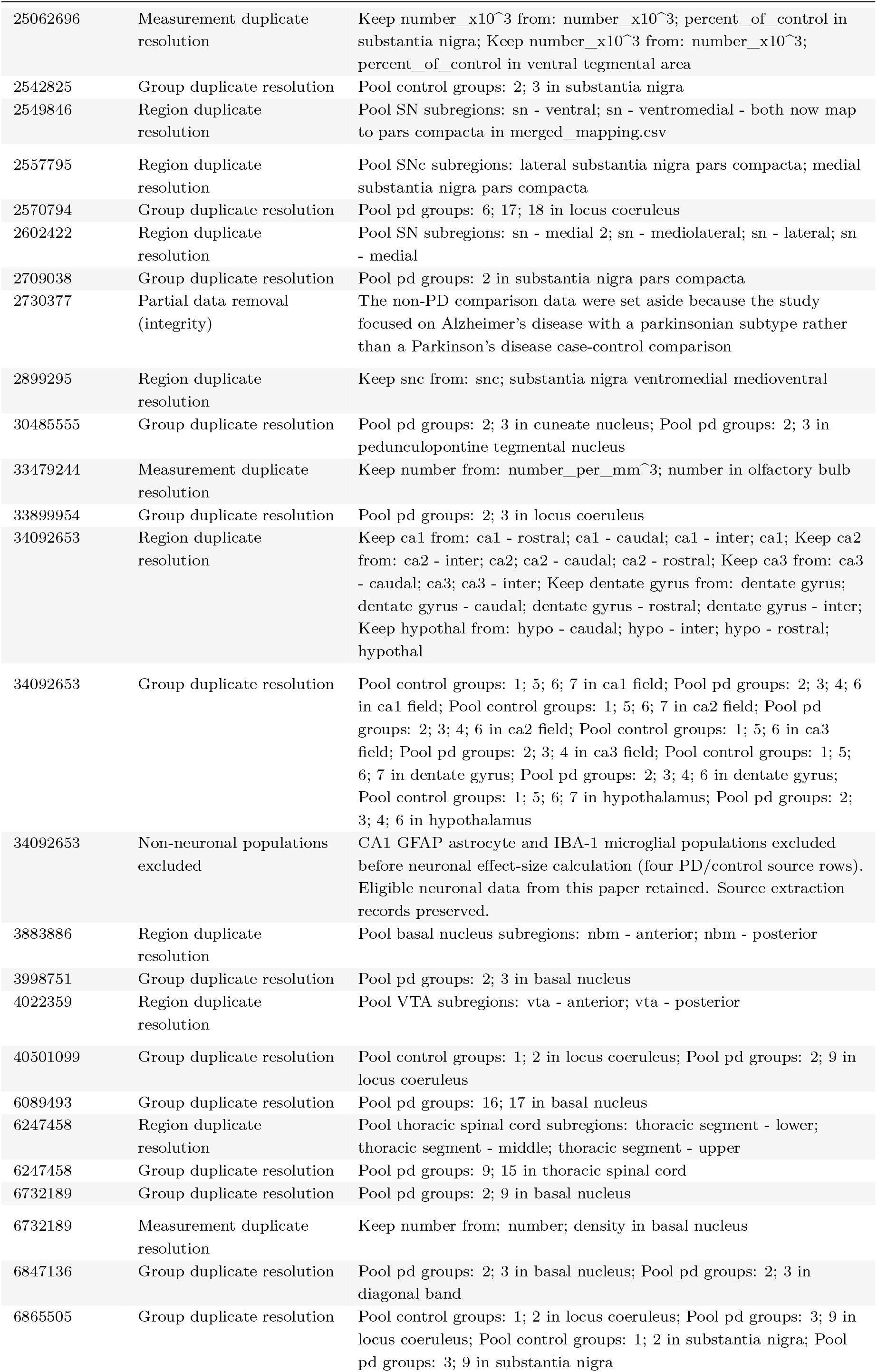
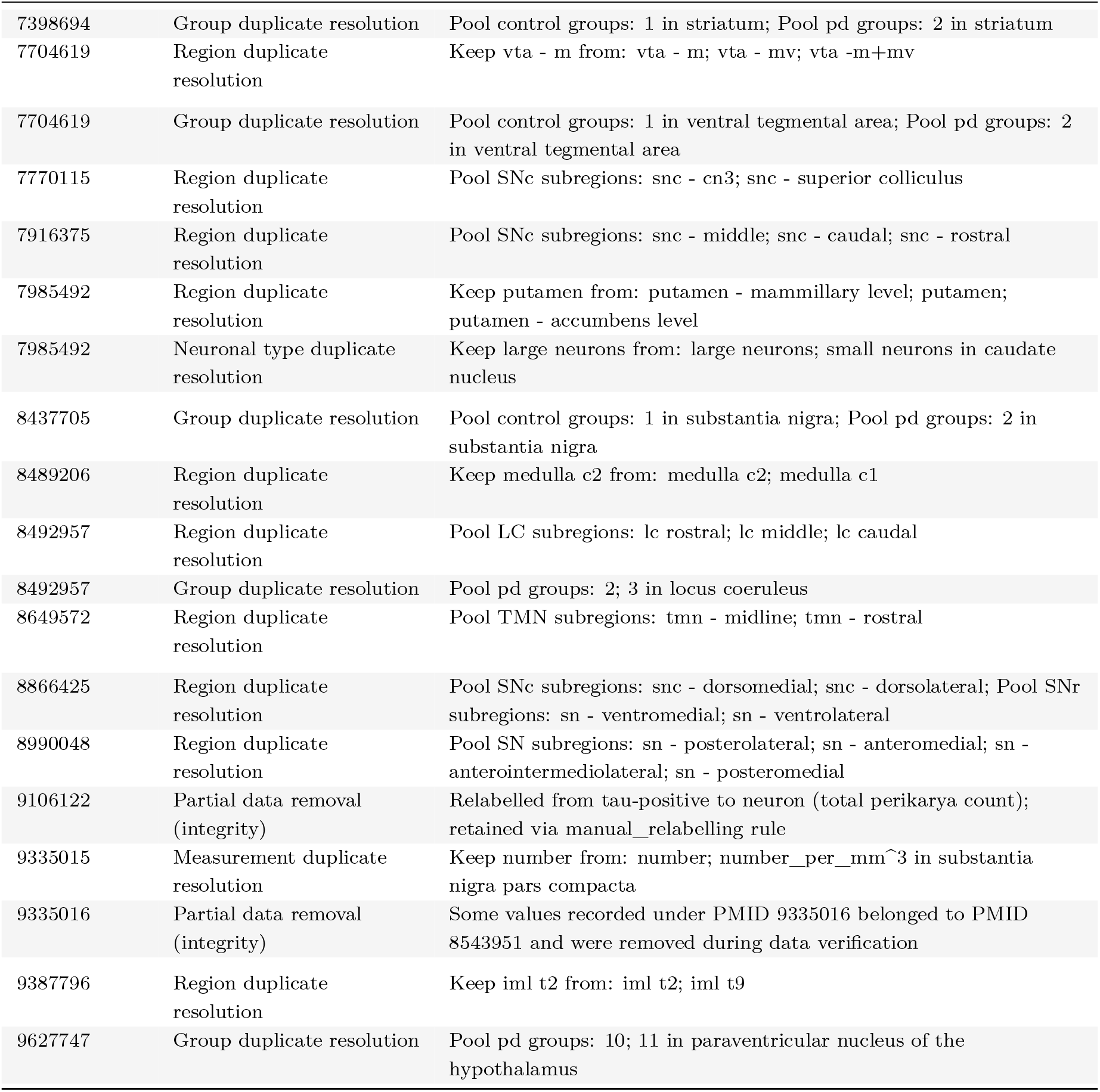
Curation after initial inclusion. The table records partial data exclusions, corrections and decisions about duplicate or overlapping measurements. It documents processing decisions rather than providing another list of eligible studies.

**Table 6:**
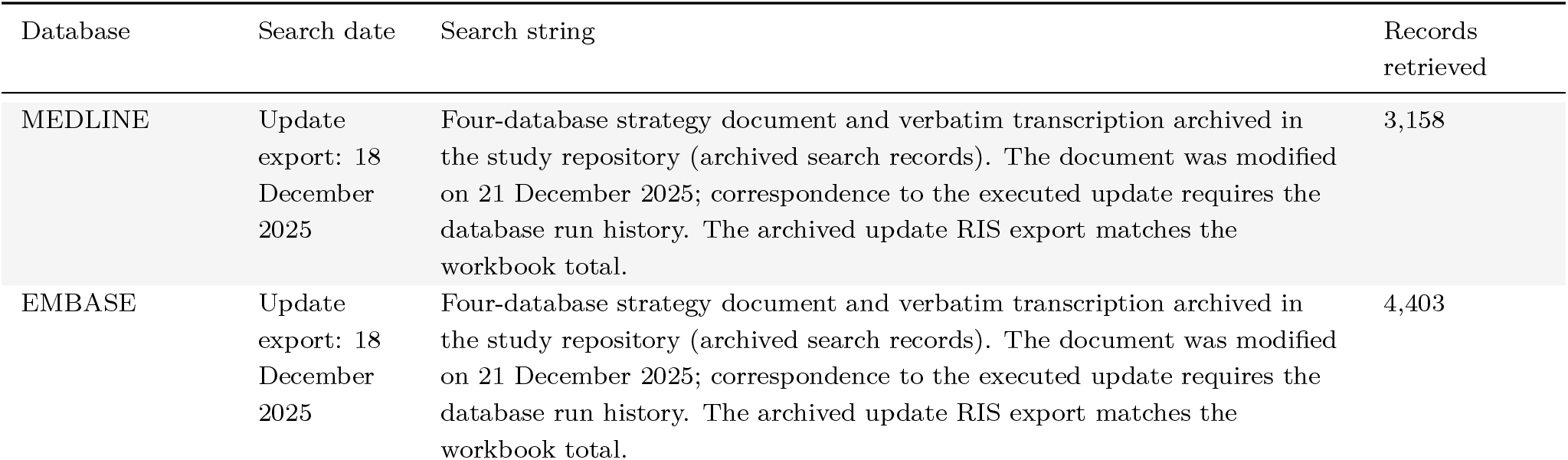
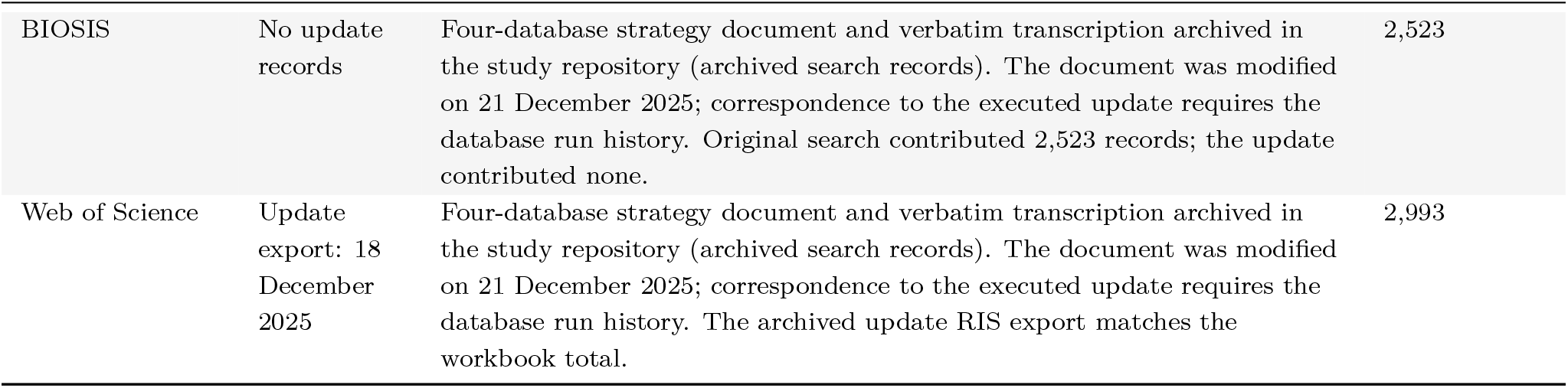
Database search audit. Counts combine the original search and the 2021-2025 update recorded in the screening workbook. The strategy document, verbatim transcription and update exports are archived with the data. Export dates are documented, but the exact executed search history remains unverified.

| Database | Search date | Search string | Records retrieved |
| --- | --- | --- | --- |
| MEDLINE | Update export: 18 December 2025 | Four-database strategy document and verbatim transcription archived in the study repository (archived search records). The document was modified on 21 December 2025; correspondence to the executed update requires the database run history. The archived update RIS export matches the workbook total. | 3,158 |
| EMBASE | Update export: 18 December 2025 | Four-database strategy document and verbatim transcription archived in the study repository (archived search records). The document was modified on 21 December 2025; correspondence to the executed update requires the database run history. The archived update RIS export matches the workbook total. | 4,403 |
| BIOSIS | No update records | Four-database strategy document and verbatim transcription archived in the study repository (archived search records). The document was modified on 21 December 2025; correspondence to the executed update requires the database run history. Original search contributed 2,523 records; the update contributed none. | 2,523 |
| Web of Science | Update export: 18 December 2025 | Four-database strategy document and verbatim transcription archived in the study repository (archived search records). The document was modified on 21 December 2025; correspondence to the executed update requires the database run history. The archived update RIS export matches the workbook total. | 2,993 |

**Table 7:**
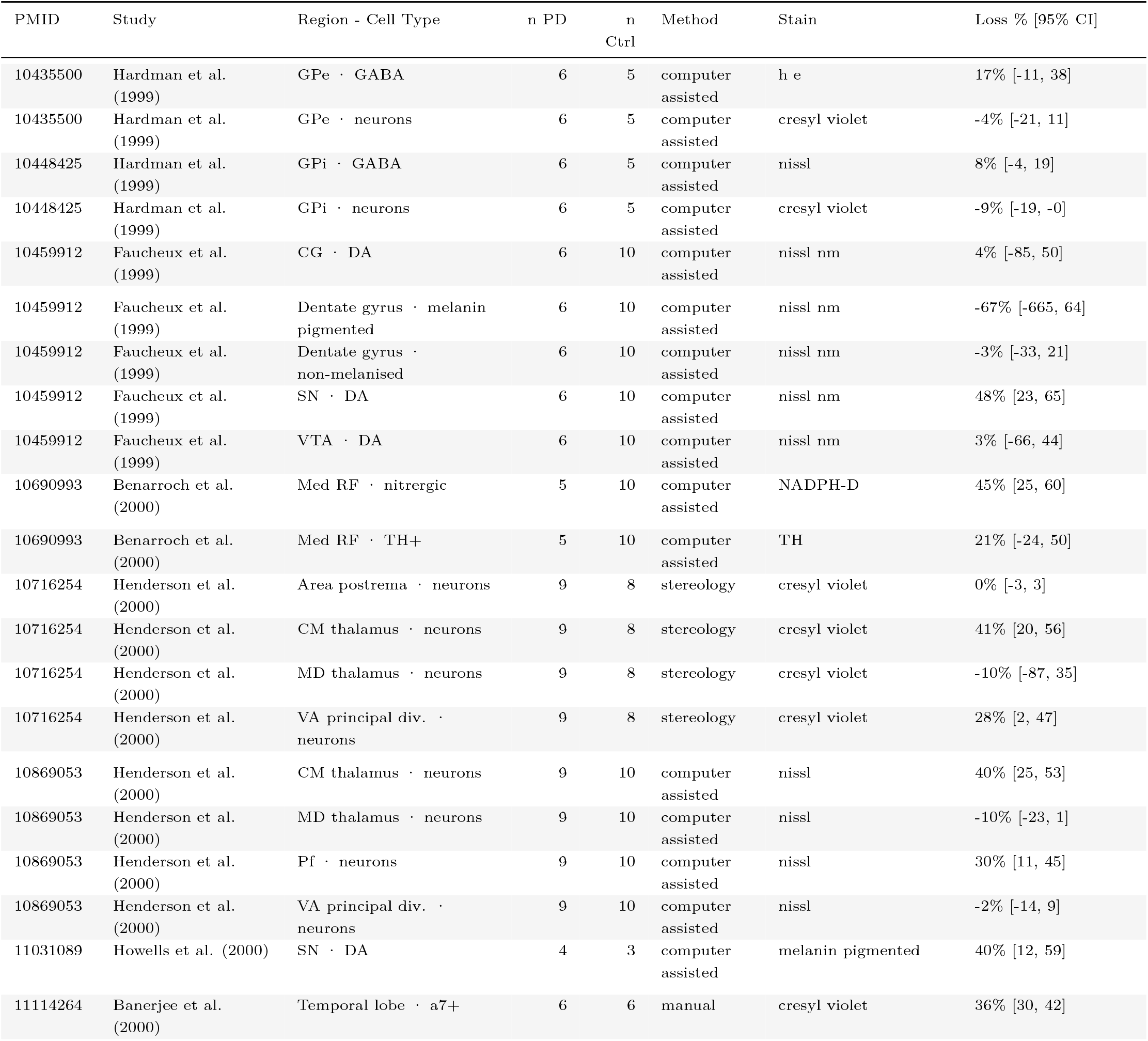
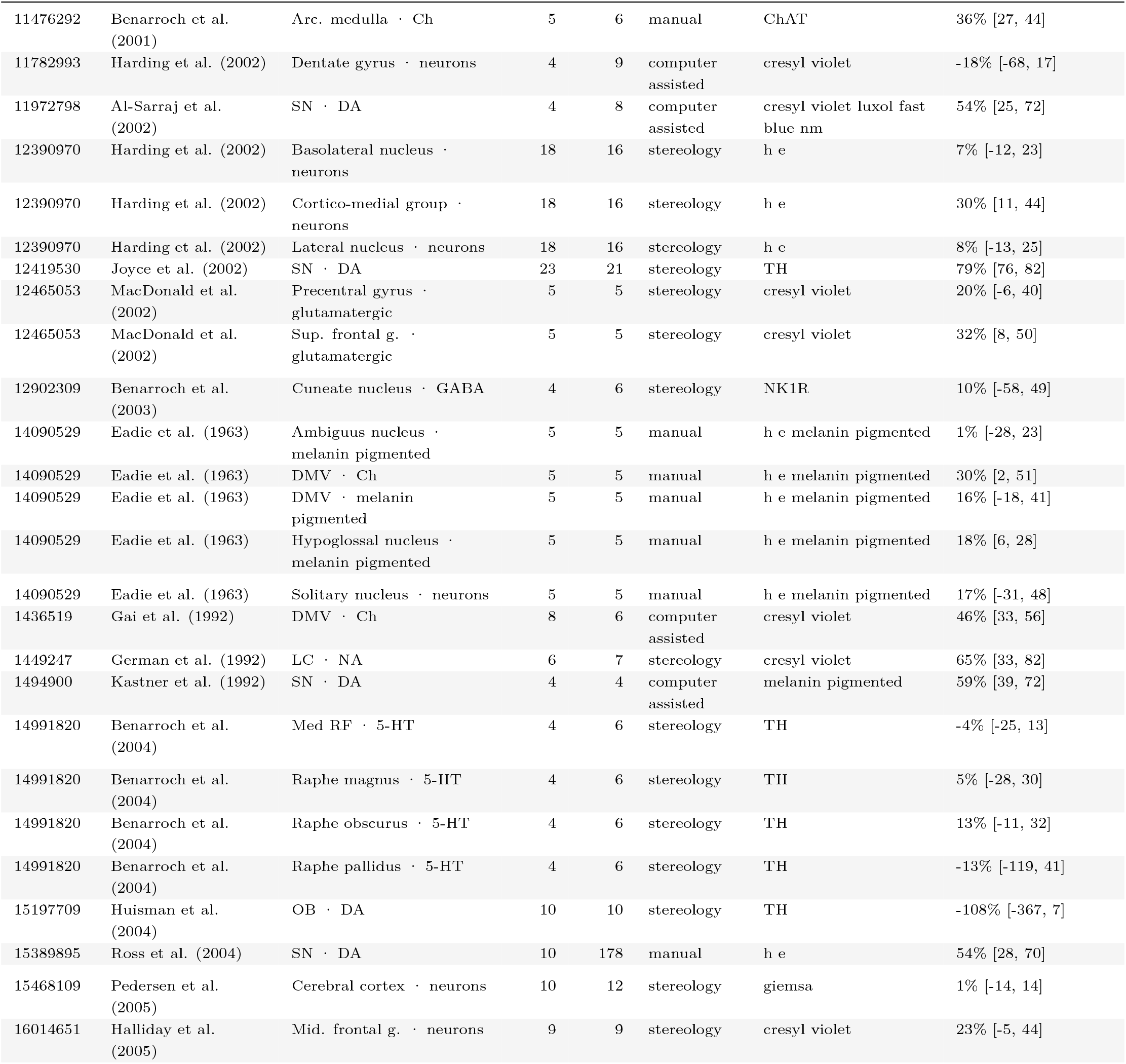
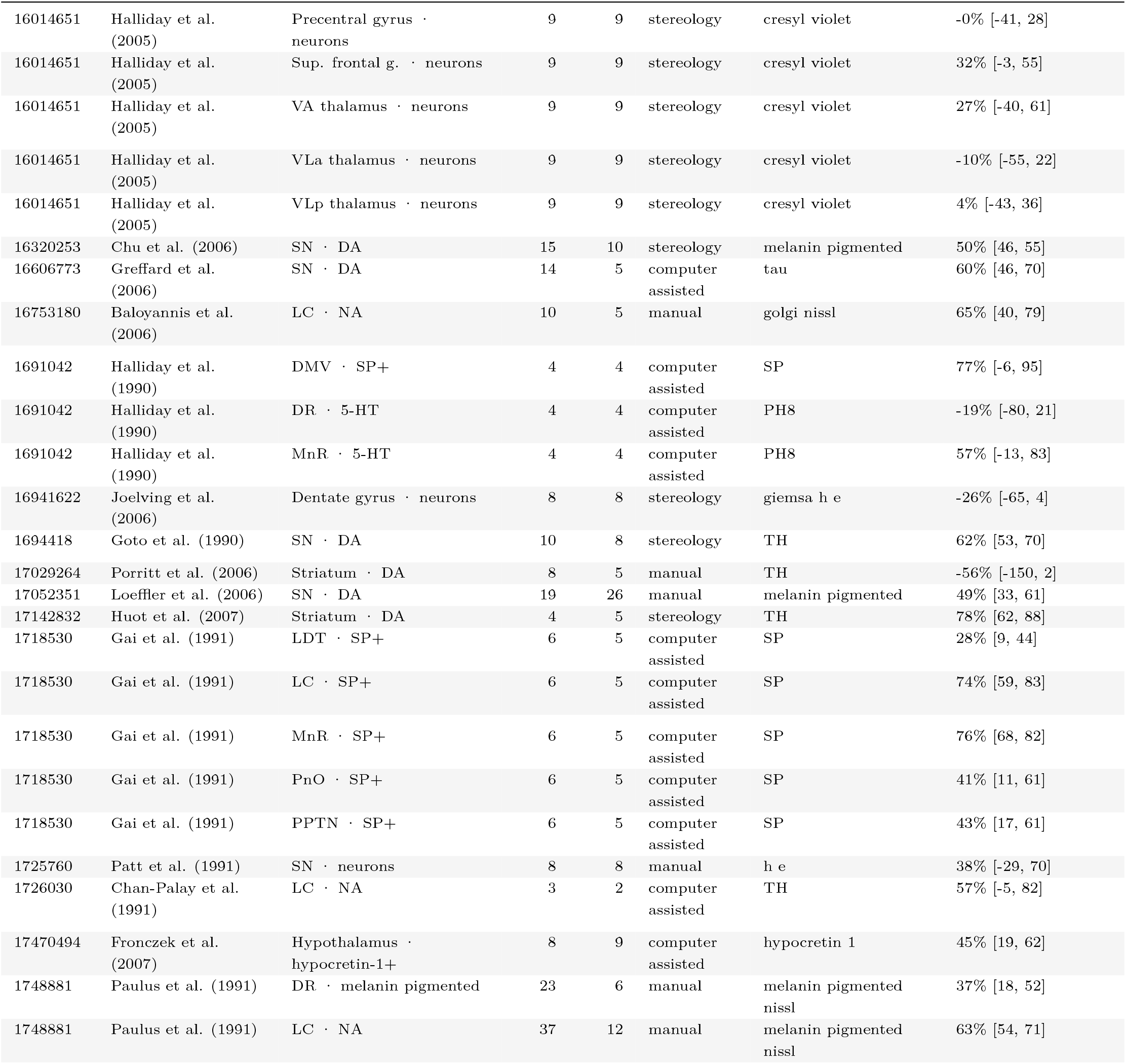
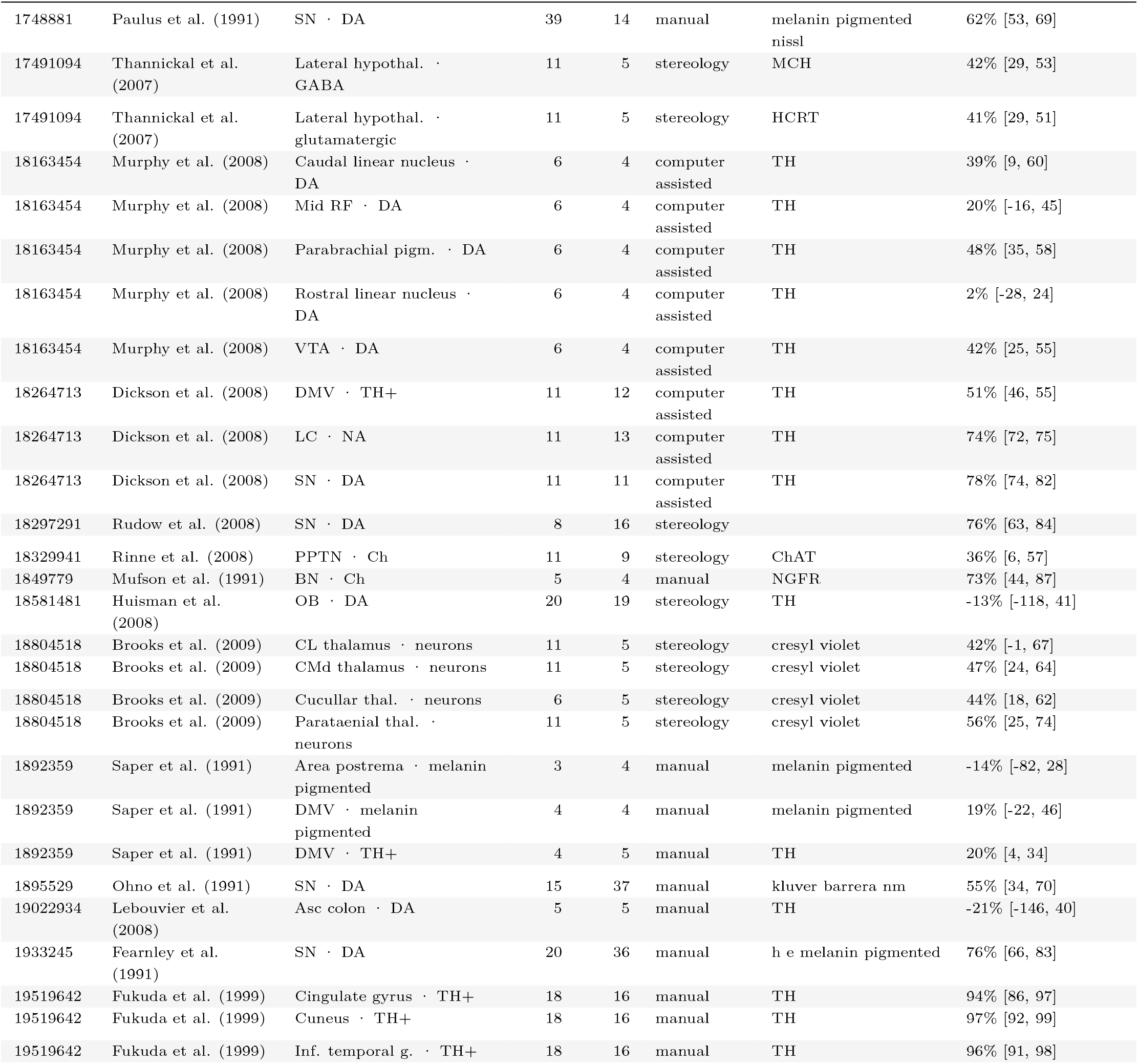
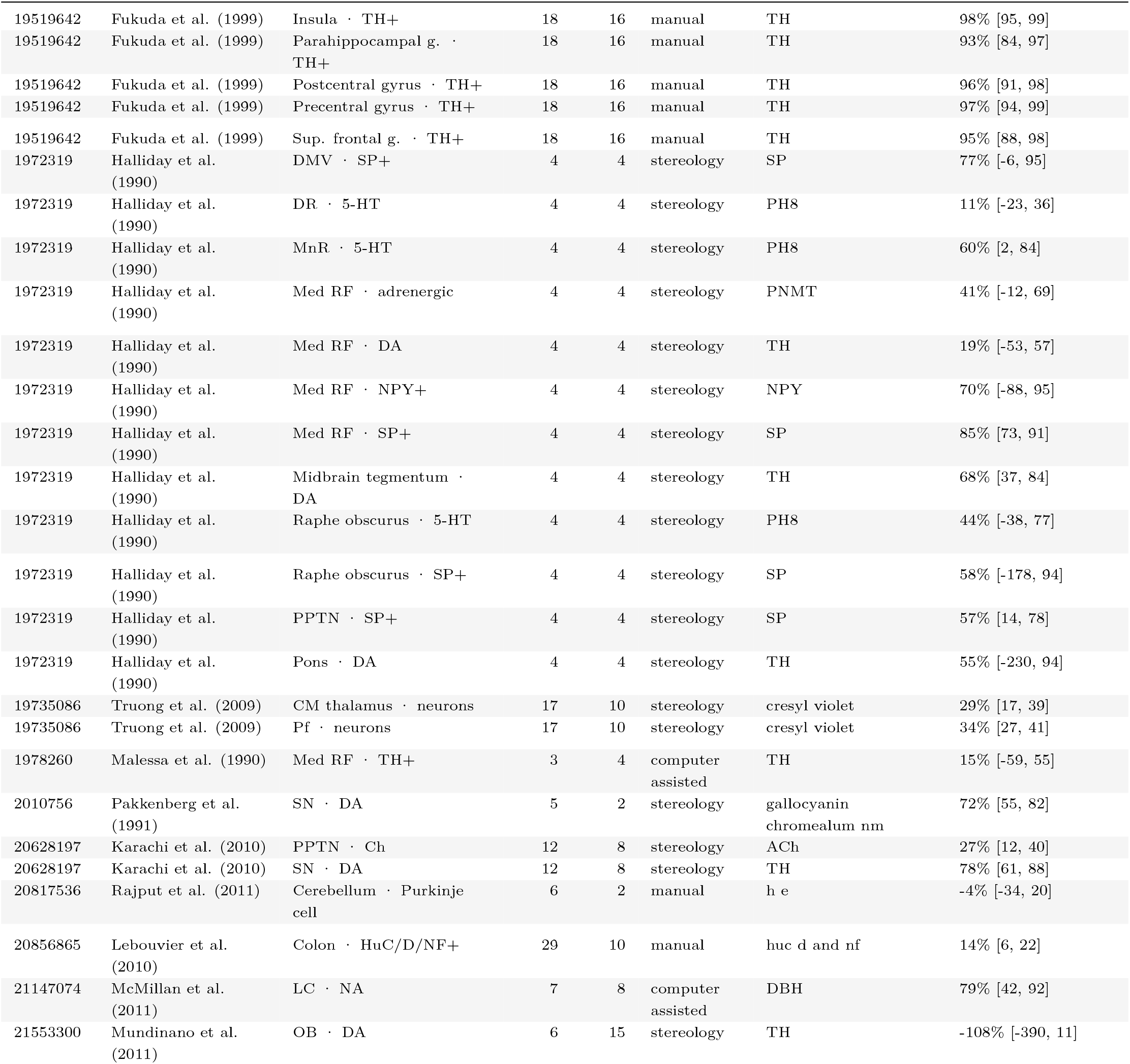
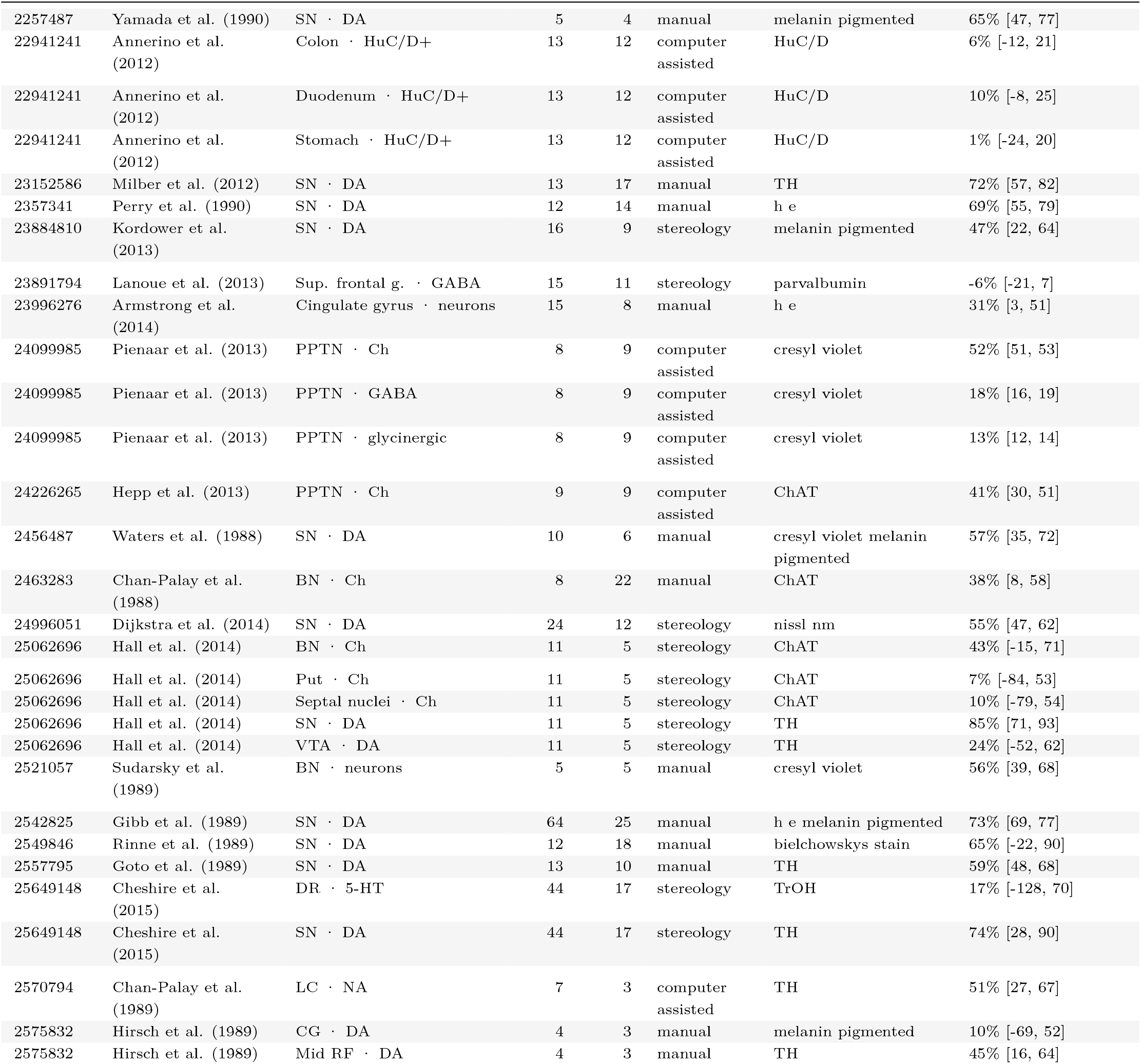
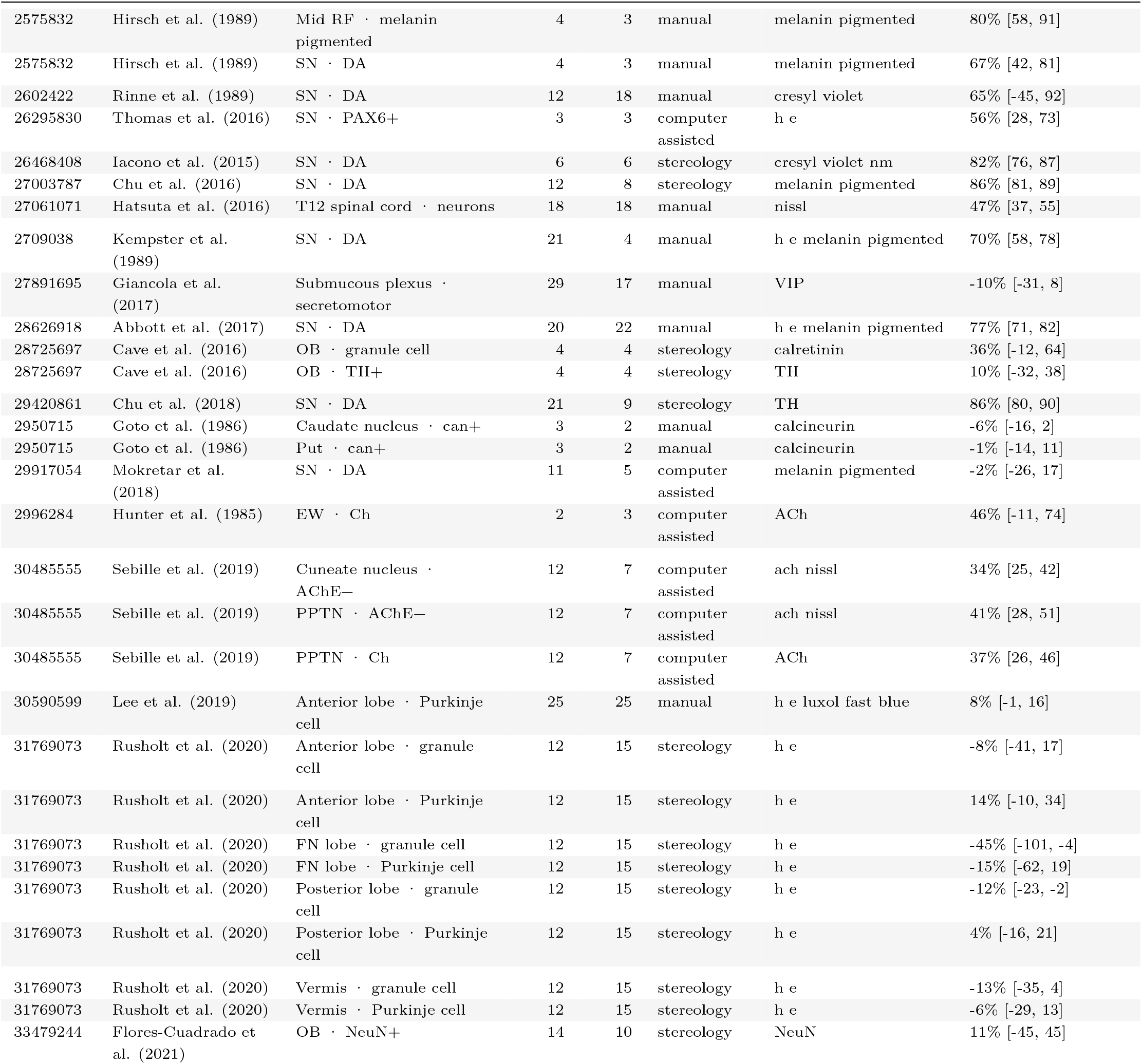
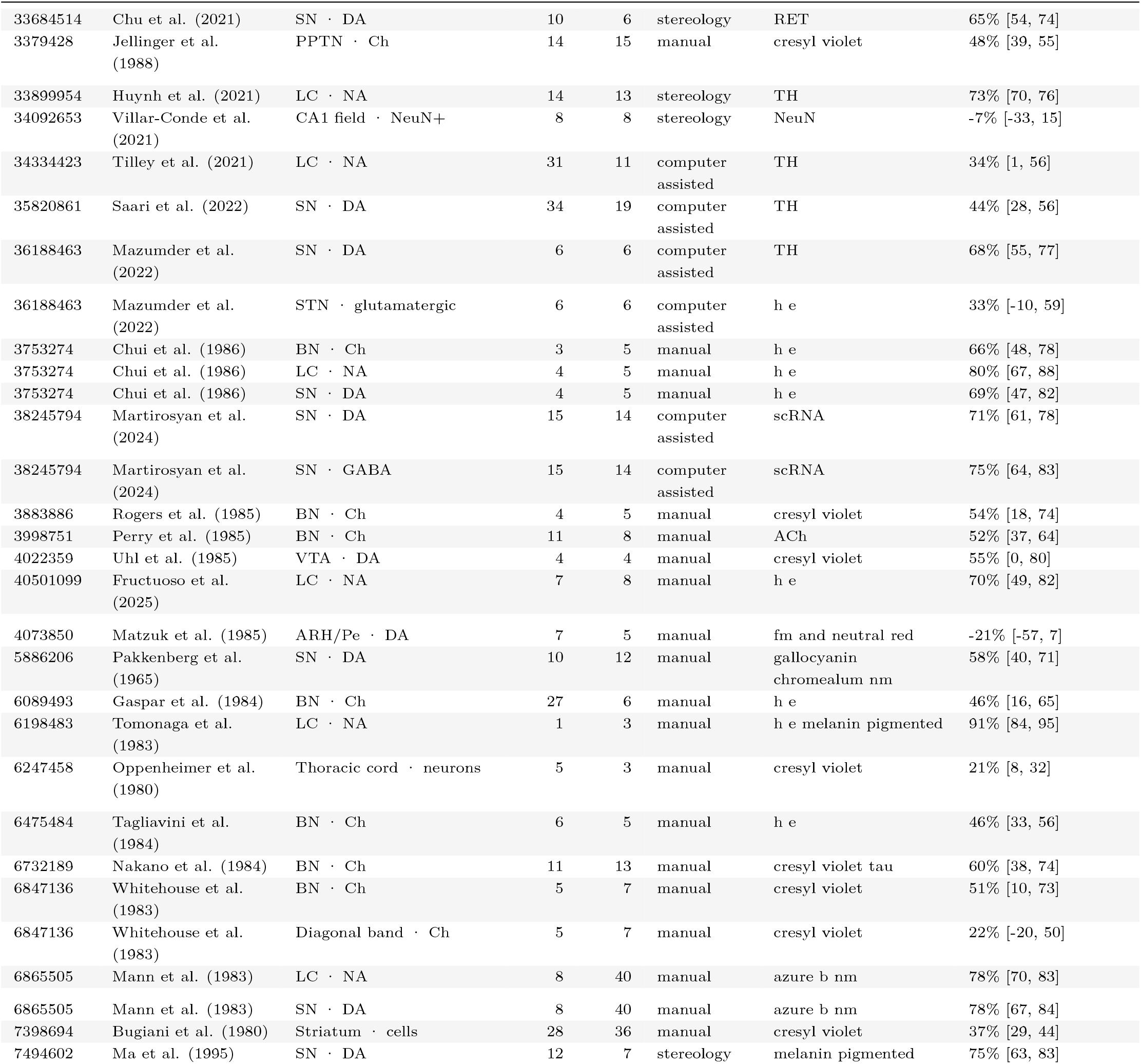
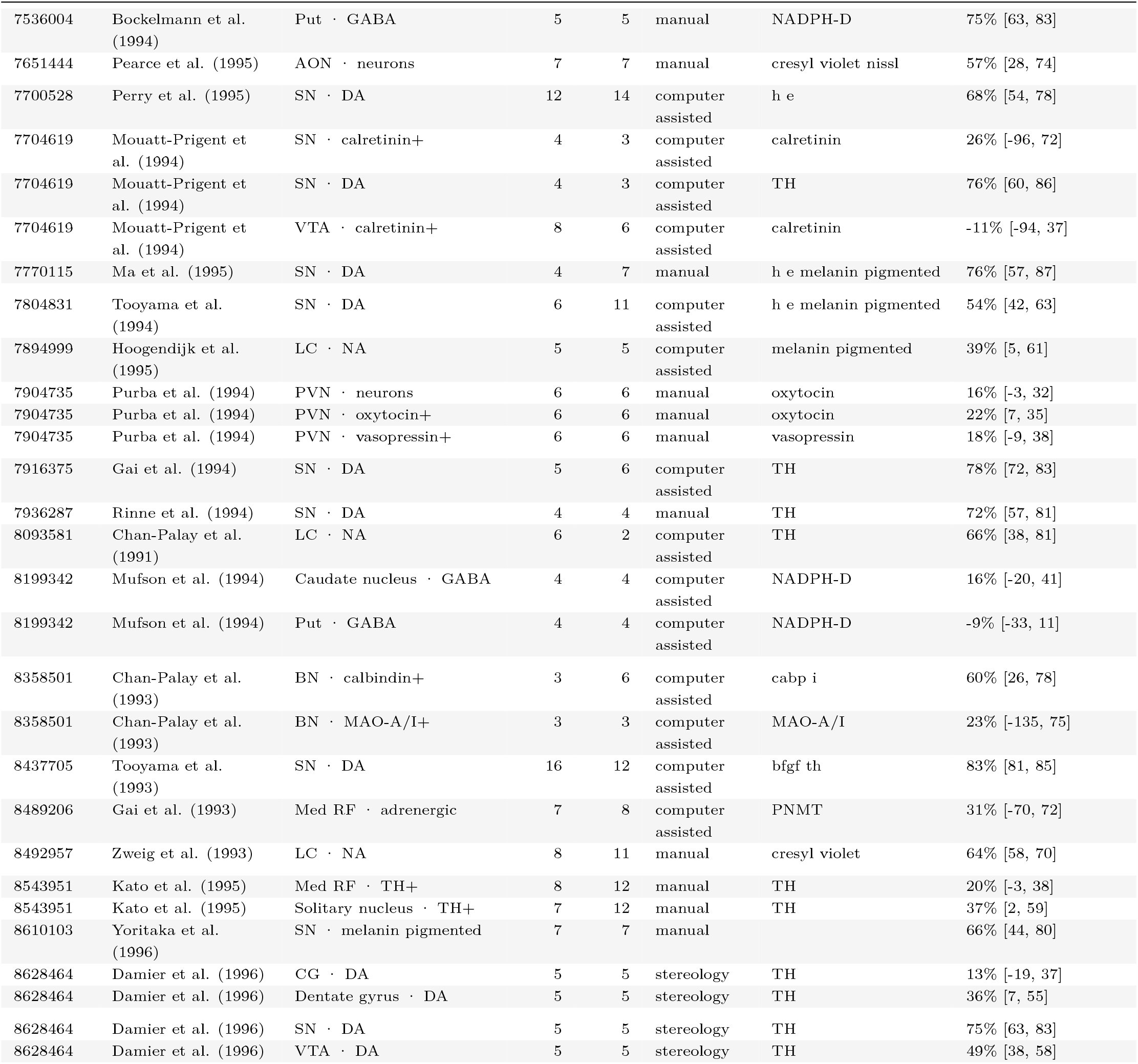
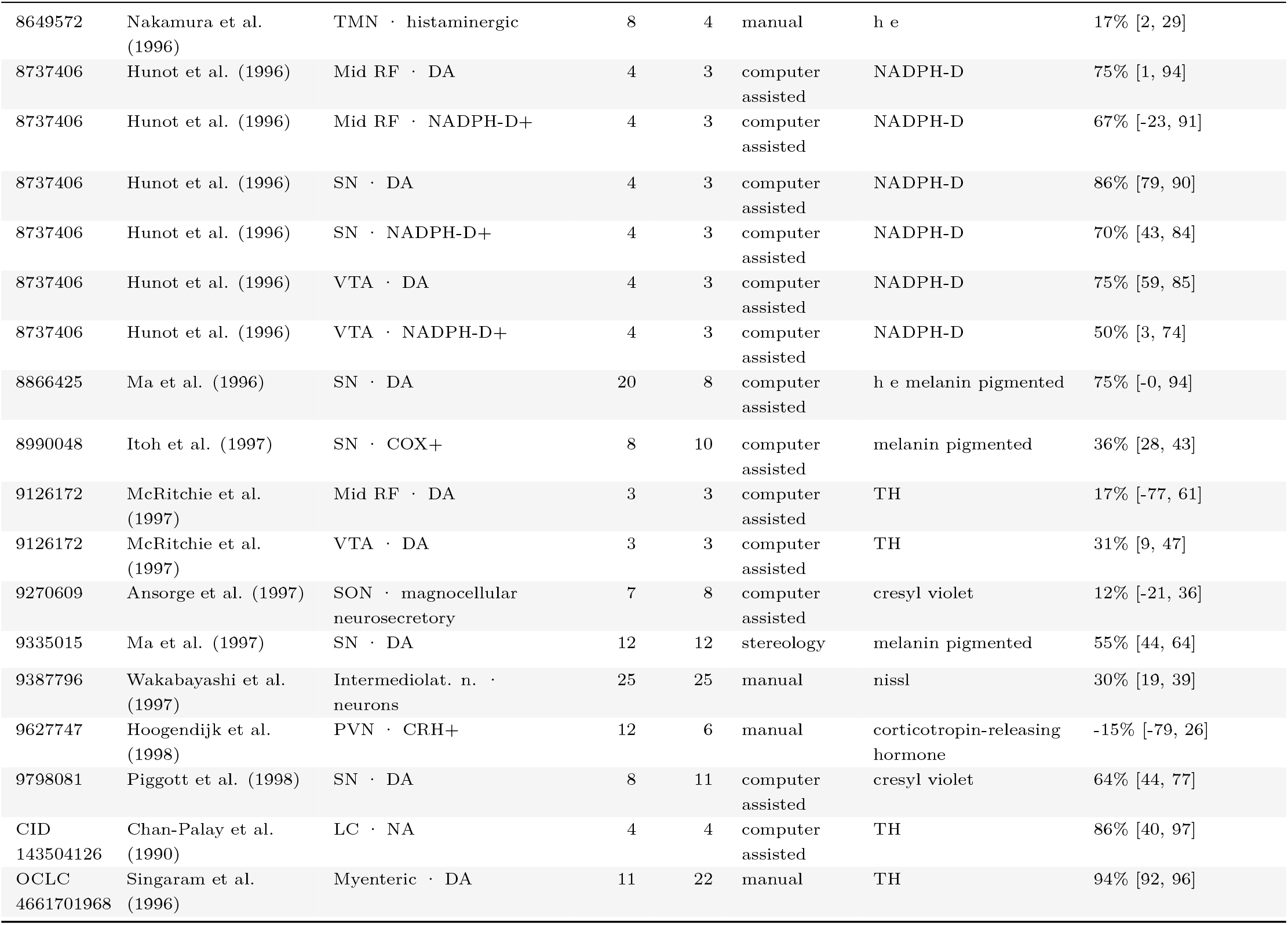
Characteristics of the 265 modelled study-population comparisons. Each row gives sample sizes, counting method, marker and reported percentage loss. Study-level intervals are descriptive delta-method 95% confidence intervals for the raw ratio of means. Pooled population estimates in the main text use Bayesian credible intervals.

**Table 8:**
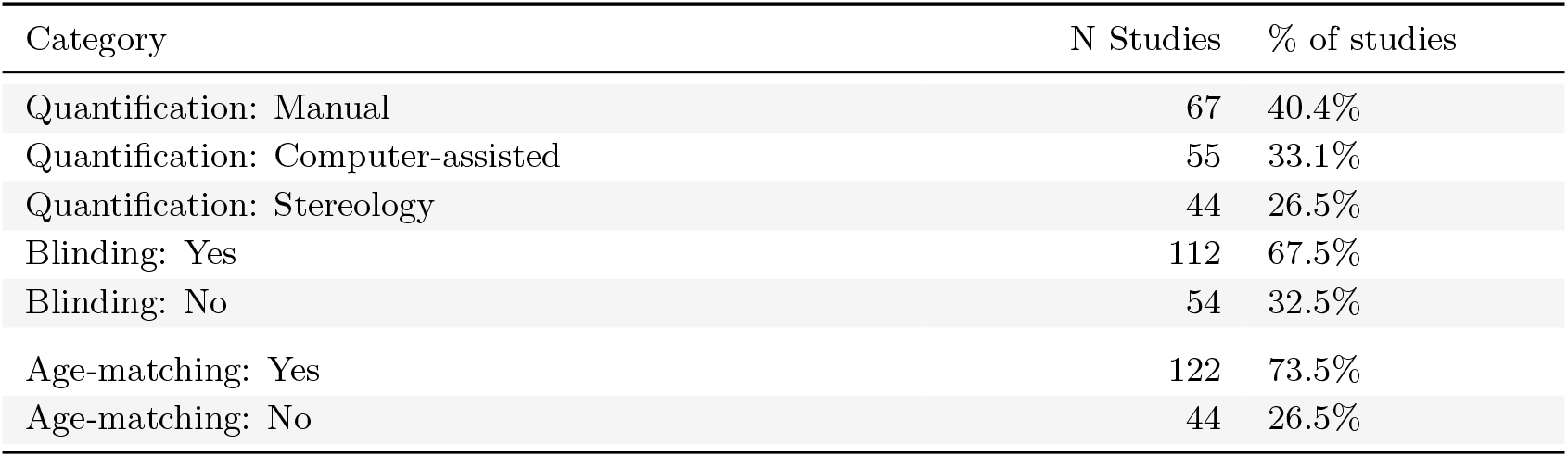
Counting methods, blinding and age matching across the 166 included studies. Each study is counted once by PMID.

**Table 9:**
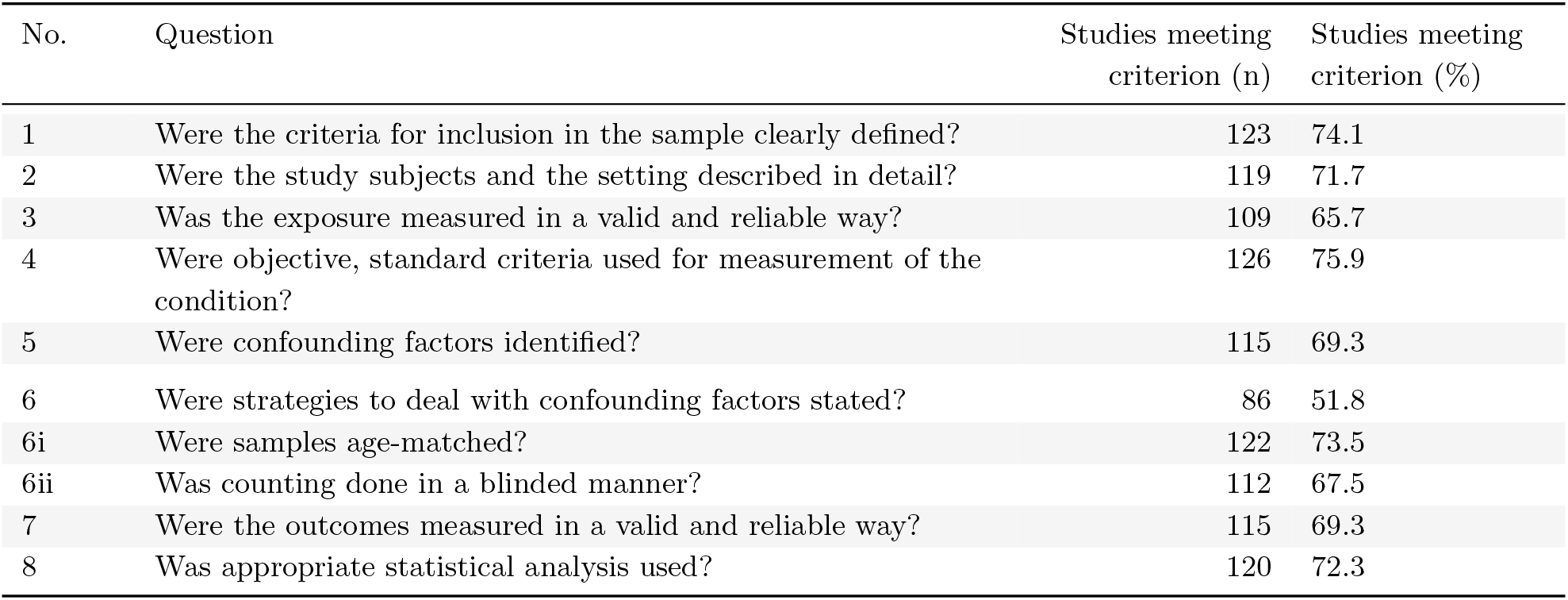
Adapted risk-of-bias checklist. Counts and percentages cover 166 studies. Eight criteria were scored Yes or No. Items 6i and 6ii describe age matching and blinded counting; the recorded rule for item 6 requires both. The composite score counts affirmative criteria.

**Table 10:**
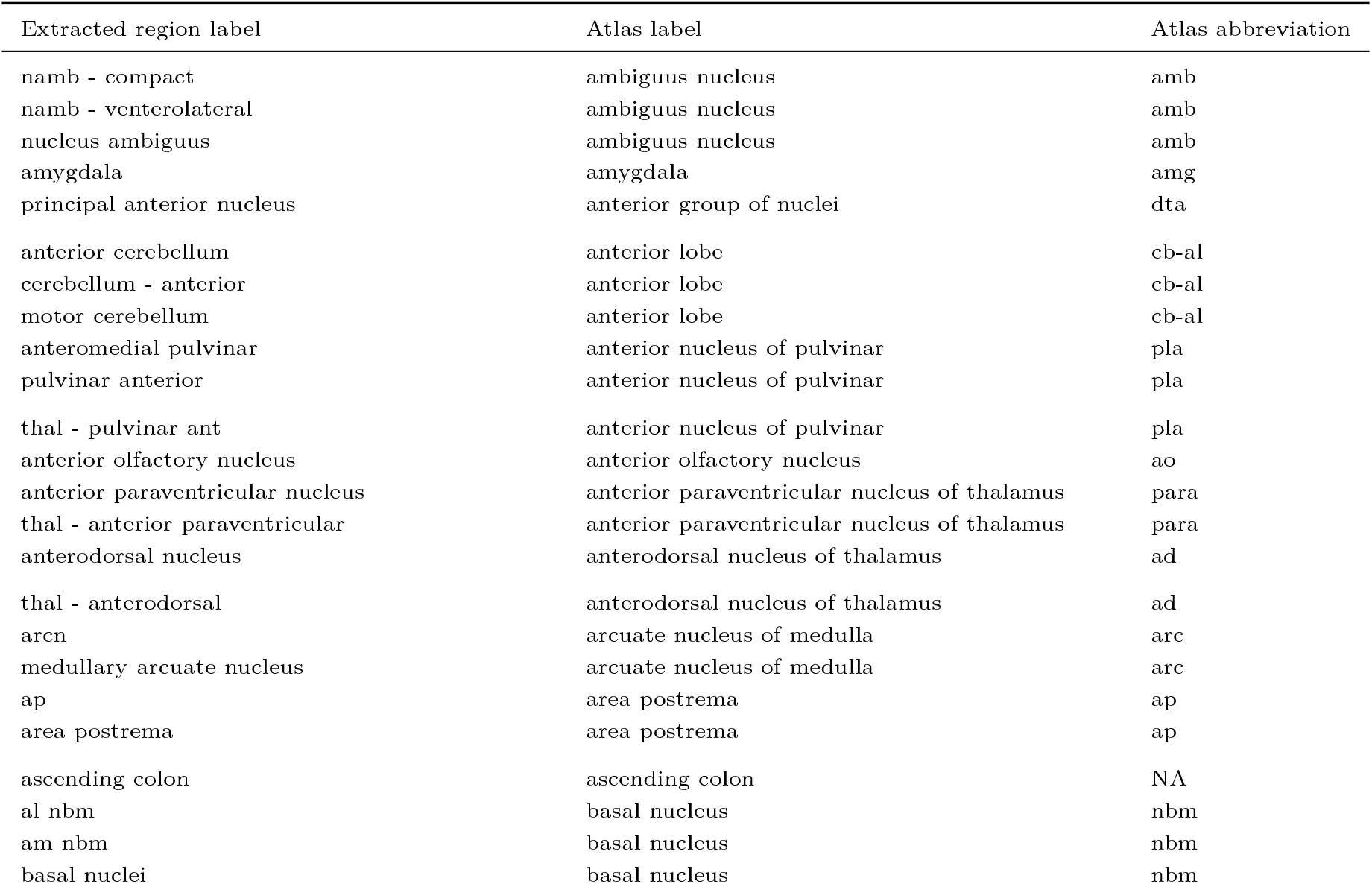
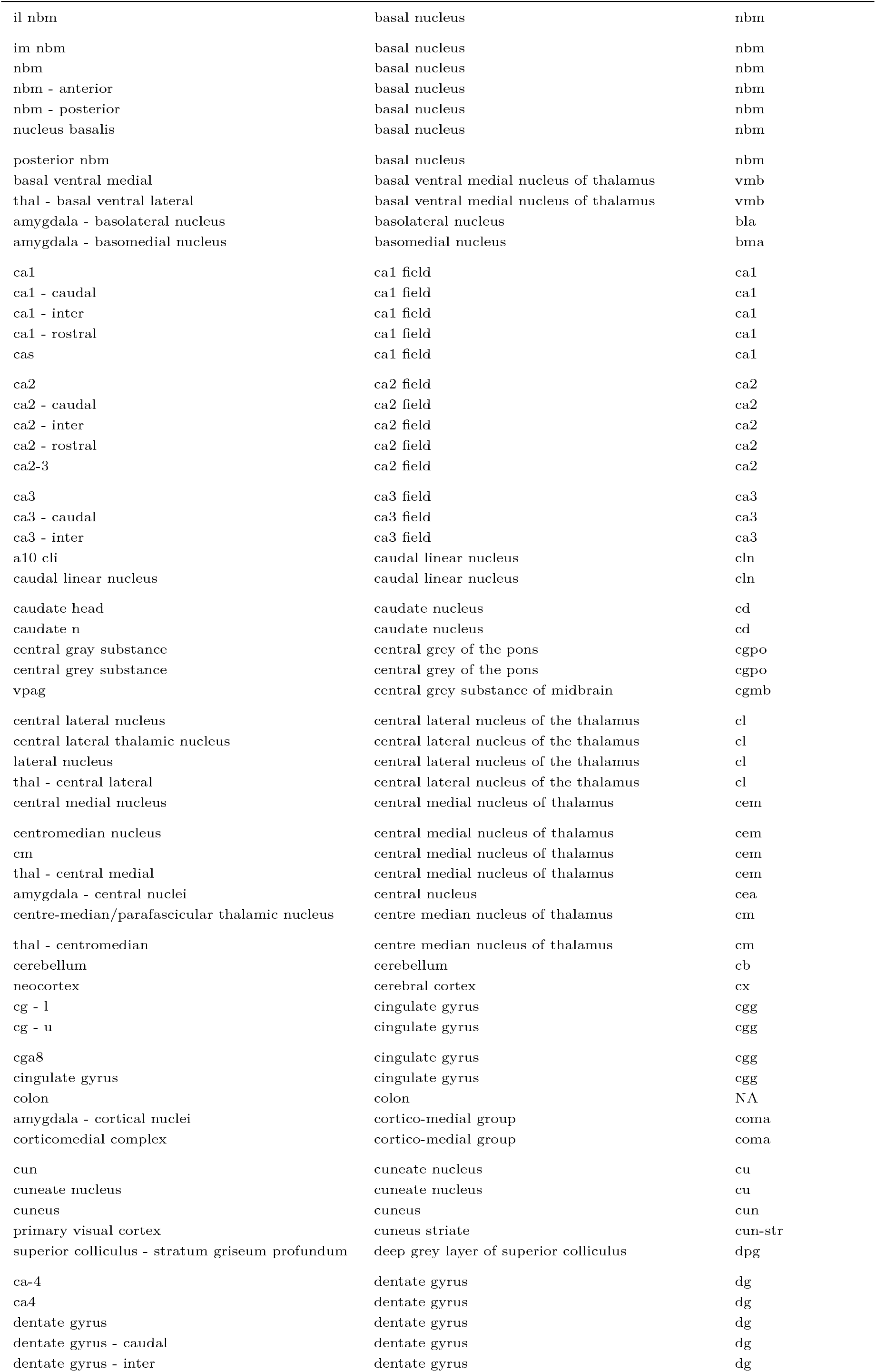
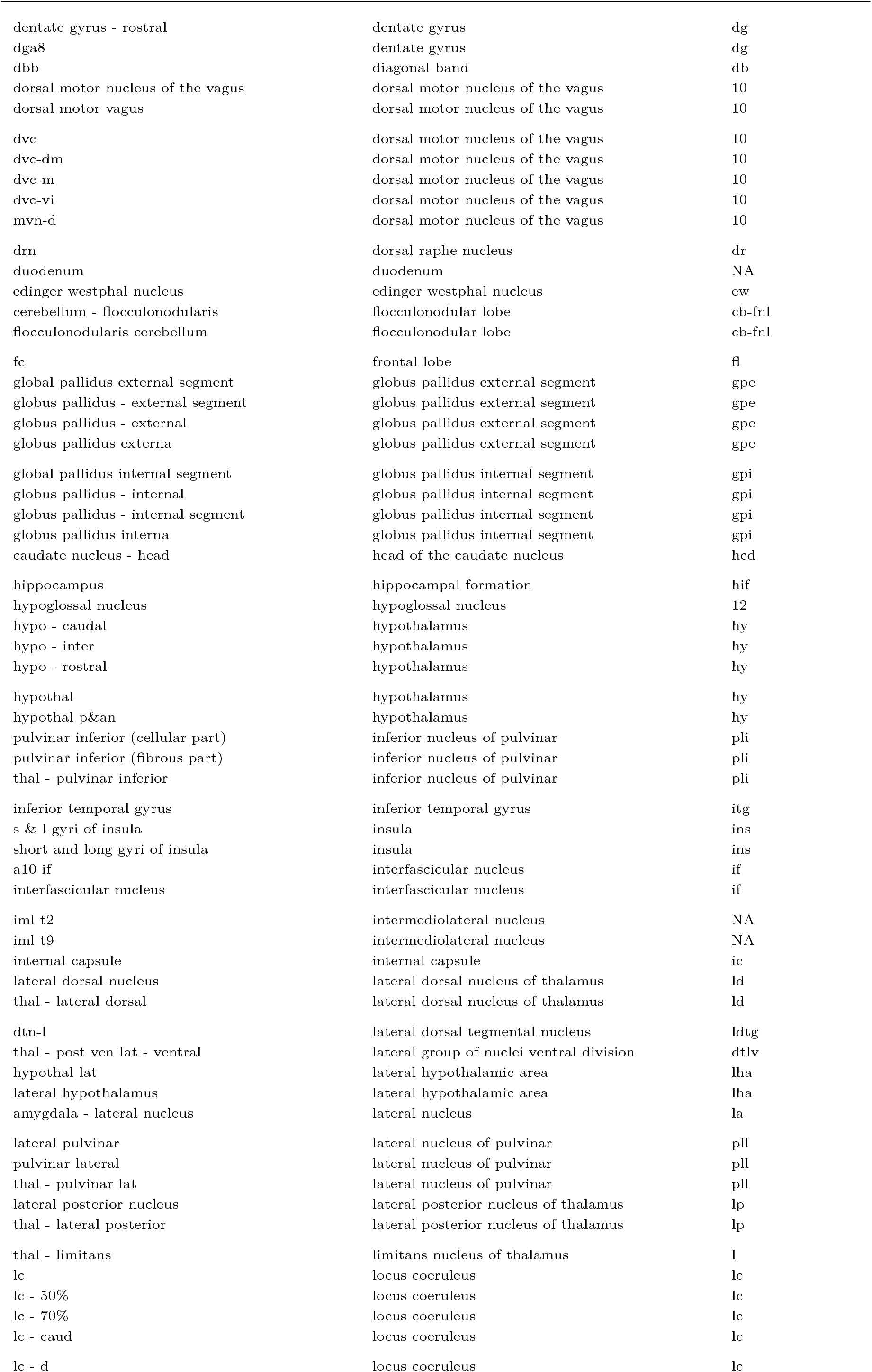
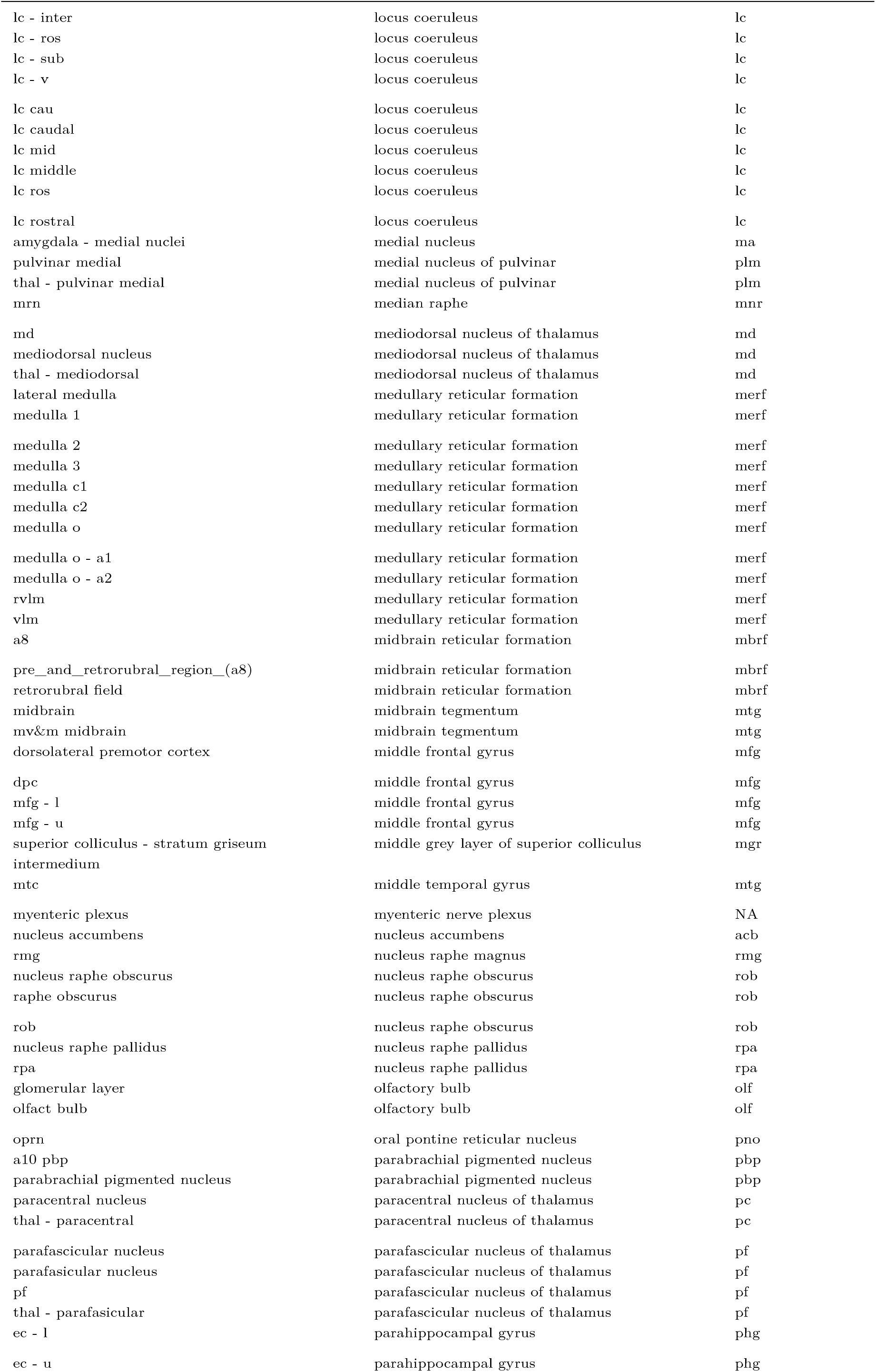
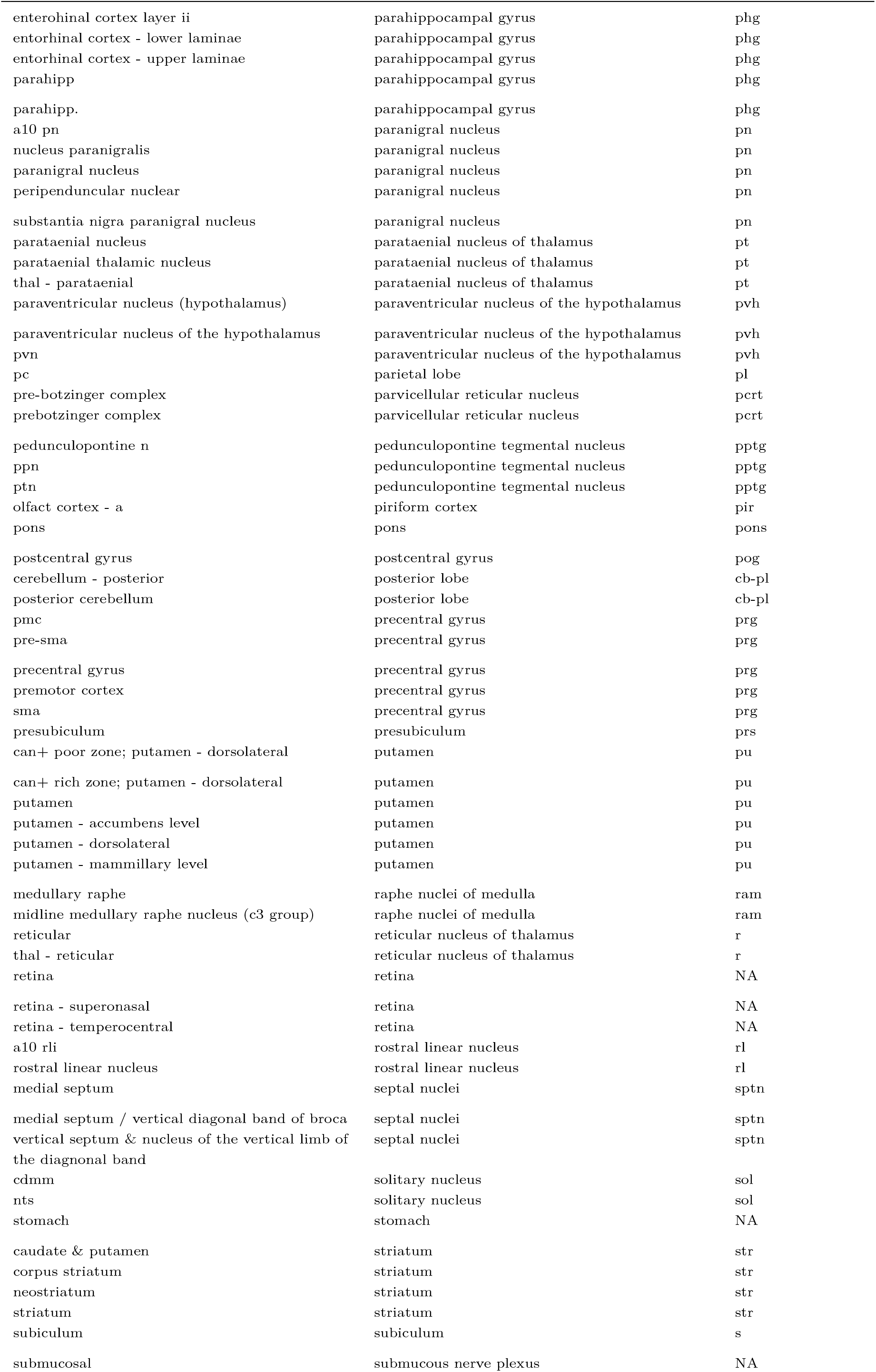
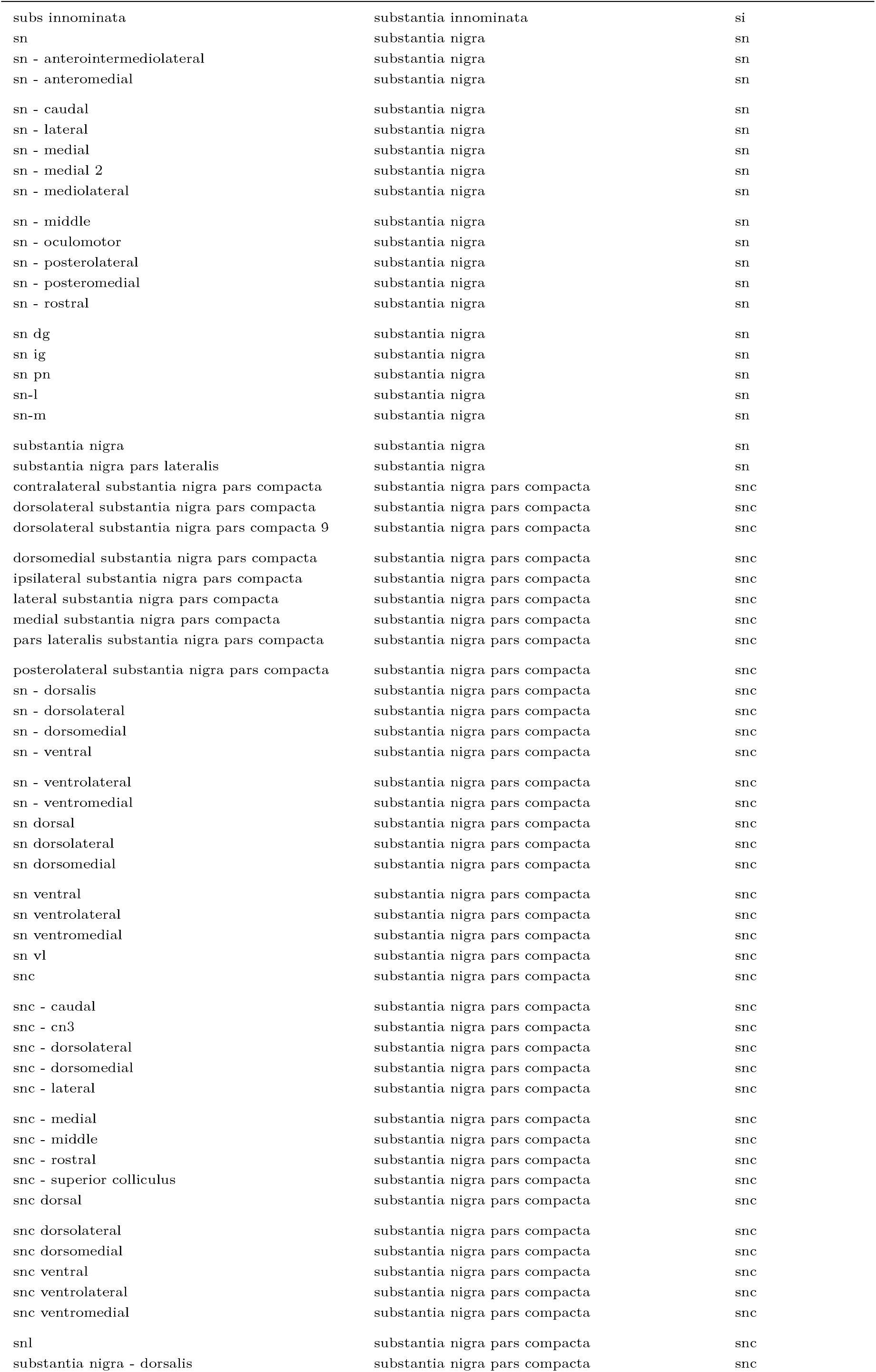
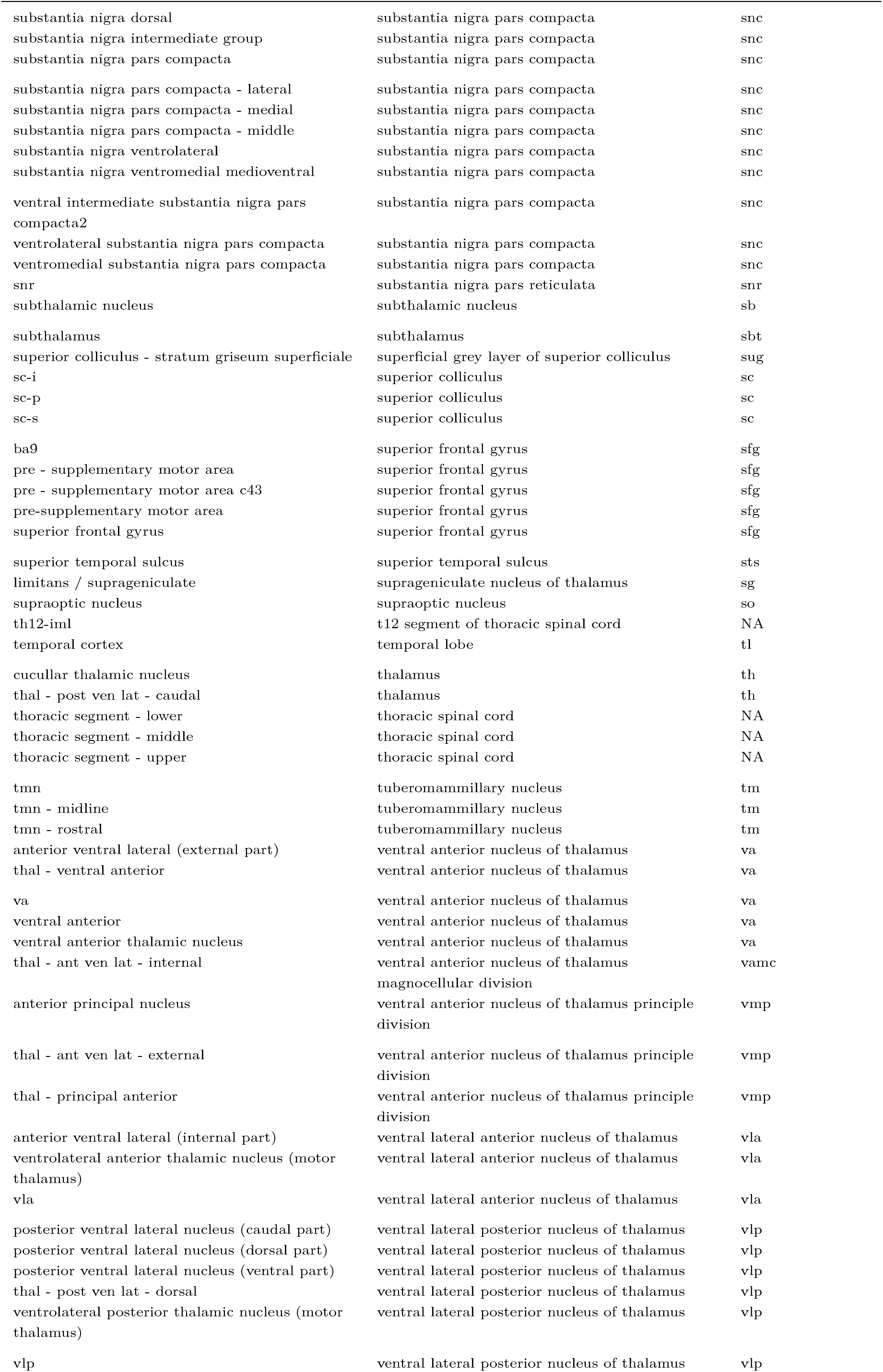
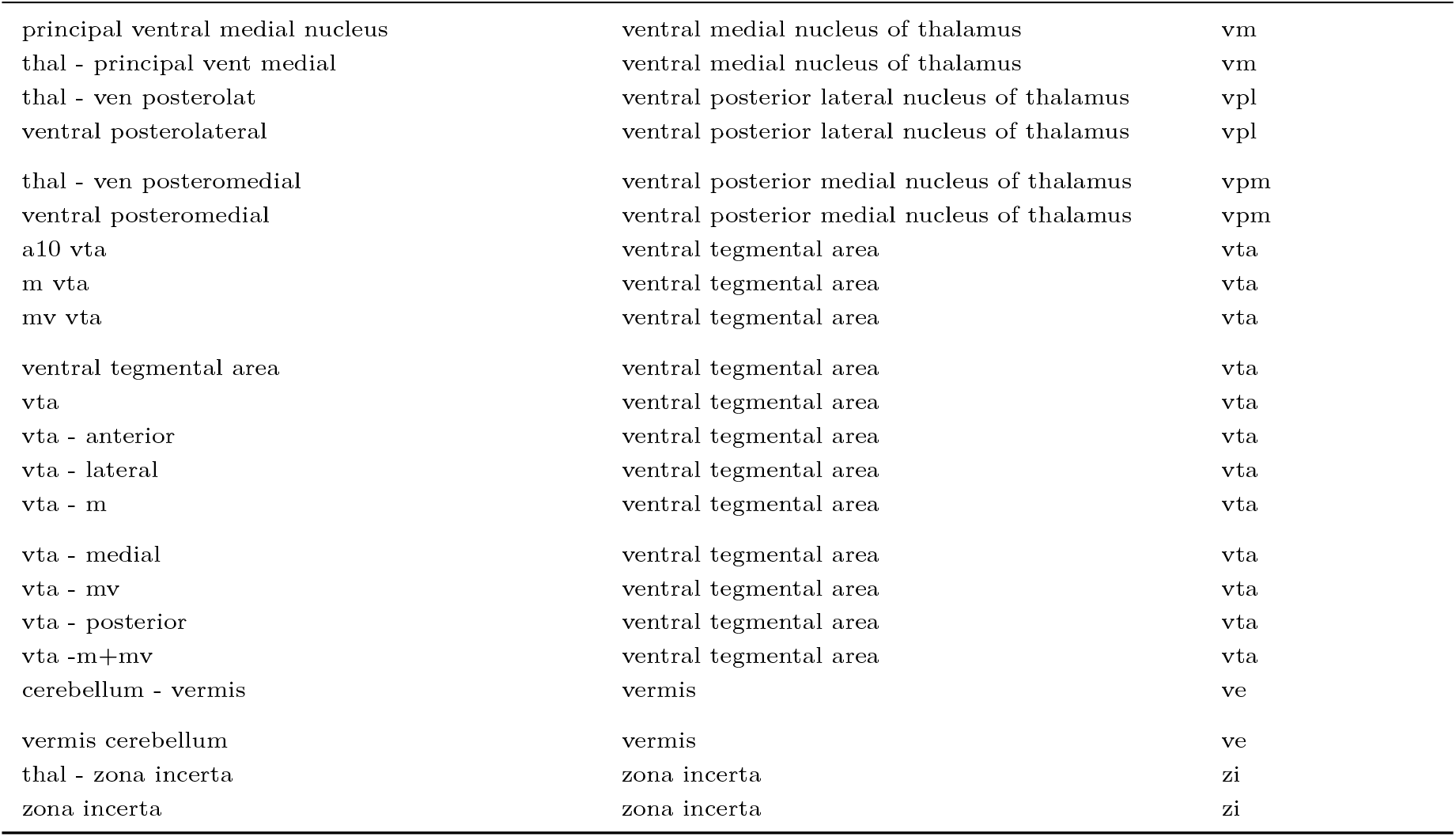
Curated mappings from extracted region labels to Allen atlas labels. Repeated ontology paths are collapsed to unique mappings. The table records the final annotations rather than a dated history of individual reviewer decisions.

**Table 11:**
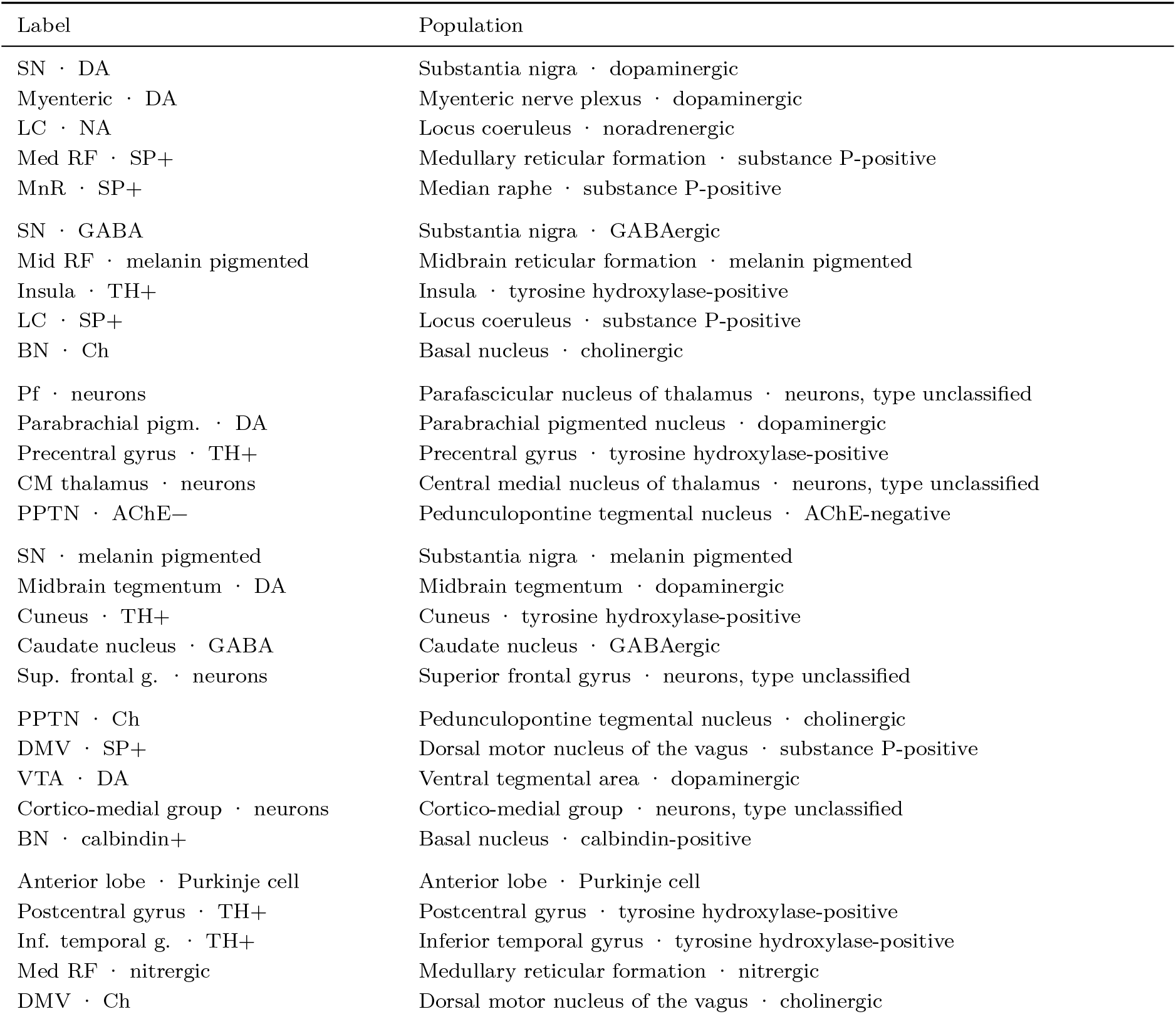
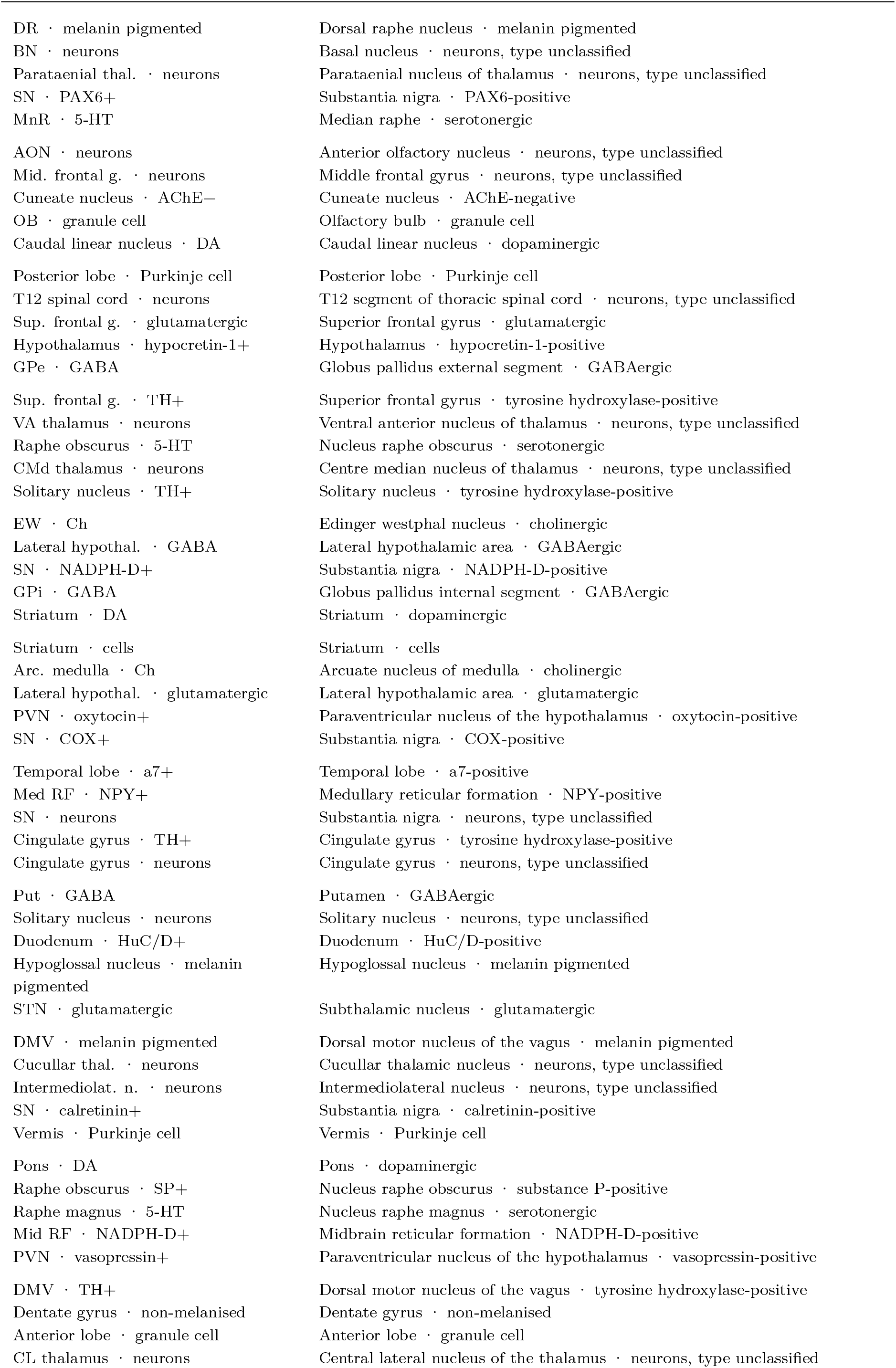
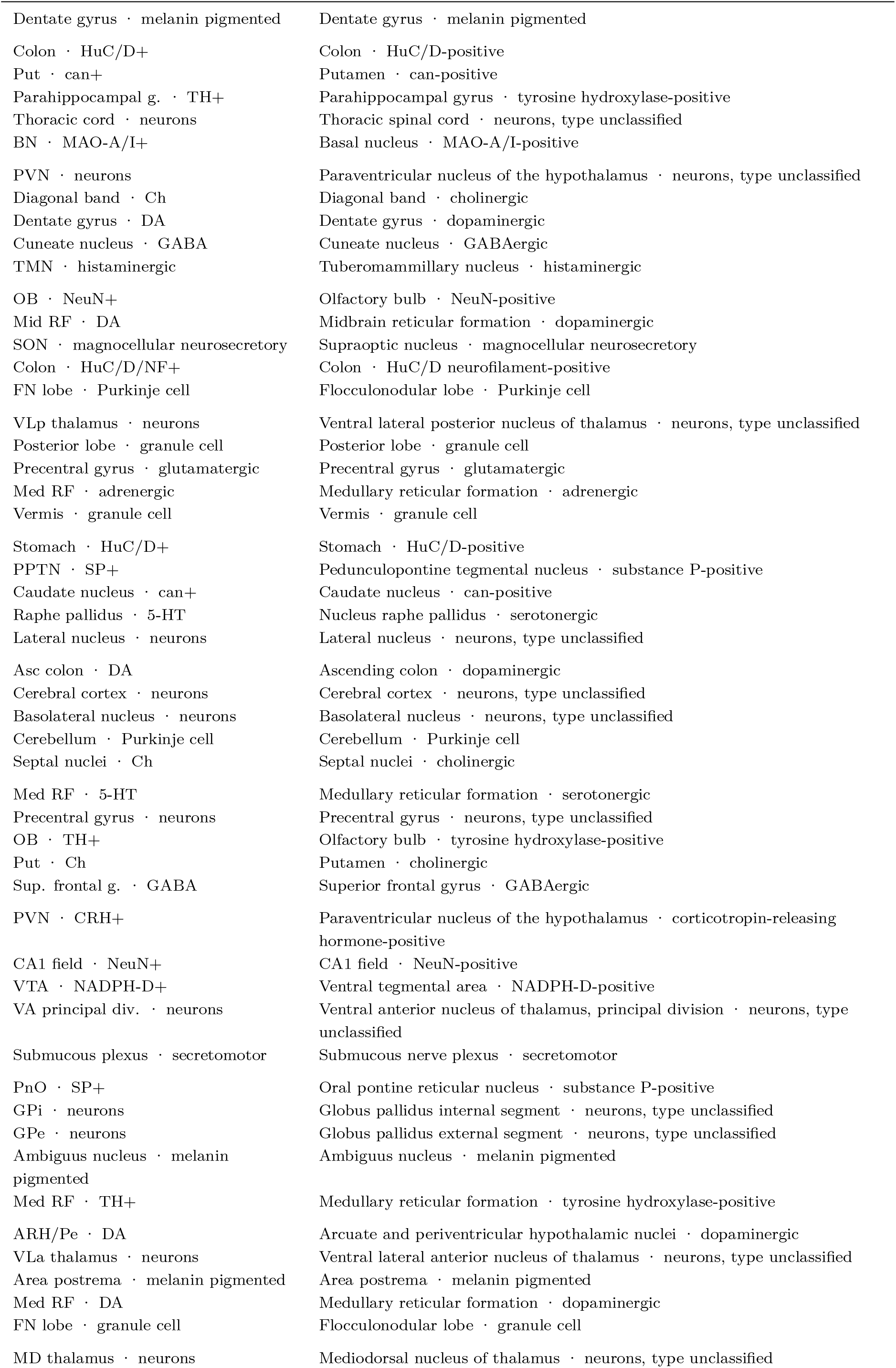
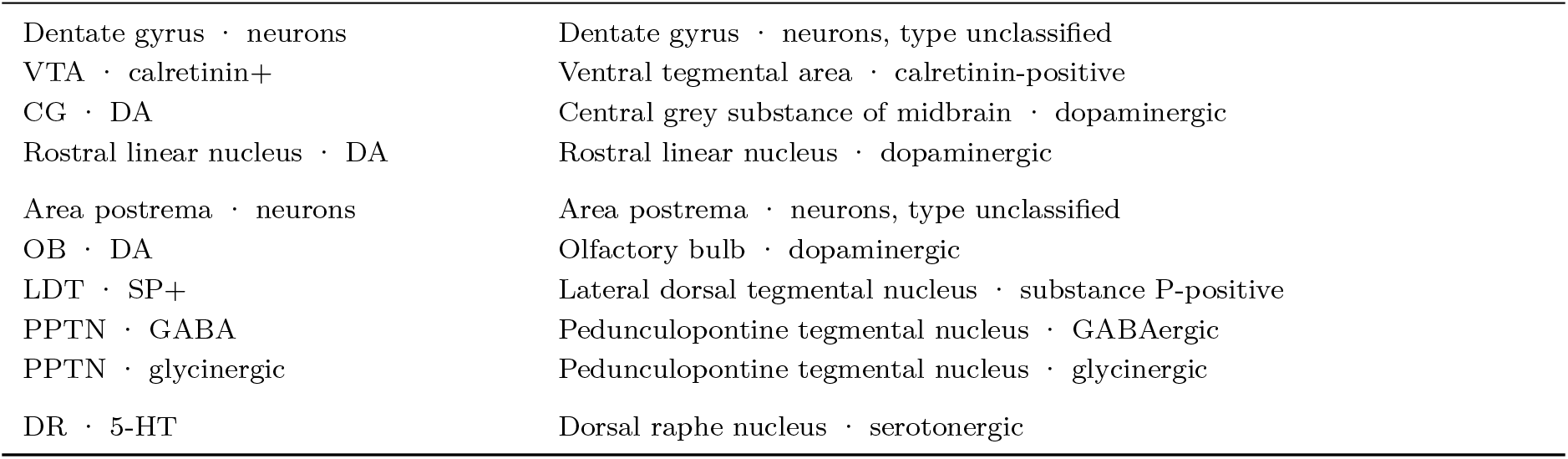
Population label key for Figures 1–3 and the analytical tables. Each compact label is paired with its full region–neuron population definition.

**Table 12:**
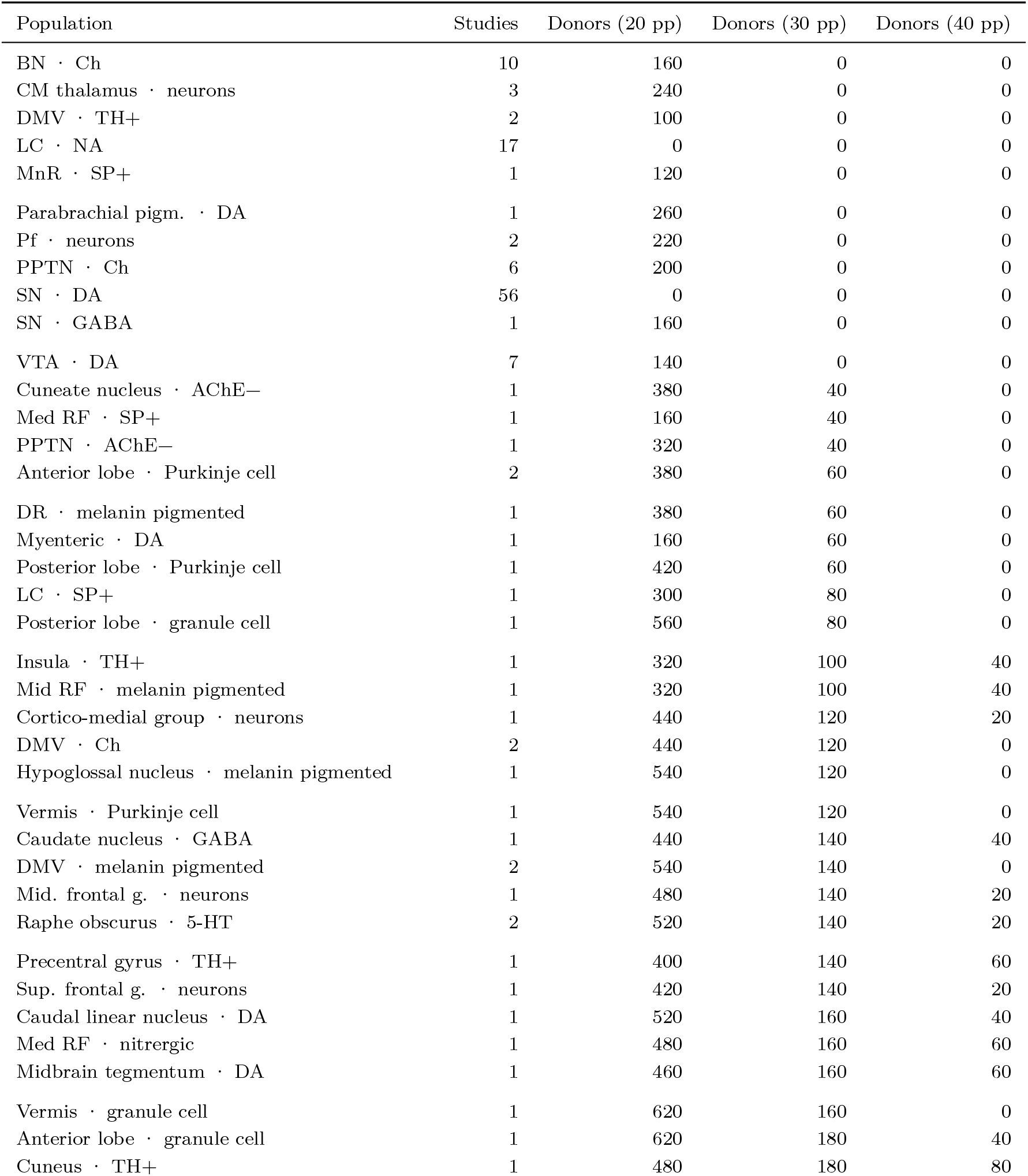
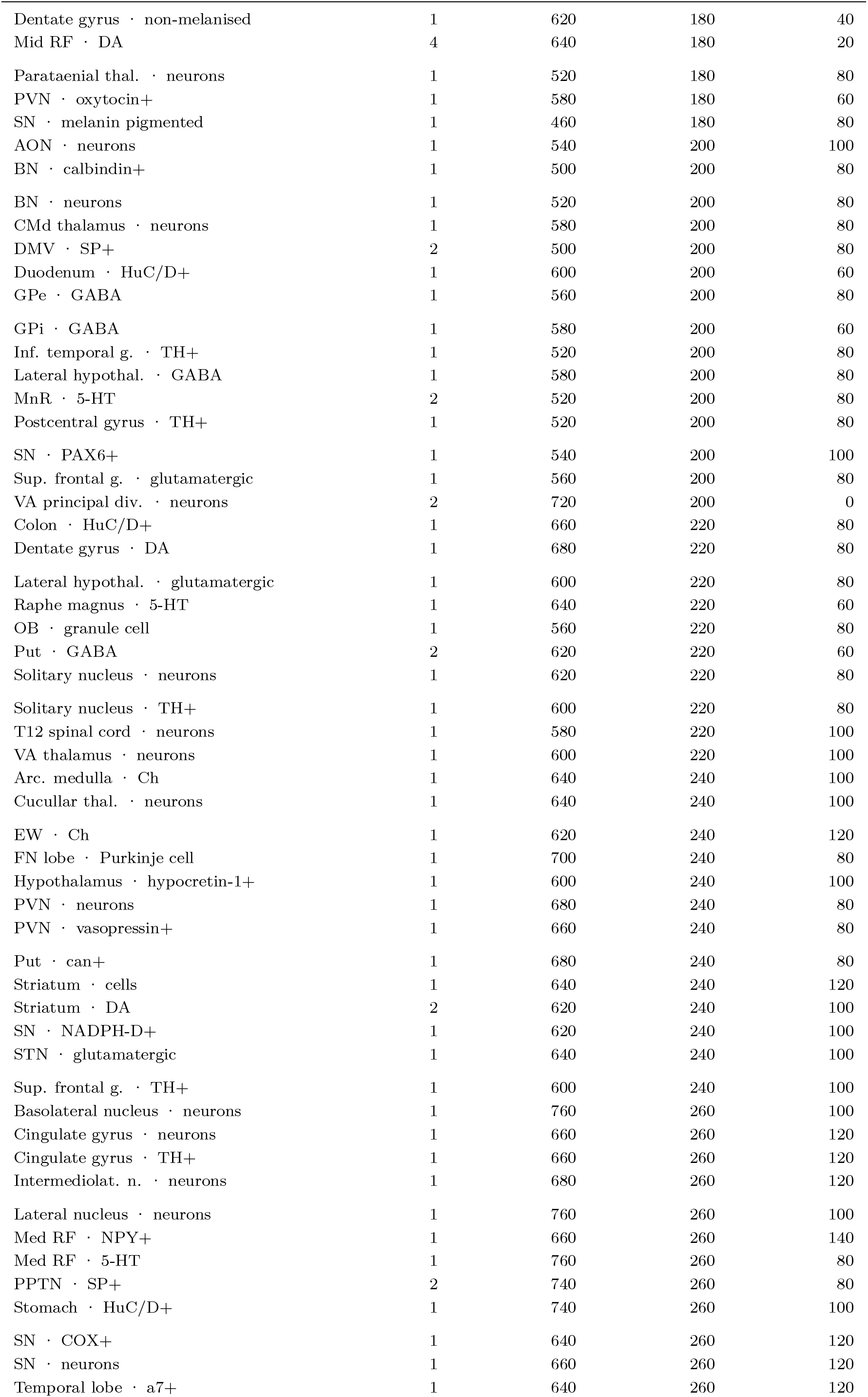
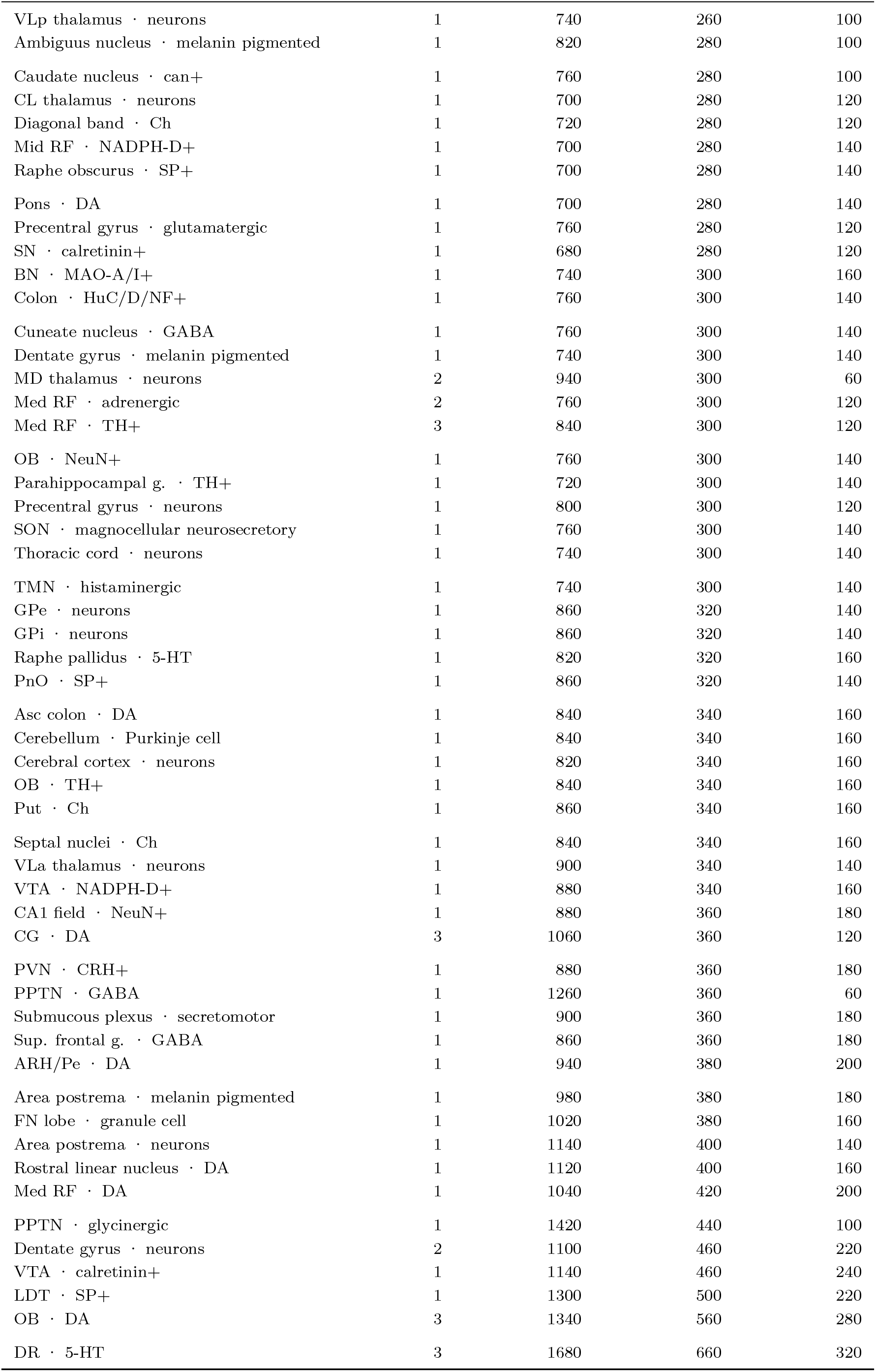
Conditional precision planning for all 146 fitted populations. The main target is a total 95% interval width of 30 percentage points. Columns show additional donors needed at widths of 20, 30 and 40 points. Zero indicates that the selected conditional precision target is already met; independent replication may still be needed. Each cohort comprises 10 PD and 10 control donors. Requirements cannot be summed across populations.

**Table 13:** Sensitivity to inferred neuron-type labels. The table compares the ten largest estimated losses in the full and reported-only datasets. Both analyses use REML fits of the combined population specification, with 95% confidence intervals for predicted population effects. The intervals are not Bayesian credible intervals and do not quantify the probability of a particular rank.

| Rank | Full model (all labels) |  |  | Reported-only model |  |  |
| --- | --- | --- | --- | --- | --- | --- |
|  | Region-Cell | k | Loss % [95% CI] | Region-Cell | k | Loss % [95% CI] |
| 1 | SN · DA | 56 | 67% [62, 71] | SN · DA | 33 | 65% [59, 70] |
| 2 | LC · NA | 17 | 64% [58, 69] | Myenteric · DA | 1 | 63% [40, 77] |
| 3 | Myenteric · DA | 1 | 63% [42, 77] | LC · NA | 10 | 63% [51, 71] |
| 4 | Med RF · SP+ | 1 | 61% [42, 74] | MnR · SP+ | 1 | 62% [47, 72] |
| 5 | MnR · SP+ | 1 | 60% [45, 71] | SN · GABA | 1 | 60% [42, 72] |
| 6 | SN · GABA | 1 | 60% [43, 72] | Mid RF · melanin pigmented | 1 | 54% [28, 71] |
| 7 | Mid RF · melanin pigmented | 1 | 52% [25, 69] | LC · SP+ | 1 | 53% [32, 68] |
| 8 | LC · SP+ | 1 | 51% [29, 67] | Insula · TH+ | 1 | 53% [26, 70] |
| 9 | Insula · TH+ | 1 | 51% [24, 69] | Midbrain tegmentum · DA | 1 | 50% [22, 68] |
| 10 | BN · Ch | 10 | 48% [34, 60] | CM thalamus · neurons | 1 | 50% [29, 64] |

**Table 14:** Association with the adapted risk-of-bias score. The study–population Bayesian model was refitted with the score as a continuous predictor. The table reports the log-ratio slope per score point, its 95% credible interval and the probability of a positive slope, followed by the predicted ratio at score 7. The fit includes 265 effects from 152 studies.

| Analysis | Estimate | 95% CrI | Posterior probability |
| --- | --- | --- | --- |
| RoB moderator: posterior slope on rob_score<br>(log-ROM per point) | 0.0639 | [0.0119, 0.1169] | P(slope > 0) = 0.99 |
| Predicted ROM at high study quality (rob_score<br>= 7) | 0.733 | [0.645, 0.830] | – |

**Table 15:** Selected effects derived from repeated grouped summaries within a study. These 19 effects from 17 studies used average means, rounded average sample sizes and root-mean-square standard deviations. This procedure does not constitute sample-size-weighted pooling of independent cohorts. PMID identifies the source study.

| PMID | Region | Neuron type | Marker |
| --- | --- | --- | --- |
| 10459912 | CG | DA | nissl nm |
| 10459912 | SN | DA | nissl nm |
| 10459912 | VTA | DA | nissl nm |
| 12419530 | SN | DA | TH |
| 15389895 | SN | DA | h e |
| 1694418 | SN | DA | TH |
| 18163454 | VTA | DA | TH |
| 18804518 | Cucullar thal. | neurons | cresyl violet |
| 1895529 | SN | DA | kluver barrera nm |
| 2257487 | SN | DA | melanin pigmented |
| 23884810 | SN | DA | melanin pigmented |
| 2549846 | SN | DA | bielchowskys stain |
| 2557795 | SN | DA | TH |
| 2709038 | SN | DA | h e melanin pigmented |
| 29917054 | SN | DA | melanin pigmented |
| 5886206 | SN | DA | gallocyenin chromealum nm |
| 7770115 | SN | DA | h e melanin pigmented |
| 8628464 | SN | DA | TH |
| 8866425 | SN | DA | h e melanin pigmented |

**Table 16:** Marker priorities for selecting measurements of the same population. Unlisted combinations receive surrogate priority. These selection rules do not establish marker specificity in every region.

| Neuron type | Marker | Priority |
| --- | --- | --- |
| DA | TH | canonical |
| DA | melanin pigmented th | canonical |
| DA | melanin pigmented nurr1 | canonical |
| DA | bfgf th | canonical |
| DA | melanin pigmented | canonical |
| DA | h e melanin pigmented | canonical |
| DA | cresyl violet melanin pigmented | canonical |
| DA | RET | canonical |
| DA | scRNA | canonical |
| DA | MAO-B | surrogate |
| DA | NADPH-D | surrogate |
| DA | calbindin | surrogate |
| DA | tau | surrogate |
| DA | p-RPS6 | surrogate |
| DA | bielchowskys stain | surrogate |
| DA | cresyl violet | surrogate |
| DA | cresyl violet nm | surrogate |
| DA | cresyl violet luxol fast blue nm | surrogate |
| DA | nissl nm | surrogate |
| DA | melanin pigmented nissl | surrogate |
| DA | h e | surrogate |
| DA | kluver barrera nm | surrogate |
| DA | gallocyanin chromealum nm | surrogate |
| DA | fm and neutral red | surrogate |
| DA | azure b nm | surrogate |
| DA | azure b | surrogate |
| DA |  | surrogate |
| NA | DBH | canonical |
| NA | TH | canonical |
| NA | melanin pigmented | canonical |
| NA | h e melanin pigmented | canonical |
| NA | melanin pigmented nissl | surrogate |
| NA | bodian h e melanin pigmented nissl | surrogate |
| NA | cresyl violet | surrogate |
| NA | h e | surrogate |
| NA | golgi nissl | surrogate |
| NA | azure b nm | surrogate |
| Ch | ChAT | canonical |
| Ch | NGFR | canonical |
| Ch | ACh | canonical |
| Ch | ach nissl | canonical |
| Ch | cresyl violet | surrogate |
| Ch | cresyl violet tau | surrogate |
| Ch | bodian cresyl violet | surrogate |
| Ch | h e | surrogate |
| Ch | h e melanin pigmented | surrogate |
| 5-HT | TrOH | canonical |
| 5-HT | PH8 | canonical |
| 5-HT | TH | surrogate |
| adrenergic | PNMT | canonical |
| adrenergic | melanin pigmented | surrogate |
| GABA | parvalbumin | canonical |
| GABA | calretinin | canonical |
| GABA | cabp i | canonical |

**Table 17:** Estimated neuronal loss by counting method after adjustment in the multilevel model. Intervals are 95% credible intervals. Differences in populations studied, publication era and study quality limit causal interpretation.

| Method | Estimate |
| --- | --- |
| Manual | 38.7% (29.5–47.0%) |
| Computer-assisted | 27.6% (15.6–38.3%) |
| Stereology | 30.7% (18.8–41.2%) |

| Neuron type | Marker | Priority |
| --- | --- | --- |
| GABA | NADPH-D | canonical |
| GABA | NeuN | surrogate |
| GABA | SMI-32 | surrogate |
| GABA | MCH | surrogate |
| GABA | NK1R | surrogate |
| GABA | cresyl violet | surrogate |
| GABA | h e | surrogate |
| GABA | nissl | surrogate |
| GABA | scRNA | canonical |
| glutamatergic | HCRT | canonical |
| glutamatergic | cresyl violet | surrogate |
| glutamatergic | h e | surrogate |
| neurons | NeuN | canonical |
| neurons | nissl | canonical |
| neurons | cresyl violet | canonical |
| neurons | cresyl violet nissl | canonical |
| neurons | giemsa | surrogate |
| neurons | giemsa h e | surrogate |
| neurons | h e | surrogate |
| neurons | h e melanin pigmented | surrogate |
| neurons | oxytocin | surrogate |
| neurons | vasopressin | surrogate |
| melanin pigmented | melanin pigmented | canonical |
| melanin pigmented | h e melanin pigmented | canonical |
| melanin pigmented | melanin pigmented nissl | canonical |
| melanin pigmented | nissl nm | canonical |
| melanin pigmented | h e | surrogate |
| melanin pigmented |  | surrogate |

## Notes

### Competing Interest Statement

The authors have declared no competing interest.

### Summary of Updates

Title, abstract and main text revised; primary analysis updated to Bayesian multilevel meta-analysis following data corrections; power analysis replaced with conditional precision planning; figures redesigned to compare proposed vulnerability mechanisms; Methods, Discussion and supplementary materials updated.

